# Double crossed? Structural and computational studies of an unusual crosslinked heme in *Methylococcus capsulatus* cytochrome P460

**DOI:** 10.1101/2024.12.21.628999

**Authors:** Hans E. Pfalzgraf, Aditya G. Rao, Kakali Sen, Hannah R. Adams, Marcus Edwards, You Lu, Chin Yong, Sofia Jaho, Takehiko Tosha, Hiroshi Sugimoto, Sam Horrell, James Beilsten-Edmands, Robin L. Owen, Colin R. Andrew, Jonathan A. R. Worrall, Ivo Tews, Adrian J. Mulholland, Michael A. Hough, Thomas W. Keal

## Abstract

Cytochromes P460 oxidise hydroxylamine within the nitrogen cycle and contain as their active site an unusual catalytic c-type heme where the porphyrin is cross-linked to the protein via a lysine residue in addition to the canonical cross links from cysteine residues. Understanding how enzymes containing P460 heme oxidise hydroxylamine into either nitrous oxide or nitric oxide has implications for climate change. Interestingly the P460 containing hydroxylamine oxidoreductase utilises a tyrosine cross link to heme and performs similar chemistry. Previous crystal structures of cytochrome P460 from *Nitrosomonas europaea* (NeP460) clearly show the existence of a single crosslink between the Nz atom of lysine and the heme porphyrin with mutagenesis studies indicating roles for the crosslink in positioning a proton transfer residue and/or influencing the distortion of the heme. Here we describe the evidence for a novel double cross link between lysine and heme in the cytochrome P460 from *Methylococcus capsulatus* (Bath). In order to understand the complexities of this enzyme system we applied high resolution structural biology approaches at synchrotron and XFEL sources paired with crystal spectroscopies. Linked to this we carried out QM/MM simulations that enabled the prediction of electronic absorption spectra providing a crucial validation to linking simulations and experimental structures. Our work demonstrates the feasibility of a double crosslink in McP460 and provides an opportunity to investigate how simulations can interact with experimental structures.

## Introduction

The oxidation of hydroxylamine is a key part of the process of nitrification, whereby fixed nitrogen is converted to biologically available forms such as nitrate, occurring in a range of bacteria. In this process, the reactive intermediate hydroxylamine is formed through the oxidation of ammonia by ammonia monooxygenases before being converted to a series of nitrogen oxides, including the damaging greenhouse gas nitrous oxide. Two enzymes that have been identified as carrying out the challenging hydroxylamine oxidation reaction each have unusual cross linked heme active sites which share a characteristic absorption maximum of 460 nm in the ferrous state. Hydroxylamine oxidoreductases (HAO) use a P460 heme with a crosslink between a tyrosine residue and the heme. In contrast, cytochromes P460 have a crosslink between a lysine and the heme. The P460 heme is thus characterised by a protein to porphyrin crosslink in addition to the canonical cross link of two cysteine residues that occurs in all c-type hemes. HAO and P460 differ in their reaction mechanism and product despite acting upon the same substrate. P460s consist of a homodimer with one heme P460 present per monomer. in contrast, HAO is an octaheme protein with seven electron transfer hemes and a single P460 heme. For this reason, cytochromes P460 are far more tractable for experimental and computational study. The properties and molecular structure of several catalytically active cytochrome P460s have been characterised, particularly those from the ammonia oxidising *Nitrosomonas europaea* (NeP460) and methane oxidising *Methylococcus capsulatus* (Bath) bacteria (McP460). The structures reveal rather different active site pockets but in both cases a solvent filled, highly polar active site pocket is present positioning residues appropriately for electron and proton transfer. McP460 lacks the C-terminal helix seen in NeP460 but harbours three arginine residues in the active site. Crystal structures have been determined for NeP460^1^ and its crosslink-deficient mutant^2^, for McP460^3^ and for an inactive cytochrome P460 from *Nitrosomonas* sp. AL212^4^ (NsALP460) and its mutants including an active one^5^.

All published crystal structures of these enzymes have been determined at 100 K using synchrotron radiation. While the structures are highly informative, the possibility of electronic states changes around the group as a result of radiation damage is highly likely^6,7,8^. Spectroscopic data from crystals has revealed that reduction typically takes place at doses far lower than those required for structure determination using conventional methods. Moreover, cryo- cooling of protein crystals can suppress functionally relevant dynamics and/or lead to the observation of structural artefacts.

Extensive work has been carried out by Lancaster and co-workers on cytochromes P460, identifying the role of the active site pocket structure and the crosslink in positioning the heme and crucial residues involved in proton transfer, in influencing the heme geometry and in the origins of the unusual spectroscopic properties of cytochromes P460. Site-directed mutagenesis of NsALP460 to introduce a glutamate residue (Glu131) mimicking the active site pocket of NeP460 (Glu97) led to gain of hydroxylamine oxidation activity.^5^ This activity can be removed by knocking out the crosslinking lysine,^9^ leading to a mutant enzyme with standard spectroscopic properties for c-type heme enzymes This identified the glutamate in that position as being important for proton transfer within the catalytic mechanism with the proposal that one role of the protein crosslink to heme was to correctly position this glutamate for proton transfer.

Molecular simulation studies of cytochrome P460 enzymes have been limited to date. These were mainly restricted to cluster calculations using density functional theory (DFT) and omit the role of the protein environment. The absorption spectra of the crosslinked heme moiety and inner-sphere ligands in NeP460 have been simulated with time-dependent density functional theory (TD-DFT). The spectroscopic calculations revealed that the experimentally observed spectral red shift in NeP460 can be attributed to geometric distortions within heme, in particular, the deviation of the macrocycle out of planarity^4^.

In this work we describe a structure of native McP460 obtained at room temperature using serial femtosecond crystallography (SFX) at an X-ray free electron laser (XFEL) to exclude any effects of radiation damage to the sample that could affect the structure or electronic state. Importantly, to provide the best model as inputs to QM/MM molecular simulations it is critical to have a true ground state structure that has not been modified by the data collection technique used to determine that structure. This could not be achieved using synchrotron radiation and required the unique properties of an XFEL.

We also describe structures validated by absorption spectroscopy carried out on crystals to explicitly identify the oxidation state and/or ligand state of the crystals. This enables a structure of the Fe(II) (ferrous) enzyme and the iron-bound ligand of the Fe(III) (ferric) structure to be assigned. Crucially, we use this data to carry out QM/MM molecular simulations of cytochrome P460 and to validate these by prediction of spectroscopic data that can be readily compared to experimental results. Our data suggest the presence of an unusual double cross link between the heme and active site lysine residue in contrast to the single cross link to lysine that is present in NeP460. The possible chemical natures of the crosslinks observed in the experimental structures were suggested as shown in Scheme 1: A) a single covalent link between Lys78-N and Heme-CHA (single crosslink, SC) , B) an additional covalent link between Lys78-CD and Heme- C2A (double crosslink, DC) and C) a double crosslinked species with a double bond in the Lysine side chain between Lys78-CD and Lys78-CE (double crosslink with unsaturated Lys side chain, DCu). These three alternatives were modelled into the active site of the crystal structures and structural and spectral determination were carried out via QM/MM simulations to identify the nature crosslink species present in the oxidised and reduced X-ray structures. Scheme 2 provides a chart of the experimental structures that were used for simulations and the oxidation state of Fe and the nature of crosslinks explored via simulations. The study provides an exemplar of close interplay between molecular simulation and experimental studies, used here to understand structure of the covalently linked Lys-heme unit in this challenging enzyme system.

**Scheme 1.**
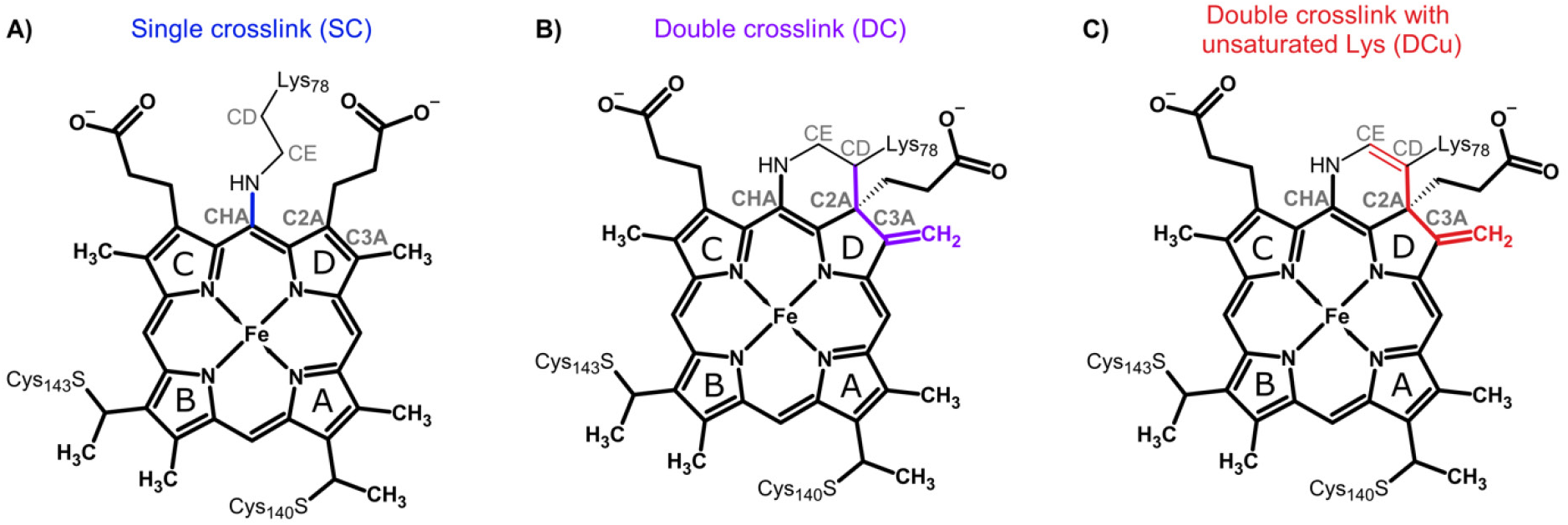
Schematic representation the heme unit showing the different type of crosslinks considered for McP460. Atom names are indicated in grey. Pyrrole ring names are labelled with black letters. Heme atoms are shown in bold. The bonds that undergo modification due to the crosslinks are coloured blue: SC, violet: DC, and red: DCu.

**Scheme 2.**
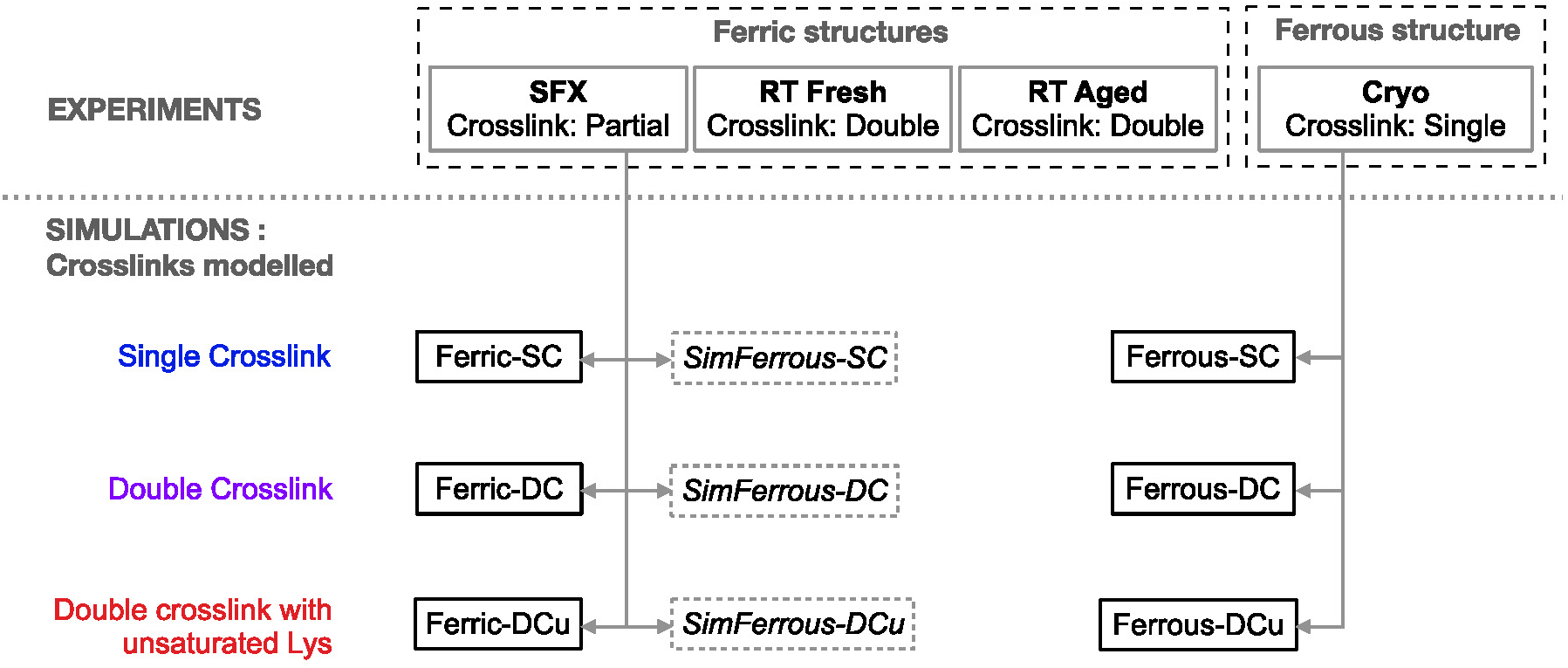
A workflow of the structures determined experimentally and those that were used as the basis for simulations. The **Ferric SFX** structure was simulated in its Ferric state for the single crosslink, double crosslink and double crosslink with unsaturated Lysine side chain models, labelled Ferric-SC/DC/DCu respectively. The **Ferrous Cryo** structure was also used as the basis for simulation of all crosslink alternatives, labelled Ferrous-SC/DC/DCu. Finally, for validation purposes ferrous models were also generated from the **Ferric SFX** structure, labelled *SimFerrous-SC/DC/DCu*.

## Materials and methods

### Activity Assays and N2O and NO2^-^ Quantification

Recombinant McP460 was expressed and purified as described previously (Adams et al 2019). Activity Assays were carried out as previously described^3^ Production of N_2_O was determined by gas chromatography-mass spectrometry using a Shimidzu GC-2014 equipped with an Porapak- Q- column (80/100 mesh 1 m × 2.3 mm I.D). All reactions were prepared in 10 mL septum-sealed headspace vials. For anaerobic reactions, samples were prepared in a glovebox. Turnover conditions were achieved with 5 µM cyt P460, 6µM PMS, 2 mM hexaamineruthenium(III) chloride ([Ru(NH_3_)_6_]Cl_3_), and 0 µM, 100 µM, 250µM, 500 µM, or 1000 µM NH_2_OH in 500 µL of buffer total. Reactions were initiated by the addition of NH_2_OH, the vial was quickly capped and inverted three times. The headspace was sampled after reactions were allowed to age overnight. N_2_O production was quantified by integrating the peak corresponding to N_2_O (retention time = 2.13 min). Nitrite production was quantified using a Griess diazotization assay. Samples were prepared aerobically with 5 µM cyt P460, 6 µM PMS, 2 mM ruthenium hexachloride, and 0, 100µM, 250 µM, 500 µM, or 1000 µM NH_2_OH in 500 µL of buffer total.

### Crystallographic data collection and processing

#### Serial femtosecond crystallography (SFX)

Micro-crystals were grown in batch using a modification of the crystallization conditions previously used for growing large single crystals, 3.8-4.0 M ammonium sulfate, 0.1 M Tris pH 8 in a 1:1 ratio of protein:crystallisation condition and 1 µl of microseeds. The final protein concentration in the batches ranged from 20 to 30 mg ml^-1^. Crystals group in 1-2 days at 18 °C to dimensions of 44 × 41 × 18 (± 9 × 11 × 6, n = 20) (Fig. S1). To remove the precipitate, batches were centrifuged at 2300 *g* for 1 min, leading to sedimentation of the crystals as a dark green band surrounded by precipitate. The precipitate was carefully removed with a pipette and the crystals were resuspended in 3.8 M ammonium sulfate, 0.1 M Tris pH 8. This process was repeated 1–2 times until most of the precipitate had been removed, as judged by inspection under a standard microscope. To generate the microseeds, crystals grown by vapour diffusion as previously described^3^ were disrupted using seed beads (Hampton Research). Crystals were transported at ambient temperature. A slurry of batch micro crystals (300 µL total) was loaded onto two silicon fixed target chips with 9-10 μm apertures ^10^ for SFX data collection at the SACLA XFEL, BL2 (Harima Campus, Japan) using a Rayonix MX300-HS detector. The experimental set up has been previously described in detail.^11^ The X-ray pulse length was 10 fs and the repetition rate was 30 Hz. The pulse energy was 360 microjoules and the photon energy 11 keV (FWHM 25.6 eV). The beam size was 1.65 μm horizontally and 1.29 μm vertically. Data were collected at 28° C. SFX data were processed using xia2.ssx^12^ with a total of 29,559 crystals included in the final dataset. A resolution cut-off was imposed at the point where the correlation coefficient fell monotonically below 0.3. CCP4i2^13^ was used to generate a free R set and REFMAC5^14^ and Coot 0.9^15^ were used to iteratively refine and build the structures based on the previously published 100 K structure.^3^ The program qFit 3.2.2^16^ was used to suggest alternate conformations based on a composite omit map generated with Phenix.^17^

#### Room-temperature synchrotron crystallography (RT fresh and aged)

*In situ* room-temperature crystallography was used to obtain the structure of McP460 with a single conformation of the crosslinking Lys78 (double-crosslinked). A Mosquito® LCP liquid-handling robot (SPT Labtech) was used to set In Situ-1™ crystallization plates (MiTeGen). It dispensed 100 nL of McP460 at 7 mg/mL (based on a molecular weight of 15.6 kDa and an extinction coefficient of 78.5 mM^-1^ cm^-1^ at 419 nm) in 20 mM Tris-HCl pH 8.0, to which was added a total of 100 nL of precipitant solution (2.05 M ammonium sulfate, 100 mM Tris-HCl pH 8.0) including crystal seed suspension made by manually crushing the batch crystals leftover from SFX experiments with a Crystal Crusher (Hampton Research) 200 times. The resulting drops were equilibrated against 35 µL of precipitant solution in the reservoir. Crystals appeared within 1 d. X-ray diffraction data at 20 °C were collected directly from crystals within the crystallisation plate on beamline VMXi (Diamond Light Source)^18–20^. The flux of the attenuated 10 × 10 µm beam was 9.93 × 10^11^ ph/s at 16 keV. Wedges of 60° rotation were recorded using a Eiger2 X 4M detector (Dectris). The “fresh” X-ray diffraction dataset was collected from 45 wedges 2 d after starting the crystallisation (Fig. S2A). The “aged” dataset was collected from 35 wedges on other crystals (<u>Fig.</u> S2B) 51 d after starting the crystallisation, by which time the crystals had stopped growing. The data were processed with multi-xia2 dials^21^ in ISPyB by specifying the cell parameters and space group. CCP4i2^13^ was used to generate a free R set and use REFMAC5^14^ and Coot 0.9^15^ to iteratively refine and build the structures based on the SFX structure. JLigand^22^ was used through CCP4cloud^23^ to prepare the restraints for heme DCu.

#### Anaerobic cryocrystallography (ferrous cryo)

Obtaining a reduced (ferrous) crystal structure required extensive optimisation of an anaerobic procedure together with cryo-trapping of the reduced form to avoid reoxidation. The crystal was obtained in an anaerobic chamber (Belle Technology) by sitting-drop vapour diffusion in XRL™ plates with Micro-Bridges® (Hampton Research) sealed by ClearVue™ sheets (Molecular Dimensions). The reservoir contained 1 mL of precipitant solution (1.9 M ammonium sulfate, 100 mM Tris-HCl pH 8.0). The drop contained 1 µL of this precipitant solution, 1 µL of seed stock in the same precipitant solution and 2 µL of McP460 at 7 mg/mL. The large fragments in the seed stock had been removed by centrifugation at 100 *g* for 2 min. Crystals appeared within 3 d and were harvested at 4 d inside the anaerobic chamber. To do so, 3.4 µL of cryoprotectant were added to the drop (precipitant solution containing 1.7 M sodium malonate) and after 7 min, 0.6 µL of precipitant solution containing 1 M sodium dithionite (SDT) was added (for a final SDT concentration of 0.1 M in the drop). This caused a visible change in crystal colour from brownish green to green (Fig. S3). After 10 min, the crystal was harvested in a mounted 80 µm clear CryoLoop (Hampton) and cryocooled from within the anaerobic chamber. X-ray diffraction data were collected at 100 K on beamline I24 (Diamond Light Source) using a CdTe Eiger2 9M detector (Dectris). A 50 x 50 µm beam of X-rays at 20 keV delivered 0.20 MGy of radiation dose for the first dataset and a further 0.30 MGy for the second dataset.^24^ The data were processed with xia2 dials. CCP4i2^13^ was used to generate a free R set and use REFMAC5^14^ and Coot 0.9^15^ were used to iteratively refine and build the structures based on the previously published 100 K structure^3^. The resolution cut-off of 1.33 A was chosen using PAIREF^25^ as including data from 1.35 A to 1.33 A decreased the Rfree value.

#### Single-crystal spectroscopy analysis

UV/Visible absorption spectroscopy from protein crystals maintained at 100 K in a nitrogen cryo stream was performed *in situ* at Diamond beamline I24 using an off axis microspectrophotometer. A fibre coupled xenon light source (Thorlabs), a Shamrock 303i spectrometer (Andor technology) and a Newton EM CCD detector were used. Data were collected over the wavelength range 203-751 nm with a focal spot of 50×50 µm for the white light incident on the crystal. The ferrous crystal was maintained at 100 K in a nitrogen cryo stream. Each spectrum was an accumulation of 10 exposures of 9.95 ms. Spectroscopy data were measured prior to and following X-ray data collection, with the crystal was returned to its original orientation following diffraction data collection in order to ensure that orientation depending changes to the visible spectrum were avoided. The ferric crystal was harvested using a copper pint then inserted into a MicroRT™ tube (MiTeGen) This plastic sleeve was filled with enough mother liquor to have the crystal near but not immersed into it. Each spectrum was an accumulation of 50 exposures. No diffraction data was collected. Smoothing was performed in Origin by LOESS of span 7%.

#### Solution spectroscopy

McP460 at 7 mg/mL in 20 mM Tris pH 8.0 was added as a 10 µL aliquot to 1 mL of each pH 8.0 solution. Spectra were recorded from 190 nm to 840 nm (1 nm resolution) in a reduced volume polystyrene cuvette of path length 1 cm on a Nanodrop 2000c (ThermoFisher). Smoothing was performed in Origin by LOESS of span 7%.

### Computational simulation of cytochrome P460

#### Determination of P460 heme partial charges

P460 has a unique heme C unit where an additional covalent bond is formed between the NZ atom of Lys78 and CHA of heme and charges were assigned following the established conventions for CHARMM forcefields (FF)^26^. A cluster model (Fig. S4) consisting of the heme cofactor (without the propionate and methyl side chains) with the sidechains of Lys78, covalently linked Cys140, Cys143 and coordinated His144 was built from the published 6HIU^3^ structure for charge evaluation. DFT geometry optimisations were carried out using the NWChem^27^ program using the B3LYP^28,29^-D3^30^ functional and 6-31G*^31,32^ basis set. The CB-CA bond for the amino acid resides were terminated by hydrogen atom and the CB positions were kept fixed to their crystallographic coordinates during the DFT optimisations. Optimisations were performed in both oxidised (ferric) and reduced (ferrous) forms of Fe treated in three different spin states: doublet (M=2), quartet (M=4) and sextet (M=6) for the ferric form and singlet (M=1), triplet (M=3) and quintet (M=5) for the ferrous form, respectively. The optimisations were followed by an electrostatic potential (ESP) charge calculation with added constraints to ensure that chemical groups had an overall neutral charge and similar groups had equivalent charges, in line with CHARMM conventions. The charges used correspond to the lowest energy spin state for the optimised cluster and are provided in the SI, Table S2.

##### Modelling the crosslinks

The damage-free, high resolution **ferric SFX** structure was used as a starting structure for generating the alternative crosslink species. The nature of the crosslink in the **ferric SFX** structure is ambiguous with Lys78-CD – Heme C2A distances of 1.94 Å and 2.11 Å for chain A and B, respectively. Single crosslink (SC) and double crosslink (DC) geometries were created from this structure (Fig. 1) and then partial optimization of the heme unit around the crosslink was performed to relax the generated geometries. To this end, cluster calculations were performed where all the heme atoms, except those involved in the crosslink and their two immediate neighbouring atoms were fixed (see Fig. S5). Both oxidized and reduced forms of Fe were calculated for each of the spin states used in the charge determination step. The resulting optimised geometries from all spin states were similar for both the oxidised ferric and reduced ferrous state. The structural parameters around the SC and DC heme units were consistent across all spin states and oxidation state of Fe and are provided in the SI (section S.2). The resulting optimised DC and SC heme C centers in the lowest spin state (M=2 for ferric and M=1 for ferrous) were aligned to the crystal heme C unit of the **ferric SFX** structure. The alignment showed good agreement at the CA – CB bond junctions in Lys78, Cys140, Cys143 and His144 (Fig. S6). The heme C unit from the **ferric SFX** structure was then replaced by the optimised SC and DC heme units, thus providing the starting structures for the SC and DC models.

**Fig. 1.**
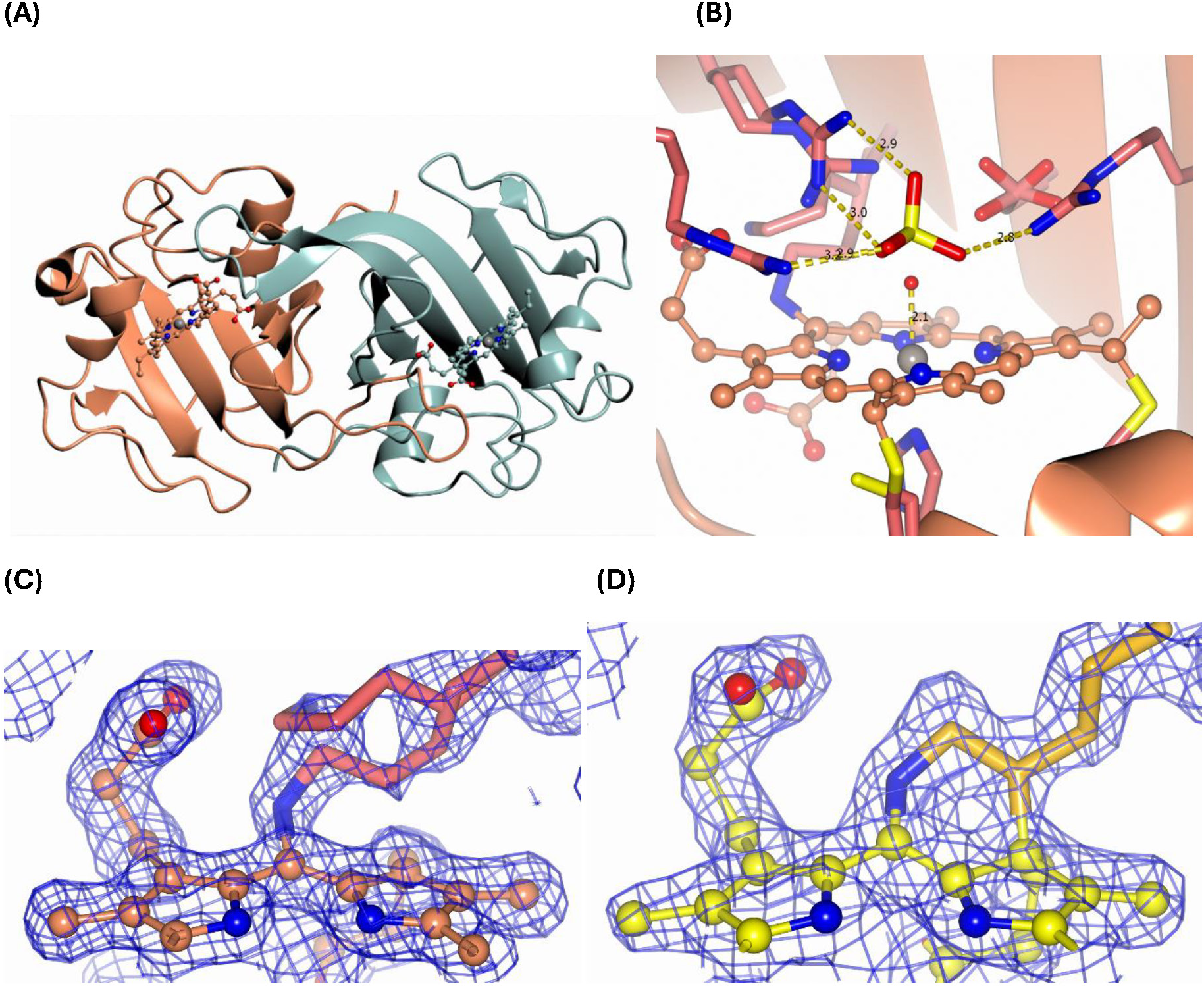
Room temperature crystal structures of McP460. The polypeptide is represented in ribbons, the heme cofactors in ball and stick and the amino acid sidechains in cylinders of a redder color. (A) **SFX** whole protein with chain A in sea green, (B) **SFX** active site with sulfate represented as cylinders. Bond lengths are indicated in Å (C) **SFX** crosslink electron density, (D) **RT fresh** crosslink electron density.

##### Simulation setup

The generated Ferric-SC and Ferric-DC structures were prepared for simulations by removing all double occupancy mainchain and sidechain atoms and partial water molecules, keeping those with greater fractional occupancy or lower B-factors. The protonation of the titrable residues of McP460 crystal structure were adjusted to pH 8 to be consistent with the pH of crystallisation using propKa^33,34^ module of the PDB2PQR^35^ suite of programs, followed by visual inspection of side chain environments. All ionizable residues were in their standard protonation states, with histidine residues singly protonated at ND. Along with setting the protonation state, PDB2PQR also added missing hydrogen atoms to the protein structures as appropriate. The water molecules present in the crystallographic structure were retained. The partial charges for the Ferric-DC model were adjusted in line with the CHARMM FF conventions for charge determination and are provided in SI (Table S2). The same protocol was repeated to set up the Ferrous models, starting from the **ferrous Cryo** structure.

The above generated structures were subjected to minimization to optimize hydrogen atom positions while maintaining the heavy atoms fixed to their crystallographic coordinates. Once minimised, the structures were ready to be used for QM/MM simulations of the crystal protein. Note that as in previous work,^36^ our QM/MM model is based on a finite cluster representation and does not consider long range periodicity, but we do not expect this to have a significant influence on the local chemical environment around the heme-Lysine crosslink.

##### QM/MM calculations

*QM/MM partitioning for single crosslink models*: In our QM/MM partitioning scheme the heme C unit was included in the QM region along with the covalently linked Lys78, Cys140 and Cys143, the coordinated His144 and water 275, which is coordinated to the Fe atom. The methyl and propionate side chains of the heme unit were not included in the QM region^37^ (Fig. S7A). All the residues were truncated at the neutral CA-CB bond and capped with hydrogen atoms. Atoms within 4 Å of the QM region were relaxed (active region) during QM/MM geometry optimizations, while the remaining atoms were frozen. The size of the unconstrained region was restricted to a small 4 Å window to allow the geometry around the QM region to relax thus avoiding extensive relaxation of the outer most residues of the protein in vacuum during the optimisation as the heme C residue is surface exposed.

*QM/MM partitioning for double crosslink models*: The same residues were considered as in SC structure along with the additional -CH_2_ side chain of pyrrole D ring of the heme C which is involved in the formation of this double crosslink (Fig. S7B).

QM/MM calculations were performed with the Python-based version of the ChemShell computational chemistry environment^38^ using NWChem and DL_POLY^39^ for DFT and molecular mechanics calculations, respectively. The biomolecular QM/MM workflow in ChemShell^40^ was used to perform the calculations including the use of DL_FIELD^41^ to import forcefield parameters in CHARMM format. The DL-FIND^42^ library was used to perform geometry optimizations. The electrostatic embedding scheme with charge shift correction was used to represent the surrounding MM partial charge distribution. The level of theory used for the calculations is B3LYP- D3 and def2-SVP basis sets were used for all QM atoms except Fe, which was treated with the def2-TZVP basis set^43,44^. Optimisations were carried out in three spin states; M=2, 4 and 6 associated with ferric state of Fe. For the Ferrous-SC systems, the system was optimised in three spin states: M=1, 3 and 5. Additionally, this is a dimeric protein, hence the calculations were performed on both chains, considering one chain heme in the QM region at a time.

#### Molecular dynamics (MD) setup

After initial preparation of the SC and DC structures for Ferric- SC/DCs and Ferrous-SC, they were solvated by padding a 20 Å layer of TIP3P^45^ water in all three directions using a pre-equilibrated cubic water box available via the solvate plugin module within VMD^46^. The ferric state was electroneutral, so no additional ions were added, while for the ferrous state two sodium counterions were added to achieve electroneutrality during MD simulation using the autoionisation feature in VMD. Explicit all-atom MD simulations were performed on these systems using NAMD^47^ with the CHARMM36 forcefield^48^. These simulations employed Langevin dynamics with periodic boundary conditions at 293 K. Long-range electrostatics were treated by the Particle Mesh Ewald method. The cutoff for nonbonded parameters was set at 12 Å with a smooth switching function turned on from 10 Å. In NPT simulations, the pressure was maintained with the Langevin piston method. All the systems were initially subjected to 5000 steps of conjugate gradient (CG) minimization to eliminate any unphysical contacts. Then, the water and ions were equilibrated in an NVT ensemble, keeping the protein fixed for 2 ns. This was followed by 100000 steps of minimization of the protein H-atoms to keep the protein structure close to the crystal geometry. This was necessary to ensure that the stability of the simulation using rigid bonds for both water and protein. Next the solvent was allowed to further relax by equilibrating the systems for 175 ns in an NPT ensemble keeping all atoms of the protein fixed. In addition, for Ferric-SC/DC systems, after minimization, the systems were allowed to relax for 175ns in an NPT ensemble keeping the backbone harmonically restrained with a force constant of 25 kcal/mol/Å^2^ while allowing the side chains and solvent to relax.

#### Spectral calculations

The QM/MM optimized crystal structures from chain A served as the basis to compute the absorption spectra of the double crosslinked and single crosslinked models. The spectral calculations in ChemShell were carried out using ORCA version 5.0.3^49^ which provides a wide range of suitable quantum chemical methods for calculating excitation energies. Excitation energies were computed for different spin states of the Ferric and Ferrous forms using simplified time-dependent density functional theory (sTD-DFT) and time-dependent density functional theory (TD-DFT). The semi-empirical method ZINDO/S^50^ is an alternative cheap and effective method known to perform well for transition metal containing organic biomolecules parameterised for lower spin states. We also used ZINDO/S to compute excitation energies for the lower spin states. The sTD-DFT^51^ and TD-DFT^52^ calculations were carried out with the range-separated hybrid functional CAM-B3LYP^53^ with a def2-SVP basis set on all atoms other than Fe which was treated using def2-TZVP. The oscillator strengths for all calculations were extracted in the dipole length representation. The resulting spectra were produced by convolution of the first 30 transitions for the Ferrous form and 50 transitions for the Ferric form using a Gaussian function with a broadening of 0.15 eV. The electronic transitions were analysed in terms of natural transition orbitals (NTO) which were generated using the TheoDore^54^ program.

## Results

### Enzyme activity assays

Activity assays were carried out to assess the protein’s ability to oxidise hydroxylamine. The reported activity of NeP460 at 4.5 μMDCPIP·(μMCytP460)^-1^·(min)^-1^ is around 3.8 times higher than that of McP460 (SI Fig. S8). The activity of McP460, 1.19, is more similar to that of the NsALP460 A131E mutant at 2.1.^55^ This suggests that while McP460 is able to oxidise hydroxylamine it may not be as efficient as NeP460. Production of nitrous oxide/nitrite by the wt protein was confirmed though GC-MS and Griess Assays. Under anaerobic conditions only nitrous oxide was detected whilst under aerobic conditions both nitrous oxide and nitrite could be seen (SI Fig. S8).

### Serial femtosecond crystallography structure of cytochrome P460

We first sought to obtain structures of ferric McP460 free of any artefacts rising from radiation damage, cryoprotectants or temperature -dependent effects. Such a structure provides the best starting point for further molecular simulations. The most effective means to achieve this was to use serial femtosecond crystallography (SFX) at an XFEL where the very short (10 fs) pulse duration allows diffraction data to be measured from room temperature microcrystals without sufficient time being available for radiation chemistry to occur (the principle of diffraction before destruction). Using room temperature microcrystals also avoids any artefacts from cryo- protection or flash cooling. Accordingly, SFX data were measured at the SACLA XFEL and produced a structure to 1.28 Å resolution, table 1. Such high resolution is rarely achieved in SFX (only two room temperature SFX structures at higher resolution were present in the protein data Bank at the time of writing).

**Table 1:**
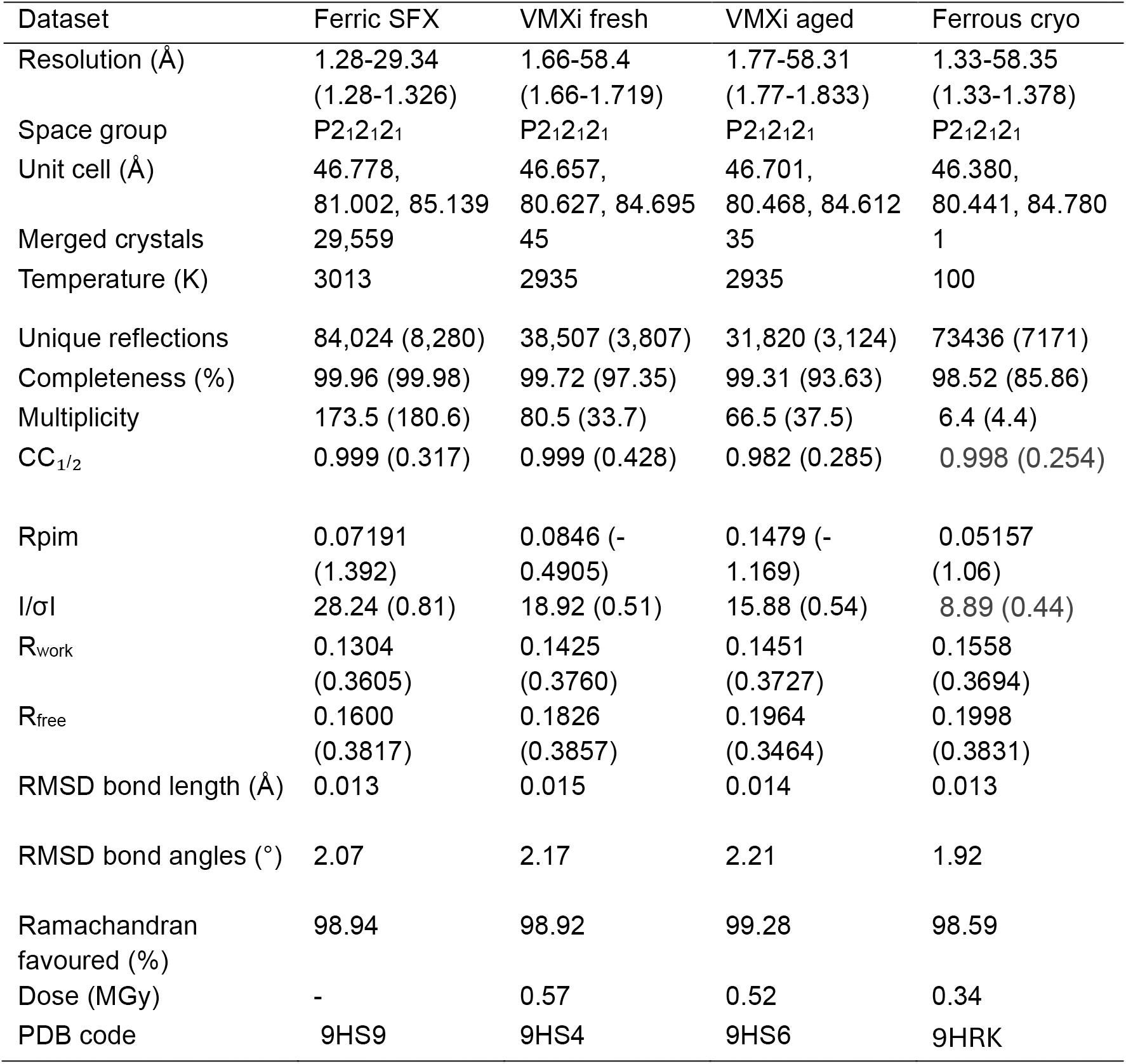
Experimental data statistics for the crystal structures of McP460 reported herein

| Dataset | Ferric SFX | VMXi fresh | VMXi aged | Ferrous cryo |
| --- | --- | --- | --- | --- |
| Resolution (Å) | 1.28-29.34<br>(1.28-1.326) | 1.66-58.4<br>(1.66-1.719) | 1.77-58.31<br>(1.77-1.833) | 1.33-58.35<br>(1.33-1.378) |
| Space group | P2<sub>1</sub>2<sub>1</sub>2<sub>1</sub> | P2<sub>1</sub>2<sub>1</sub>2<sub>1</sub> | P2<sub>1</sub>2<sub>1</sub>2<sub>1</sub> | P2<sub>1</sub>2<sub>1</sub>2<sub>1</sub> |
| Unit cell (Å) | 46.778,<br>81.002, 85.139 | 46.657,<br>80.627, 84.695 | 46.701,<br>80.468, 84.612 | 46.380,<br>80.441, 84.780 |
| Merged crystals | 29,559 | 45 | 35 | 1 |
| Temperature (K) | 3013 | 2935 | 2935 | 100 |
| Unique reflections | 84,024 (8,280) | 38,507 (3,807) | 31,820 (3,124) | 73436 (7171) |
| Completeness (%) | 99.96 (99.98) | 99.72 (97.35) | 99.31 (93.63) | 98.52 (85.86) |
| Multiplicity | 173.5 (180.6) | 80.5 (33.7) | 66.5 (37.5) | 6.4 (4.4) |
| CC<sub>1/2</sub> | 0.999 (0.317) | 0.999 (0.428) | 0.982 (0.285) | 0.998 (0.254) |
| R<sub>pim</sub> | 0.07191<br>(1.392) | 0.0846 (-<br>0.4905) | 0.1479 (-<br>1.169) | 0.05157<br>(1.06) |
| I/σI | 28.24 (0.81) | 18.92 (0.51) | 15.88 (0.54) | 8.89 (0.44) |
| R<sub>work</sub> | 0.1304<br>(0.3605) | 0.1425<br>(0.3760) | 0.1451<br>(0.3727) | 0.1558<br>(0.3694) |
| R<sub>free</sub> | 0.1600<br>(0.3817) | 0.1826<br>(0.3857) | 0.1964<br>(0.3464) | 0.1998<br>(0.3831) |
| RMSD bond length (Å) | 0.013 | 0.015 | 0.014 | 0.013 |
| RMSD bond angles (°) | 2.07 | 2.17 | 2.21 | 1.92 |
| Ramachandran<br>favoured (%) | 98.94 | 98.92 | 99.28 | 98.59 |
| Dose (MGy) | - | 0.57 | 0.52 | 0.34 |
| PDB code | 9HS9 | 9HS4 | 9HS6 | 9HRK |

As was the case in the previous structure from a single crystal at 100 K, cytochrome P460 forms a homodimer with each monomer adopting a largely beta sheet fold connected by several loops. Superposition of the SFX structure with the 100 K structure (PDB 6HIU) yielded an RMSD value of 0.30 Å over 286 residues. The region of the backbone that differs most between cryogenic conditions and room temperature is the loop from Thr85 to Ser91, which harbours the Phe89 residue that is in contact with the heme from the other monomer. Indeed, this loop is shifted by approximately 0.5 Å in chain A and 0.4 Å in chain B (Fig. S8A and B) based on analysis by Gesamt.^56^ At room temperature, it is therefore closer to the heme of the other chain. The heme groups and heme pockets in the SFX structure of ferric P460 reveal six coordinate Fe(III) geometry with a His at the proximal face with an Fe-His (N) bond length of 2.1 Å.

Unlike the previously published 100 K structure where a cryoprotectant molecule is bound to Arg43 (Fig. S9), the SFX structure shows a sulfate molecule on that site (Fig. 1B). This happens through potentially charged hydrogen bonds (N to O distances of 2.8 Å and 3.1 Å in chain A, 2.9 Å and 3.2 Å in chain B). The presence of the sulfate may explain the differences in the conformations of Arg 50. For example, the major conformation on Arg50 in the SFX structure is the one binding the sulfate with potentially charged hydrogen-bonds (N to O distances of 2.9 A and 3.1 Å in chain A, 2.9 and 3.0 Å in chain B). The minor conformations of Arg50 could affect its interactions with the propionate of the heme. Arg121 also forms a potentially charged hydrogen-bond to the sulfate (2.9 Å between its nitrogen and a sulfate oxygen in chain A, 3.0 Å in chain B).

The SFX structure reveals a new conformation for the sidechain of Gln46A, a very mobile residue according to B-factors. It also shows hydrogen bond between the sidechains of Ser80 and Asn48 that could be involved in a proton relay between the iron-bound ligand and the surface (through Asp102, HOH403 and HOH441, better seen in chain B). The largest shift between the r.t and 100 K structures is from Lys60 on the surface of chain B, at a crystal contact with chain A of another dimer. A minor r.t conformation of Arg66A not seen at 100 K may force Lys60B further away than in its 100 K conformation. An indicator of radiation damage in the 100 K structure is the iron-water distance of over 2.3 Å compared to the 2.1 Å distance in the damage-free SFX structure^57^. This shorter iron-water distance in the SFX structure could explain the why the sidechain of its Asp102 is 0.2 Å closer to the iron centre (based on the distance to the CG carbon) than in the 100 K structure.

B-factor analysis indicated regions that the sidechain of His156B is more disordered than that of His86A which lies some 3.5 A away. Surprisingly, the opposite is the case in the 100 K structure. Anisotropic B-factors allow us to see the direction of displacement to be estimated. In the case of the iron-bound water, its position varies in the direction of Asp102 (Fig. S10), so it may depend on which of the two conformations Asp102 assumes. The sulfate in the active site pockets undergoes a rocking motion, followed by Arg50. The atoms in Tyr130 and Glu139 that are close to the entrance to the active site have some thermal motion towards it, especially their terminal oxygen.

Intriguingly and in contrast to the previous 100 K structure of NeP460, the **SFX** structure of McP460 to very high resolution contained electron density features suggestive of a second cross link between residue Lys78 and the porphyrin. Indeed, the model of the main conformation of Lys78 causes an overlap between its CD carbon and the C2A carbon of the heme (1.4 Å overlap in chain A and 1.3 Å in chain B, 0.2 Å more severe than in the published 100 K structure). The chemical nature of this new modification was unclear due to the multiple conformations of the Lys78 sidechain.

Intriguingly, a spectral shift occurs in the UV-vis spectrum of McP460 on exposure to ammonium sulfate in solution (Fig. S11). Testing of different compounds containing ammonia or sulphate ions suggested that ammonia was the species responsible for the observed spectral shift. We therefore investigated whether ammonia could have bound to the heme group (potentially displacing any heme coordinated water) in crystals grown in high concentrations of ammonium sulphate as precipitant. Importantly, single-crystal spectra consistently show spectra consistent with ferric protein in the absence of ammonia and we do not observe any evidence of ammonia binding in crystals. (Fig. 2B).

**Fig. 2.**
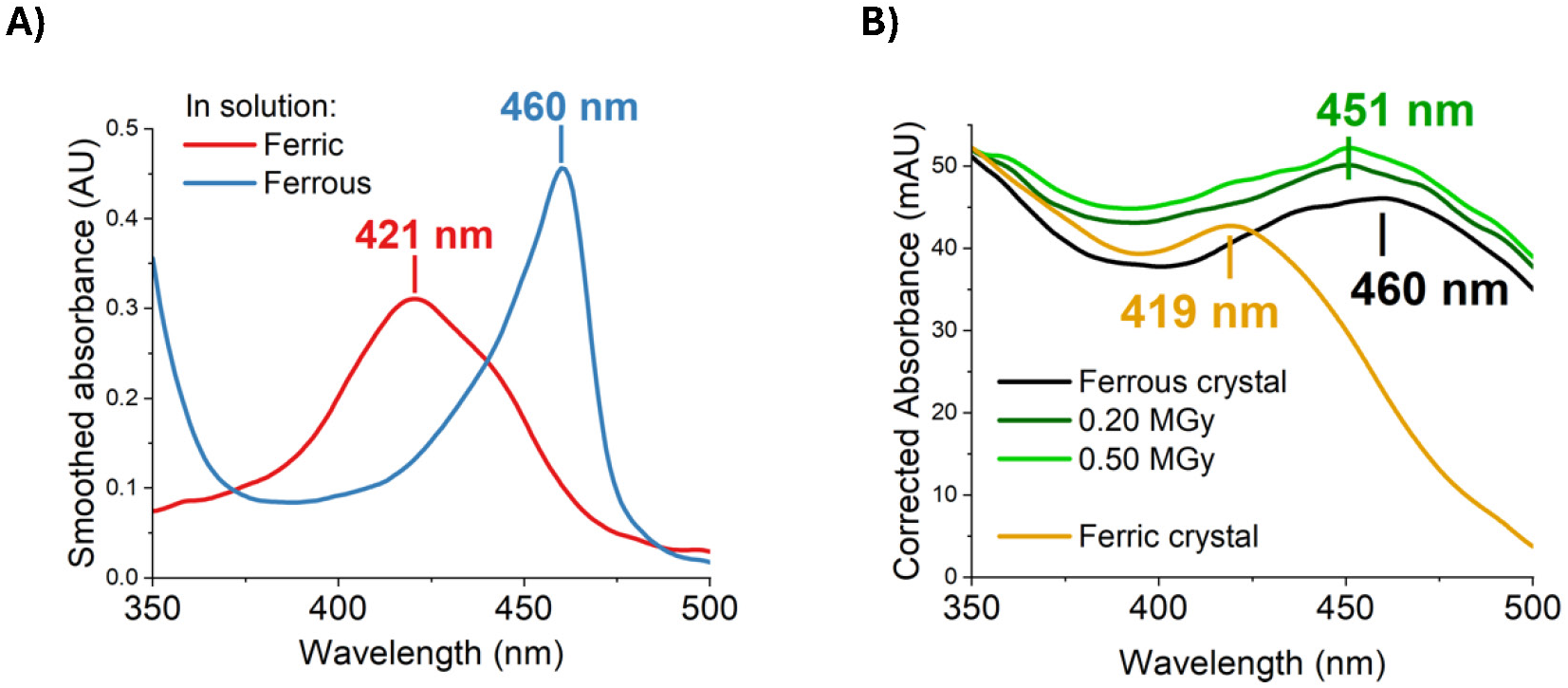
Experimental UV-vis spectra of McP460 at pH 8.0 A) in solution (smoothing of 2.5% only due to the sharp 460 nm peak) and B) in crystals. A clear shift is evident upon changes of oxidation state

### Fresh and aged crystal structures (double-crosslink only)

To ascertain whether degradation of crystals over time was responsible this variability, we collected diffraction data from crystals *in situ* 2 d after setting the crystallisation plate, and again 7 weeks later using other crystals from the same plate. Pleasingly, the fresh crystals yielded a structure with a clearly resolved conformation of Lys78. The aged crystals gave a structure that was essentially identical, albeit at a lower resolution. This suggests that the additional Lys78 conformation is not caused by degradation over time. We could only speculate that it is due to a combination of the batch of protein or the method of crystallisation.

This structure collected on fresh crystals clearly shows a second crosslink between Lys78 and the porphyrin: a covalent bond between the CD carbon of Lys78 and the C2A carbon of the porphyrin (Fig. 1D). Since there are now four carbons bonded to C2A, they must all do so through single bonds to satisfy its valence. This means the C2A-C3A bond has been changed from double to single (Fig. 3). The geometry around C3A having remained clearly planar in the structure, we assign it an exocyclic double-bond, as evidenced by a shorter bond length to the terminal carbon on this tetrapyrrole ring D compared to all others (1.46 Å vs 1.52 Å when using restraints without this double-bond; 1.33 A Å when using double-bond restraints).

**Fig. 3.**
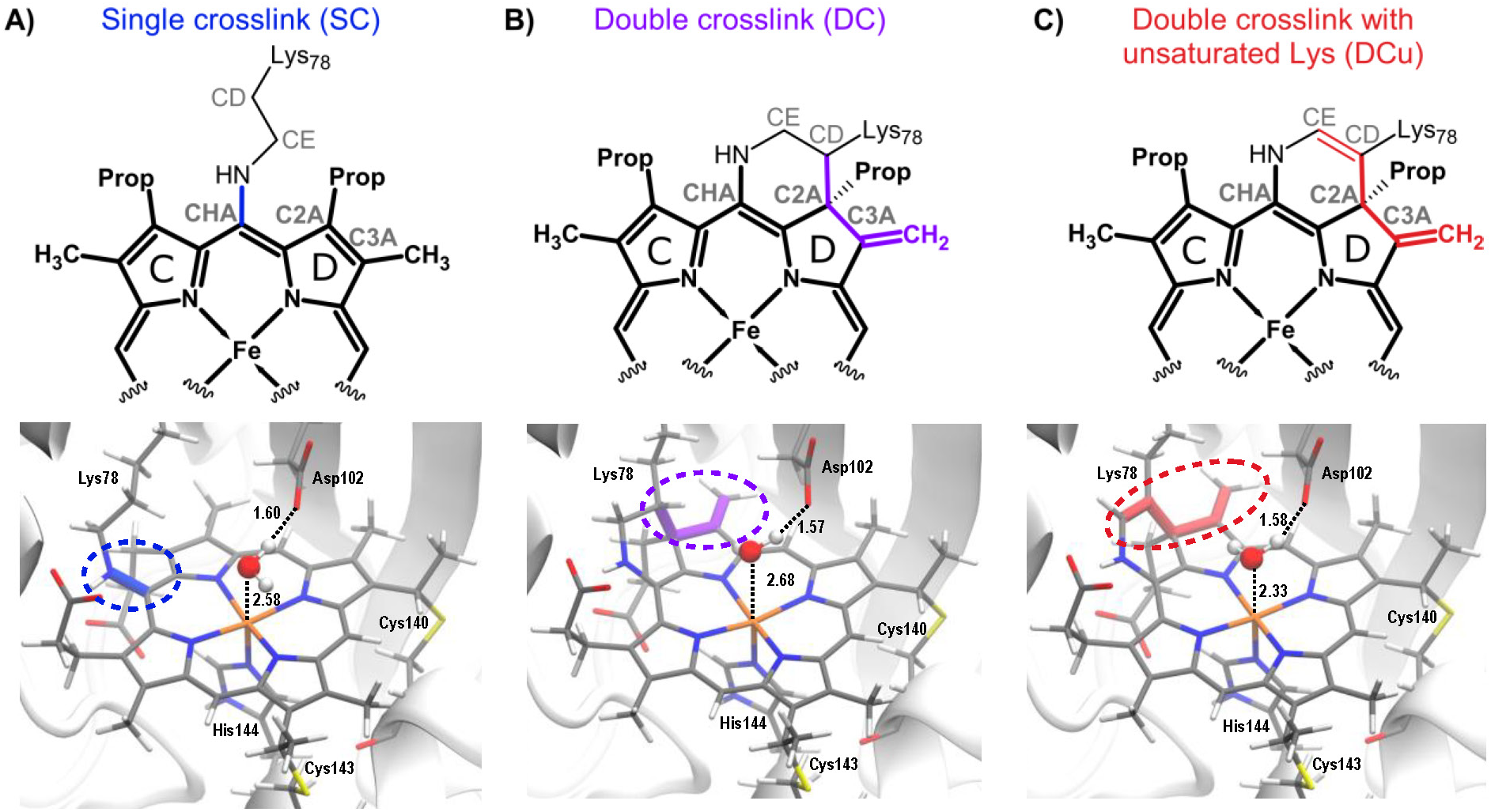
QM/MM optimised geometries of A) single crosslink, SC, B) double crosslink, DC and C) double crosslink with unsaturated Lys side chain, DCu. The heme C unit is shown with sticks and the different crosslinks are shown in the same colour as in Scheme 1. The 6^th^ site coordinated water is shown in ball and stick. The active site residue Asp102 which interacts with the coordinated water is shown in sticks. Distances are given in Å.

When examining the geometry around the CD of Lys78 in the electron density, it was unclear whether it had four covalently bonded atoms (with a hydrogen pointing towards the iron-bound water) or three bonded atoms thanks to an additional double-bond on the lysine. We therefore reduced model bias by calculating a composite omit map (Fig. S12). This shows a mostly planar arrangement of atoms around the Lys78 CD, supporting a double-bond on the lysine sidechain. This also provides the reactivity needed to form a second crosslink, especially if α to a heteroatom. We therefore tentatively built a double-bond between the CD and CE carbons of Lys78 in the RT fresh and RT aged models.

Electron density near the heme-linked sulfur atom of Cys143 was assigned to a minor alternate conformation of the residue. Since this other sulfur position is close to bonding distance from the heme carbon CAC (1.9 Å), it was also modelled as linked. Because of the nearly planar geometry around CAC from this new bond, there could remain a C-C double bond to the terminal carbon CBC. This can unfortunately not be distinguished within the mainly single bond from the major conformation of Cys143. We do not expect the UV-vis spectrum to be affected as this double-bond is not in plane of the heme and therefore would not be conjugated with its π-system.

The iron in the **RT fresh** structure is likely to have remained in the ferric state, as X-rays seem unable to reduce it in McP460 based on single-crystal UV-vis spectroscopy (Fig. S13). The **RT fresh** structure is very similar to the damage-free **SFX** structure (RMSD 0.14 Å across 286 residues). We built fewer alternate conformations in the **RT fresh** structure, likely due to its lower resolution. Most differences are in the alternate conformations (such as Gln46B,B), which were not used in simulations of the **SFX** structure. The iron-water distances are similar in both structures (2.1 Å ). The **RT aged** structure is essentially identical, apart from fewer waters and alternate conformations due to a lower resolution. The conformations of Gln46 in the active site are slightly different, but it is a very mobile residue, evidenced by its high temperature factors.

### Cytochrome P460 in the ferrous state (structure reduced *in crystallo*)

We next sought to understand the nature of the heme structure in the ferrous form of the enzyme. Initial attempts to reduce the protein in the crystalline state using chemical reagents or by exposure to X-rays were unsuccessful as assessed by electron density (checking for movement of the iron coordinated water molecule) and crystal spectroscopy (checking for optical spectra changes that would indicate reduction). Preparation of crystals in an anaerobic glove box followed by incubation with sodium dithionite and cryo- trapping resulted in successful generation of one crystal in the ferrous form as assessed by single-crystal spectroscopy of cryo-trapped crystals after reduction, Fig. 4 and S14. The difficulty of obtaining a stable reduced form rendered it unfeasible to obtain a room temperature or SFX structure of the ferrous protein. The optical absorption spectrum revealed a large spectral shift relative to the as isolated ferric form consistent with successful reduction. Crystallographic data were measured from this reduced crystal yielding a structure at 1.33 Å resolution. A superposition of the ferrous structure with the previous ferric structure (PDB 6HIU) and with the **SFX** structure of the intact ferric state revealed structural changes including the iron atom being five coordinate in the ferrous form (and therefore moved towards the distal side of the plan of the heme, see Table 2) and the loss of the iron-coordinated water molecule present in the ferric form. Instead, a water is 3.1 Å from the nitrogen of either pyrrole ring A or D, with an additional water 3.1 Å from the nitrogen of pyrrole ring C. Residue Lys78 only shows the single crosslink between its sidechain nitrogen and the γ*-meso* carbon of the heme (CHA carbon), not the crosslink between its CD carbon and the C2A carbon of the heme (Scheme 1B/C). Consequently, the terminal carbon of the pyrrole ring D is single-bonded (1.51 Å).

**Fig. 4.**
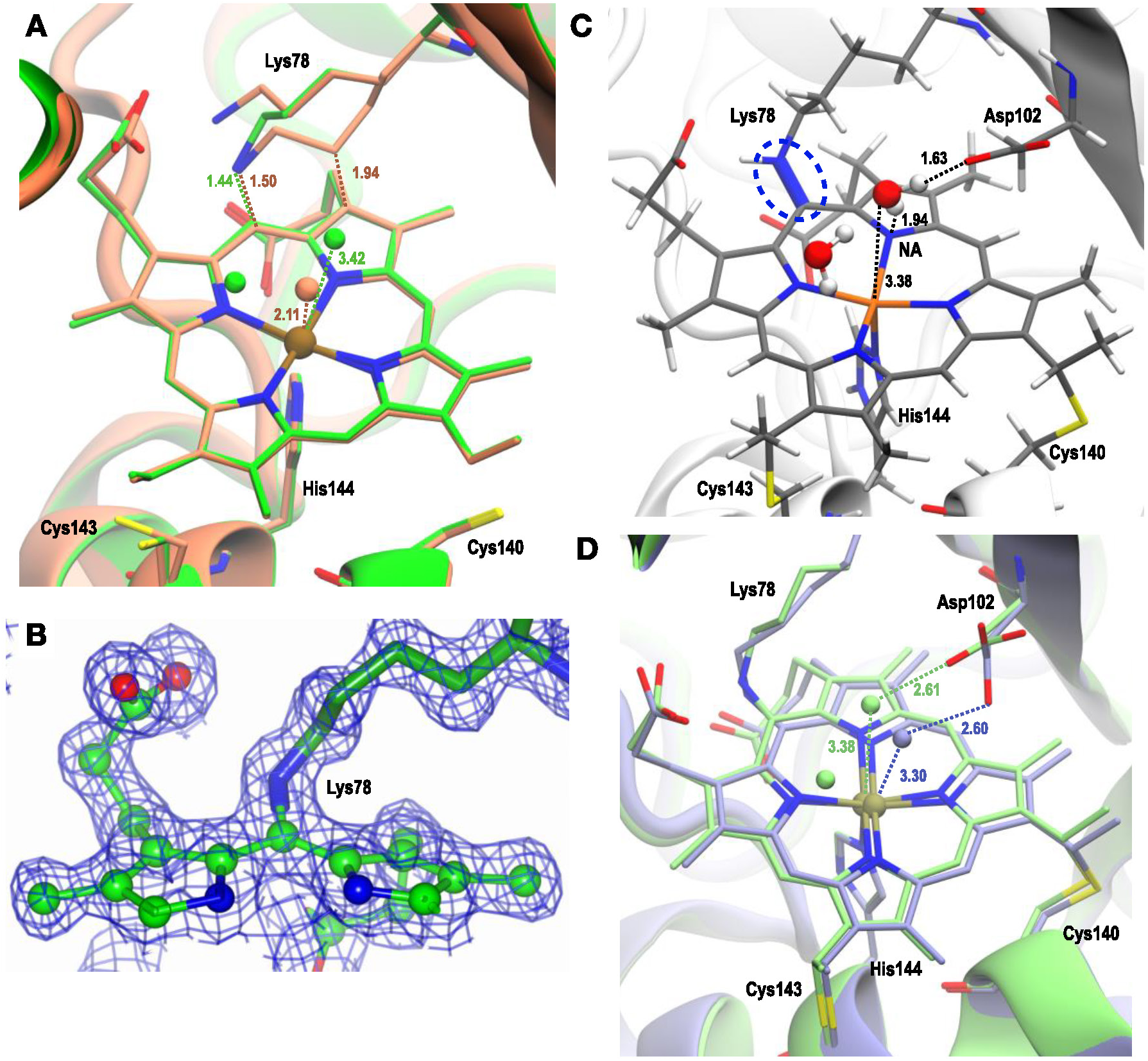
A) **Ferrous Cryo** structure (green) overlapped with **ferric SFX** (brown) structure; the crosslink with Lys78 and water(s) near Fe are shown. The active site residues and H-atoms are omitted to give a clear view around the heme unit and crosslink(s). B) Electron density map showing SC character of the **ferrous Cryo** structure. C) QM/MM optimised Ferrous-SC showing SC (highlighted as blue dashed lines), active site residue Asp102 and the water molecules near Fe. D) QM/MM optimised Ferrous-SC (light green) overlapped with QM/MM optimised *SimFerrous-SC* (light blue); the water(s) near Fe and active site residue Asp are shown. The H- atoms are not shown to enable a clear view around the heme unit and SC. All distances are in Å.

**Table 2.** – Heme C site parameters for McP460 structures, including: the distance between proximal His143 N and Fe (Fe-HisN), the distance between the coordinated water to Fe (Fe- water), Fe to porphyrin N of pyrrole ring D (Fe-PyrNA), the distance between Lys78 and CHA of heme (LysN-CHA), the distance between CD of Lys78 and C2A of heme in DC systems (LysCD- C2A), the distance between heme C3A and CMA distance (C3A-CMA) that undergoes exocyclic modification in DC systems and Fe out of plane motion (Fe-OOP).

|  | Fe-HisN<br>(Å) | Fe-water<br>(Å) | Fe-PyrNA<br>(Å) | LysN-CHA (Å) | LysCD-C2A (Å) | C3A-CMA (Å) | Fe-OOP<br>(Å)* |
| --- | --- | --- | --- | --- | --- | --- | --- |
| **Experiments** |  |  |  |  |  |  |  |
| **Ferric SFX** | 2.14 | 2.11 | 2.03 | 1.46 | 1.94 | 1.43 | 0.06 |
| **Ferric RT Fresh** | 2.11 | 2.15 | 2.06 | 1.40 | 1.58 | 1.33 | 0.11 |
| **Ferrous Cryo** | 2.10 | 3.33 | 2.10 | 1.44 | - | 1.53 | 0.35 |
| **Simulations** |  |  |  |  |  |  |  |
| **Ferric-SC (M=6)** | 2.09 | 2.58 | 2.08 | 1.46 | - | 1.47* | 0.24 |
| **Ferric-DC (M=6)** | 2.11 | 2.68 | 2.22 | 1.34 | 1.57 | 1.35 | 0.20 |
| **Ferric-DCu**<br>**(M=2)** | 1.93 | 2.33 | 2.10 | 1.38 | 1.54 | 1.35 | 0.12 |
| **Ferrous-SC**<br>**(M=5)** | 2.11 | 3.38 | 2.14 | 1.35 | - | 1.47* | 0.40 |
| **Ferrous-DC (M=5)** | 2.11 | 3.26 | 2.15 | 1.37 | 1.57 | 1.35 | 0.30 |
| **Ferrous-DCu (M=5)** | 2.13 | 3.21 | 2.14 | 1.38 | 1.54 | 1.35 | 0.25 |
| **SimFerrous-SC (M=5)** | 2.16 | 3.30 | 2.08 | 1.41 | - | 1.47* | 0.29 |
| **SimFerrous-DC (M=5)** | 2.15 | 2.61 | 2.11 | 1.37 | 1.56 | 1.35 | 0.21 |
| **SimFerrous-DCu (M=5)** | 2.16 | 2.5 | 2.12 | 1.39 | 1.53 | 1.35 | 0.19 |

\*CMA was not a part of the QM region but of the MM region in these structures.

The single-crystal UV-vis spectrum of the anaerobically reduced crystal showed a 460 nm absorbance peak (Fig. 2B), which matches the value from solution experiments. However, after collection of the X-ray diffraction data (0.20 MGy dose), the peak shifted to 451 nm. The structure presented here came from a second diffraction dataset from the same crystal (additional 0.30 MGy of dose), so that the UV-vis spectra before and after its collection both had a peak at 451 nm.

### Computational Studies on Ferric-SC/DC/DCDB: Energies and geometrical parameters

P460 is a homomeric protein and the parameters around heme from experimental **ferric SFX** structure indicate very similar heme C geometry in both chains. As each simulation contains only one chain’s heme unit in the QM region, the two chains, chain A and chain B, can be considered as independent systems in our QM/MM simulations. The lowest energy spin state was sextet (M=6) for the chain A heme for both Ferric-SC and Ferric-DC. The energies are provided in SI Table S3A and the optimised structures are provided in Fig. 3. The results for chain B are provided in SI Table S3A. The two chains, however, on many occasions optimised to different spin states; for chain B, the lowest energy spin states for Ferric-SC and DC systems, were M=4 and M=2, respectively. The relative energy of chain B spin states for Ferric-SC and DC with respect to lowest M=6 state of chain A are -2.5 and -1.1 kcal/mol, respectively. These differences are comparable or less than variations in QM/MM energies between the spin states of individual chains, hence for the subsequent spectral calculations we only simulated chain A (see section: Computational Spectroscopy of Ferric SFX and Ferrous Cryo structures).

We also considered the possibility of the DCu crosslink featuring a double bond between CD and CE atoms of lysine as tentatively assigned for DC species from the obtained crystal structures (**RT fresh**). The charges were adjusted to accommodate this double bond and are provided in the SI (Section S.1 and Table S2) and the systems were optimised at QM/MM level from the corresponding DC predecessors. The DCu system optimised to the doublet state (M=2) for chain

A. Chain B, though optimised to the M=4 state. The energy of this system was -2.13 kcal/mol lower in energy compared to M=2 of chain A. The energies are provided in the SI (Table S3A) and the structure in Fig. 3C.

The geometrical parameters measured around the heme unit for chain A are provided in Table 2. The Fe out-of-plane (Fe-OOP) is measured as the distance between the center of mass of the four N atoms of the porphyrin ring to that of Fe. The heme site geometric parameters from QM/MM optimised geometries are very sensitive to the spin states (see SI, Table S4A). The average distance for Fe-His in the QM/MM optimised structures is 2.04± 0.10 Å and falls within the observed distance of 2.11 ± 0.01 Å. The Fe-water distance is longer than that observed experimentally; 2.53 ± 0.18 Å compared to 2.08 ± 0.03 Å from experimental structures. The measured Fe-OOP distance is dependent on the spin state of the optimised geometry. Higher spin states are associated with larger Fe-OOP distances. Hence, the correlation of Fe-OOP distance to experimental structures is difficult. In case of DC and DCu species, one distinguishable observation is the lengthening of the Fe - PyrNA distance by ∼ 0.1Å in M=6 state compared to the same in the M=2 state. This Fe-PyrNA distance belongs to the pyrrole ring D harbouring C2A atom to which the second crosslink with CB-Lys78 is formed. Such elongation is also present, though less in significance, in the crystal structures (SI Table S1).

Comparing geometric parameters around the heme C unit of the two chains shows subtle differences among them. The results of chain B are provided in SI Table S4A. For all type of crosslinks considered; the Fe-hisN bond lengths are larger in chain B than A, both chains show similar lengthening of the the Fe-PyrNA distance, but the main difference is observed in Fe-OOP distances. In chain A the Fe-OOP distance increases with increase in spin multiplicity whereas it is invariant in chain B and is comparable to the Fe-OOP distance measured from the crystal structures (SI table S1). Beyond the coordination sphere of Fe in **ferric SFX,** there are alternative conformations of active site residue Asp102, one oriented perpendicular to another, and one conformation was modelled for chain A and the other for chain B (SI Fig. S15). This Asp102 forms an H-bond interaction with the water coordinated on the 6^th^ site of Fe in chain A, while in chain B, Asp102 forms an H-bond interaction with another active site residue Gln100. Hence changes in conformation in chain A and B cause a different pattern of H-bond interactions around the active site. To verify whether the above hypothesis is correct, we removed the environmental effect by calculating the QM energies in gas phase for the DC systems. Without the environment the energies for the two chains match and they both optimise to the same lowest energy spin state, M=4 (SI table S5). The other spins states, M=2 and 4 are 2.8 and 5.7 kcal/mol less stable than the M=4 state, respectively.

The difference in the active site conformation of Asp102 is also reflected in the water distribution around Fe during MD (detailed analysis is provided in SI, S.3 and Fig. S16). The conformation of Asp102 in chain A facilitates the water to retain the 6^th^ coordination site of Fe. This results in shorter Fe-water distance 2.73 ± 0.20 Å and 2.8 ± 0.21 Å for SC and DC systems, respectively, negligible interaction with pyrrole N and strong H-bond interaction with Asp102. In chain B the water fails to fulfil the 6^th^ coordination site of Fe; the distance is 3.24 ± 0.16 Å and 3.25 ± 0.15 Å, for SC and DC, respectively. It has negligible interaction with pyrrole N similar to chain A, however, it also has ∼ 50% less interaction with Asp 102. This difference in water occupation near Fe, is however lifted in the second set MD simulations where the sidechains were allowed to relax. The results reveal restoration of similar distribution of water around the active site in both chains (SI Fig. 17) for both Ferric-SC/DC systems. In case of the SC system, the Fe-water distances are 2.85 ± 0.24 Å and 2.80 ± 0.25 Å for chain A and B, respectively and that for DC systems they are 2.63 ± 0.16 Å and 2.62 ± 0.16 Å for chain A and B, respectively. The water has negligible interaction with pyrrole N atoms for both chains. The interaction with Asp102 is similar among the two chains but quite different between SC and DC systems. In SC system, water predominantly interacts with Asp102, whereas in double crosslink system this water interacts with both Asp102 and Arg50.

### Computational studies on Ferrous-SC

The experimental **ferrous Cryo** structure featured a CD Lys – C2A heme distance of ∼3.7 Å, indicative of an SC structure. QM/MM optimisation the Ferrous-SC model resulted in a high spin, quintet (M=5) optimised state for both chains. The QM/MM optimised structure observed in chain A is provided in Fig. 4C and the heme C site parameters are provided in Table S4B. Results from chain B are provided in SI (Table S4B) Unlike the Ferric-SC state, the geometrical parameters are in better agreement with the experimental structure: the active site water, (which fulfilled the 6^th^ CN of Fe in Ferric-SC structure) is > 3 Å away from Fe and the Fe-OOP deviation is in accordance with that observed in the X-ray structure (Table S2, S4B).

In this **ferrous Cryo** structure, both chains have equivalent conformation of Asp102 in the active site. Trajectories obtained from MD simulations on Ferrous-SC was analysed for the distribution of water near Fe in the same manner as done for the Ferric-SC system. The last 150 ns of the 175 ns production trajectory reveals that the average distance of Fe-water is 3.33 +/- 0.1 Å and 3.39 +/- 0.09 Å for chain A and B, respectively. The water in chain A had a greater propensity to form H-bond interactions with the N atom of pyrrole ring D than in chain B. The same is reflected in the interaction with active site Asp102 residue. The plots are provided in the SI Fig. 18 left panels A(I) and B(I).

A long CD-C2A distance of ∼3.7 Å in the **ferrous Cryo** structure suggests an SC structure, however, the possibility of other crosslinks cannot be neglected, so DC and DCu systems were generated and simulated. A DC system structure was generated for this system in-silico by first aligning the gas phase optimised DC model (as used in Ferric-DC) onto the crystal structure, assuring good alignment and then replacing the crystal heme unit with the DC heme unit followed by reoptimisation at QM/MM level. The optimised geometry is given in Fig. S19 (I) and energies are provided in table S3B. This Ferrous-DC structure optimised to quintet state and the geometric parameters around heme bear similarities to its SC counterpart (Table 2 and SI Table S4B). The DCu system was also created from DC structure, optimised and the results are provided in Fig. S19(I) and table S3B for energies and S4B for geometric parameters around heme.

### Simulated reduced state

The SC form of the **ferric SFX** structure was reduced in-silico by adding an extra electron to the Ferric-SC system to create the *SimFerrous-SC* system. QM/MM calculations on the crystal protein model and solvation followed by MD were carried out following the same protocol as described above for the Ferric-SC system. QM/MM optimisation of the crystal system resulted in a quintet (M=5) lowest energy state for chain A. The optimised geometry of this *SimFerrous-SC* is overlaid on Ferrous-SC in Fig. 4D and the geometric parameters around heme are provided in Table 2 for chain A. There is significant similarity between the two structures with respect to Fe- water distance, Fe-OOP movement and overall structure of the heme unit. The geometry parameters around heme reflect a longer Fe-water distance and a prominent out plane movement of the Fe and were comparable with that observed in **ferrous Cryo** structure. The energies and geometries for all spin states and for both chains are provided in SI table S3C and S4C.

The MD trajectory was investigated for the same properties, namely distribution of any water molecule within 3.5 Å from Fe and its interaction with heme and active site residues. The residence time of water near Fe shows a difference in distribution among chain A and B as seen in the Ferric-SC system. It also shows distinct difference from that of the Ferrous-SC structure. Chain A shows a bimodal distribution of water molecules in this *SimFerrous-SC* system, one averaging at 2.64 +/- 0.11 Å and the other at 2.94 +/- 0.11 Å. Furthermore, in this *SimFerrous*-SC system, the water in chain A maintains H-bond interaction with the N-atom of pyrrole ring D for more than 50% of time which is absent in chain B. Also, the interaction with active site Asp102 is much pronounced in chain A than B (See Fig. S18 let panels A(II) and B(II)). We also simulated the other crosslinks, DC (S*imFerrous-DC*) and DCu (*SimFerrous-DCu*), for this in-silico reduced **ferric SFX** structure. The optimised geometries are given in Fig. S19(II), and the heme parameters in Table 2 for chain A. The lowest energy spin state was also quintet, like that observed for Ferrous- DC and Ferrous-DCu structures. The heme parameters were also compared in both and provided in Table S4C.

### Computational spectroscopy of Ferric SFX and Ferrous Cryo structures

Computation of electronic absorption spectra is a useful way to help distinguish species present in a crystal or in solution. To shed light on the nature of the crosslink and discern the species present, we have computed the absorption spectrum for the proposed crosslinked models in both Ferrous and Ferric forms. The absorption spectra of the **Ferric SFX** and **Ferrous Cryo** structures computed using TD-DFT with the QM/MM methodology are given in Fig. 5. The experimental absorption maxima of the Soret band for McP460 in the Ferrous and Ferric forms observed in solution are 460 nm (2.69 eV) and 419 nm (2.96 eV) respectively. The experimental spectra for the two forms are given in Fig. 2.

**Fig. 5.**
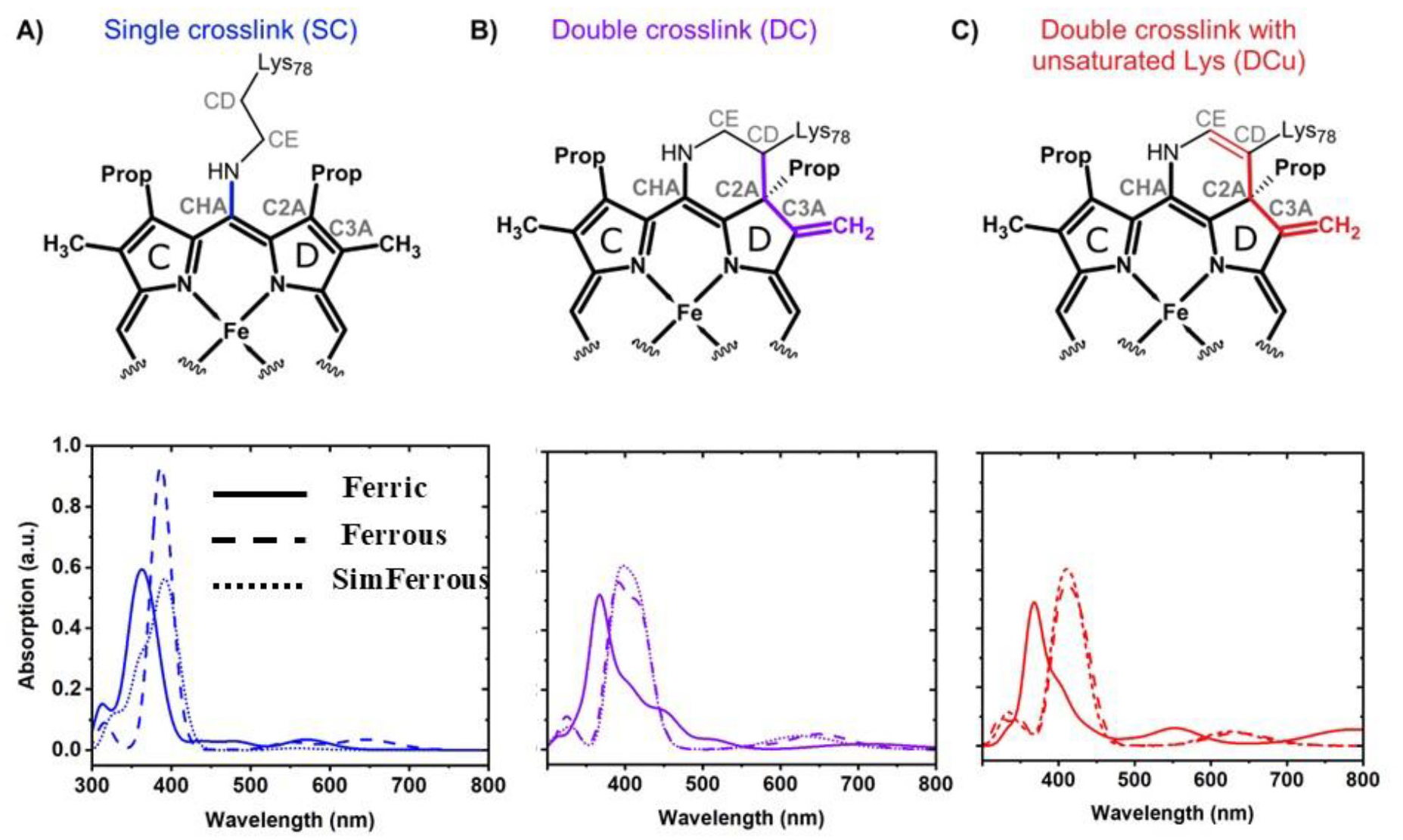
Computed absorption spectra for the three crosslink models using CAM-B3LYP (TD-DFT). Ferrous and *SimFerrous* forms for all models are in a quintet spin state whereas the Ferric forms of DC and DCu are in a doublet spin state and SC is in sextet spin state. The computed absorption spectra for the individual models are given below the structures of each proposed model.

The computed absorption maxima comprising the Soret band using CAM-B3LYP (TD-DFT) for Ferrous-DC and *SimFerrous*-DC for various spin states are given in the SI (see Table S7). The functional CAM-B3LYP has been used to successfully describe the spectrum for heme-based systems in previous studies^58^. The lowest energy spin state for chain A for the ferrous forms is a quintet (M=5). The quintet Ferrous-DC and *SimFerrous*-DC, display absorption maxima at 419 and 418 nm respectively. This peak corresponds to a π-π* transition within the heme moiety (see Fig. S30 in SI for orbitals). Upon introduction of the double bond in the DC models, we observe a red shift in the computed bands. To this end, the Ferrous-DCu and *SimFerrous*-DCu, display peaks at 430 nm and 420 nm respectively. Both DC and DCu forms show a systematic overestimation of the excitation energies compared to the experimental absorption maximum of 460 nm observed for the Ferrous form in solution. We note that the overestimation observed in the case of TD-DFT is a known problem.^59^ The single crosslink models display a blue shift compared to Ferrous-DC models. To this end, Ferrous-SC and *SimFerrous*-SC show maxima at 389 nm and 401 nm respectively (see Table S7 in SI).

In the case of the ferric form, the lowest energy spin state for Ferric-DC and Ferric-SC is a sextet while for Ferric-DCu it is a doublet (M=2). The absorption maximum computed for Ferric-SC is 386 nm while for Ferric-DC, the corresponding value is 388 nm. The doublet Ferric-DCu exhibits a maximum at 397 nm.

The experimental observed spectral shift between the Ferric and Ferrous forms in solution is 0.27 eV. This direction and the shift are reproduced comparing the quintet Ferrous and *SimFerrous* forms for the DCu model with that of the doublet Ferric-DCu form as shown in Fig. 5. The computed shift is 0.21 eV for *SimFerrous*-DCu and 0.2 eV for Ferrous-DCu in the right direction. For the DC model, the sextet Ferric form shows a spectral shift of 0.2 eV (see Fig. S24 in SI). As discussed earlier, the relative energies between the sextet and the doublet for Ferric-DC is ∼1 kcal/mol (see Table S3A in SI), making both spin states accessible. Comparing the doublet Ferric- DC with the Ferrous and *SimFerrous* forms, we observe the right direction of the shift as well and the computed shift is 0.07 eV, as shown in Fig. 5. We note here that the predicted shift for the DC model is lower than the DCu model. In case of the SC model, we observe a lower spectral shift for the sextet ferric form. We note that changing the spin state for the ferrous and ferric forms has a lower effect on the computed spectral shifts.

To further understand the origin of the Soret peak, the electronic transitions computed for the DC and DCu forms were analyzed in terms of natural transition orbitals (NTOs) (see Fig S30 in SI). The NTOs represent the electronic transitions in terms of hole and particles, giving a compact picture for the excitations involved. The computed NTOs for the Soret peak observed for the Ferrous-DC and Ferrous-DCu forms indicate a localized excitation involving all the four rings of the heme. In particular, we observe a localization of the densities on the double bond, providing evidence that the double bond is involved in the electronic transitions.

For comparison, the excitation energy calculations from CAM-B3LYP using simplified time-dependent density functional theory for various spin states and ZINDO/S for the lowest spin states are given in the SI (For sTD-DFT see Fig. S25, Fig. S26 and Fig. S27 and for ZINDO/S see Fig. S20). The sTD-DFT computed maxima for the quintet Ferrous-DC and *SimFerrous*-DC are at 455 nm and 459 nm respectively. The corresponding values for Ferrous-DCu and *SimFerrous*-DCu are 470 nm and 466 nm respectively. All these peaks are in good agreement with the experimental value of 460 nm. In case of the Ferric-DC (M=6) and Ferric-DCu (M=2) forms, the corresponding values are 413 nm and 412 nm respectively. Comparing Ferrous-DCu and *SimFerrous*-DCu with that of Ferric-DCu, the predicted shifts are 0.37 eV and 0.34 eV respectively, in the direction observed experimentally. The corresponding values for the DC models are 0.28 eV and 0.30 eV respectively. Similar to TD-DFT, both DC and DCu models show the right direction of shift for the Soret band compared to experiment (see Fig. S28 and Fig. S29 in the SI). The Ferrous and SimFerrous-SC forms do not display absorption in the range 450-460 nm observed in the DC models. They display maxima at 418 nm and 430 nm respectively. These calculations give confidence in the TD-DFT calculations and provide further evidence for the presence of a double crosslinked species in the crystal.

## Discussion

In this work we combined multiple crystallographic and spectroscopic methods with state-of-the-art computational simulations to elucidate the interaction between heme and protein in cytochrome P460 in the ferric and ferrous oxidation states.

### New crosslink observed experimentally in ferric structures and comparison with simulations on this structure

Our previously published structure of McP460 at lower resolution [PDB 6HIU) showed evidence of a close contact between its heme and the δ-carbon (CD) of the crosslinking lysine. We suspected this to be a sign of a more complex modification than the single crosslink between the lysine nitrogen (NZ) and γ-meso carbon (CHA) of the heme found in NeP460. The electron density for the lysine in 6HIU was difficult to assign to a particular chemical description, likely due to multiple conformations. To ascertain whether the additional conformation was the result of radiation damage, which metalloproteins are particularly susceptible to, it was important to obtain a high resolution structure of the true ground state of the protein free of experimental artefacts. Accordingly, we obtained a structure by serial femtosecond crystallography (SFX) to 1.28 Å resolution. By using an X-ray free-electron laser (XFEL) with 10 fs X-ray pulses, we also avoid any possibility of electronic state changes as a result of exposure to the X-ray beam because the x-ray pulse is too short to allow time for radiation induced chemistry to occur. The structure is one of the highest achieved resolutions for room temperature structures using SFX, which helped provide even greater confidence that a new bond was formed between the lysine and the heme. Intriguingly, even after ruling out the effect of radiation damage, the structure still exhibited multiple conformations of the lysine, making the chemical nature of its crosslink unclear. In contrast, a room temperature synchrotron structure from fresh crystals revealed the position of a single conformation of the lysine with two crosslinks to the heme. This was not affected by crystal ageing, suggesting that it is not crystal size or age that determines the cross- linking but that this is an inherent property of the crystalline protein. We can only speculate about the cause of variation in crosslinking between the SFX and previously published McP460 structure, both with multiple conformations of Lys78, versus the RT fresh and aged structures that exhibit only a double crosslink. Very different crystal sizes were used for the SFX microcrystals than for the three single crystal structures We suggest that the formation of the 2^nd^ crosslink also results in a shifting of the adjacent tetrapyrrole double-bond to become exocyclic, based on its bond length and angle. We speculate that the crosslinking lysine sidechain is oxidised to form a double bond on it, which provides the reactivity to form a second crosslink in an interesting case of C-H activation. Based on the electron density in an omit map of the **RT fresh** structure, we suggest the final structure still harbours a double bond on the lysine.

The QM/MM optimizations of the Ferric-SC/DC/DCu models revealed that the crosslink between Lys and heme C in Ferric-SC/DC and additional double bond on Lys besides the crosslinks in Ferric-DCu were stable minima on the potential energy surface. The structural deviations between the crystal and optimized structures were minimal (Table 2). Simulation also allowed us to inspect the geometrical changes within spin states. The geometric parameters change with increasing spin multiplicity, with significant effects seen on the Fe-water distance and Fe-pyrrole N distances; they increase with increasing spin multiplicity. The latter parameter is very prominent for DC and DCu systems where there is an almost 0.1 Å increase in the Fe-N distance of the pyrrole D ring in both chains. Elongation of one Fe-N distance compared to other Fe-N porphyrin distances is also observed in **RT fresh** and **aged** experimental structures albeit to a lesser extent (changes are less than 0.1 Å). Though the lowest energy spin state for DCu is M=2, it is energetically near degenerate with spin state M=6 (energy difference = 0.9 kcal/mol) Comparing the active site geometries of the DC and DCu systems reveal that the geometrical parameters between them are very similar when considering all the spin states geometries (table 2 and SI table S3A).

The differences seen in the two chains of this dimeric protein in the QM/MM optimisations is most likely related to alternative conformation of the active site residue Asp102. This alteration of Asp102 conformation perturbs the water distribution around heme Fe as seen from MD results when the protein was maintained at its crystallographic positions (Fig S16). When the side chains were relaxed, allowing this Asp102 to adapt to other conformations, the distribution and interaction of the water near Fe becomes equivalent in the two chains (Fig. S17).

The new conformation of Arg50 seen at RT (conformation B in chain A) could affect heme ruffling and crosslink formation through its decreased interaction with the heme propionate oxygen OD2 relative to the conformations previously observed.^61^ Asp102 in McP460 is likely to be involved in a proton relay like the equivalent Glu96 in NeP460.^61^ In the previously published 100 K structure of McP460, Asp102 was further from the iron center (average 5.7 Å from CG to Fe) than in the **SFX** structure (average 5.5 A), which is likely more physiologically relevant due to the absence of radiation damage and cryogenic artifacts. The RT fresh structure is in between (average 5.6 Å). The differences could be due to a lengthening of the iron-water bond with radiation damage^57^, with **RT fresh** being a relatively low-dose structure (32 kGy). In the **ferrous cryo** structure, Asp102 is the furthest from the iron centre (average 5.9 Å), consistent with proton transfer not being needed at this stage of the catalytic cycle, since no ligand is bound to the iron. Proton transfer has been suggested not to occur when the proton relay carboxylate is far enough (e.g. Fe-CG distance of 6.4 A in the crosslink-deficient inactive mutant of NeP460).^61^

### Ferrous Cryo structure and comparing energies and structural parameter with simulations on this structure

Obtaining a structure of the ferrous form of cytochrome P460 proved highly challenging whether using chemical reduction or via electrons generated through X-ray exposure. While crystals change colour in a manner consistent with reduction by sodium dithionite, they are extremely prone to reoxidation under aerobic conditions. In contrast, under anaerobic conditions it was possible to determine a structure from a crystal with a corresponding single-crystal spectrum consistent with the ferrous enzyme. In the crystal structure, the heme distal binding position was vacant, consistent with loss of water molecule upon reduction. This results in 5-coordination of Fe(II), which therefore moves 0.2 Å towards the proximal side of the plane of the heme. This is clearly different to the 6-coordination of Fe thanks to the water in the **SFX** and **RT fresh** structures, giving us confidence that they are indeed in the ferric state. The second crosslink (Lys CD-heme C2A) is broken in the crystallographic ferrous structure, leading to the terminal (CMA) carbon of pyrrole ring D to change from methylene (-CH_2_) to methyl (-CH_3_). This raises the intriguing possibility that this happens as part of the catalytic cycle, and removing the second crosslink could be a way to abstract hydrogen radicals from NH2OH to form N2O. This could apply to HAO too, as it can also exist in single- and double-crosslink form^62^.[6M0Q]

Simulations were carried out with the obtained **ferrous Cryo** structure following the same protocol as used for the Ferric-SC system. The reduced X-ray structure displays a long CD-C2A distance consistent with an SC structure. Following the same trend as seen in the ferric state, the QM/MM optimisation resulted in lowest energy for the quintet spin state. The heme-water distance was 3.26 Å and is in good agreement with that observed for **ferrous Cryo** (3.42 Å). The Fe-OOP distance also showed good agreement; 0.30 Å computed to 0.35 Å observed experimentally. The enhancement of Fe-water distance is also reflected in the MD trajectories, the average distance between 3.33 +/- 0.1 in and 3.39 +/- 0.09 in of the chains A and B, respectively. This loss of water from heme coordination results in the displacement of the water towards pyrrole rings and formation of H-bond interaction with the N of the pyrrole rings, a behaviour not observed in the oxidised state (see Fig. S18 B(I)).

#### Double crosslink systems in Ferrous cryo

For comparison we also prepared DC and DCu geometries starting from the **ferrous cryo** structure (Ferrous-DC/DCu) and compared optimised geometries and simulated UV-visspectra. The optimised geometries are provided in SI (Fig S19(I)) and geometrical parameters are given in Table 2. In comparison with the simulated DC and DCu reduced state forms generated from Ferric-SFX, the geometrical parameters are quite similar, except that longer Fe-HisN distances, shorter Fe-water distances and shorter Fe-OOP are observed in *SimFerrous-DC/DCu* compared to Ferrous-DC/DCu. The spectral calculation on these DC and DCu systems, however, provide a better insight to ascertain the nature of the crosslink.

##### Comparing Ferrous-SC with *SimFerrous-SC*

Obtaining a structure of the ferrous form of cytochrome P460 proved highly challenging whether using chemical reduction or via electrons generated through X-ray exposure. Our studies provide the opportunity to compare whether a simulated reduced system (*SimFerrous-SC*) from an oxidised structure (**ferric SFX**) can provide results comparable with simulations performed on the reduced structure itself (Ferrous-SC). Comparison of the geometrical parameters around heme, *SimFerrous-SC* systems does provide similar lengthening of the Fe-Water distance and increase of the Fe-OOP movement as observed in **ferrous Cryo** structure. The MD results show more variations in the in *SimFerrous-SC* system due to the asymmetry in the two chains rather than the overall behaviour. MD trajectories reveal that Fe losses it’s water coordination from the 6^th^ coordination site for both systems. The lost water in turn forms H-bond interaction with pyrrole NA, chain A showing more prominence in both systems. This is also the case for the H-bond interaction with Asp102.

#### Evidence from calculations of the UV-Vis spectra

Spectral calculations of the McP460 systems revealed that the computed spectral shifts are relatively insensitive to the spin state of the ferrous and ferric forms. Therefore, the spectral shift observed experimentally is qualitatively reproduced irrespective of changes in the spin state on reduction. Although the optimised minima for each spin state indicates a complex relationship between geometry, spin multiplicity and energetics, this finding is consistent with spin states calculated for the DCu model in the ferric form and SC model in the ferrous form, in line with the assignments in the experimental structures.

We note that although we see differences in the spin energetics for chain A and chain B arising due to the conformation of the Asp102 residue, the energy differences between the spin states are small (less than ∼4.0 kcal/mol). These calculations altogether provide evidence for the presence of a double crosslinked species in the crystal. In transition metal complexes, multiple spin states may be involved in the reaction and can interconvert between each other through spin crossovers, known as Multiple-State reactivity.^63^ Due to the small energy differences observed between the individual spin states here, Multiple-State reactivity is possible. Characterizing these spin crossovers require computation of Minimum Energy Crossing Points (MECP) between spin states and is beyond the scope of the current paper.

The double-crosslink we observe in McP460 is reminiscent of that sometimes present in the P460 heme from HAO^62,64^. Indeed, HAO also harbours a crosslink between the meso carbon of its heme (albeit the opposite one, α-meso a.k.a CHC) and an amino acid sidechain, though from a tyrosine residue. A second crosslink is sometime observed between the tyrosine sidechain oxygen and the pyrrole carbon next to the α-meso carbon, forming a new 5-membered ring (as opposed to the herein reported new 6-membered-ring). However, the π bond in the HAO 5-membered ring is not part of the heme π-system, as it is orthogonal to it. On the other hand, the double bond on the lysine of McP460 is involved in the electronic transition making the Soret peak as revealed by the natural transition orbitals (Fig. S30 in SI). The difference between the Soret bands of McP460 (∼420 nm) and NeP460 (440 nm)^57^ was previously attributed to the iron-bound water of McP460^3^. The second lysine crosslink of McP460 is likely to be a bigger factor.

## Conclusions

We have described the first serial femtosecond crystallography structure of any cytochrome P460, achieving the third highest resolution room temperature structure from an XFEL in the PDB at the time of writing. As the structure of ferric McP460 is free of the effects of radiation induced chemistry due to the principle of diffraction before destruction and avoids the presence of any artefacts due to cryo it provided the best starting point for molecular simulation. The presence of a second cross link between the active site lysine residue and the heme was suggested by this structure with supporting room temperature multi-crystal structures supporting this concept and excluding the possibility that it could be caused by ageing of crystals. An anaerobic, freeze trapped structure of the ferrous form of the enzyme was also determined from a single-crystal with the oxidation state confirmed using single-crystal UV-visible spectroscopy.

QM/MM simulations of the ferric form of McP460 indicated that a double crosslink structure is stable and compatible with the available experimental data. Notably, distortion of the heme between Fe and pyrrole D ring nitrogen is observed in the optimised structures, in qualitative agreement with the crystallographic structure. QM/MM geometry optimisations on the ferrous form also reproduced key experimental structural features such as the interaction of water and iron at the 6^th^ coordination site. The QM/MM calculations of both the oxidised and reduced forms revealed a complex energetic landscape with several spin states accessible in both and sensitive to the local chemical environment of the heme. UV-vis spectroscopic simulations using TD-DFT suggest that the shift in absorption maximum observed experimentally between the ferric and ferrous forms is compatible with and supports a double crosslink model. For validation we also considered an in silico reduced model starting from the ferric crystal structure. This was also able to reproduce key features of the experimental data, indicating that in silico reduction is a valid approach to modelling the ferrous state in the case that only an oxidised experimental structure is available, and adding further support to the double crosslink proposal.

Intriguingly, no evidence of a second cross link was observed in previous crystal structures of a different cytochrome P460[NeP460; 2je3] from an ammonia oxidising organism which may be reflected in differing catalytic activity and chemical conditions within the organism. Notably the structure of NeP460 contained a hydroxyl modification to the heme group evident in low-dose structures but that disappeared in higher dose structures, a feature not present in McP460.

Our work highlights the power of combining multiple experimental and computational techniques to address the structure and mechanism of an enzyme. Electron density features that may be ambiguous or unclear even at very high resolutions may be investigated and validated using simulation while the simulations themselves benefit from starting structures that are free of experimental artefacts such as radiation damage and changes due to cryo cooling. In addition to comparison of coordinates and bonds, the ability to predict experimental structures and to measure them from the same crystals used for structure determination allows for further cross validation. Future work will focus on applying this combined approach to the ligand bound states of cytochrome P460 and site-directed mutagenesis of protein residues proposed to be relevant to the catalytic mechanism.

## Acknowledgements

This work was supported by BBSRC grant awards BB/V016660/1, BB/V01577X/1 and BB/V016768/1, and BBSRC Japan Partnering Awards BB/R021015/1 and BB/X01844X/1. XFEL experiments were performed at BL2 EH3 of SACLA with the approval of the Japan Synchrotron Radiation Research Institute (JASRI); proposal 2022B8041. Travel support from the UK XFEL Hub at the Diamond Light Source is gratefully acknowledged. We acknowledge use of the Crystallization facility at the Research Complex at Harwell (Dr Halina Mikolajek) and of Diamond beamlines VMXi and I24 under proposal nr27313. We acknowledge the support of Drs Juan Sanchez Weatherby and James Sandy. Andrey Lebedev is thanked for his help preparing the HX2 heme restraints. We thank Dr Jos Kamps and Cicely Tam for their help with anaerobic work. Professors Emma Raven and Jonathan Clayden are thanked for helpful discussions. This work made use of computing resources provided by STFC Scientific Computing’s SCARF cluster, Bristol ACRC (http://www.bristol.ac.uk/acrc/) and the ARCHER2 UK National Supercomputing Service (https://www.archer2.ac.uk). H.R.A. was supported by a PhD studentship from the University of Essex.

