## Supporting information for "Double crossed? Structural and computational studies of an unusual crosslinked heme in *Methylococcus capsulatus* cytochrome P460"

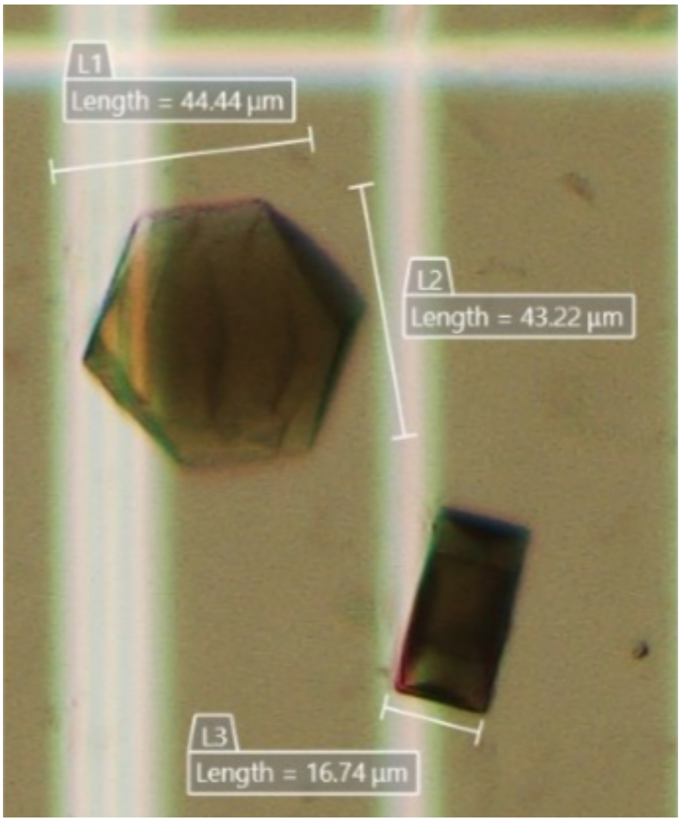

**Fig. S1.** Batch crystals from SFX experiments. L1 for length, L2 for width and L3 for thickness

A)

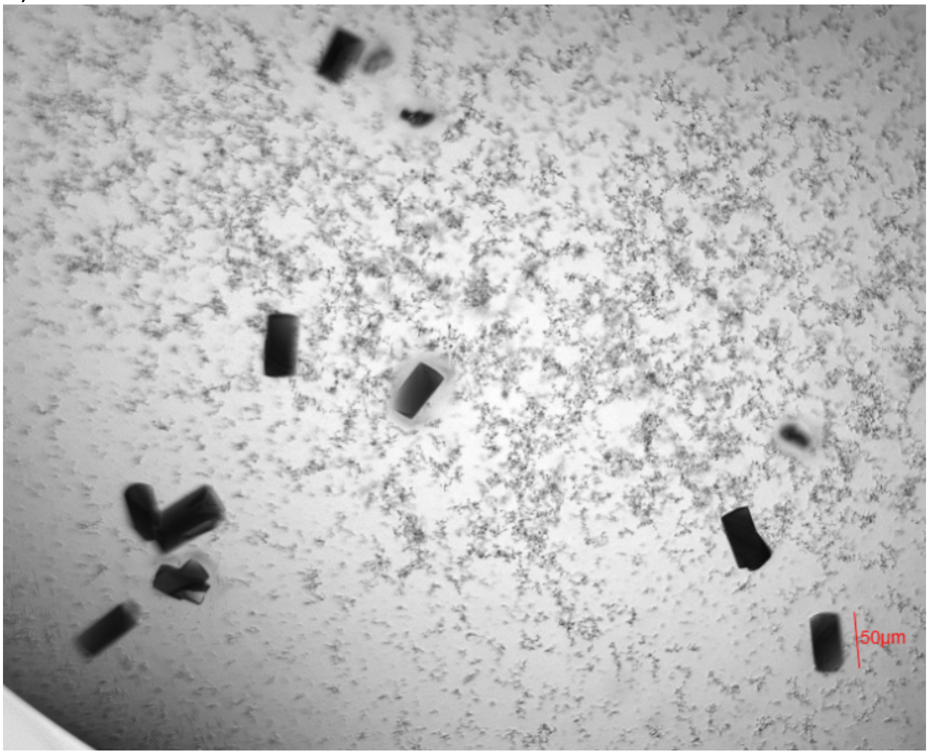


B)

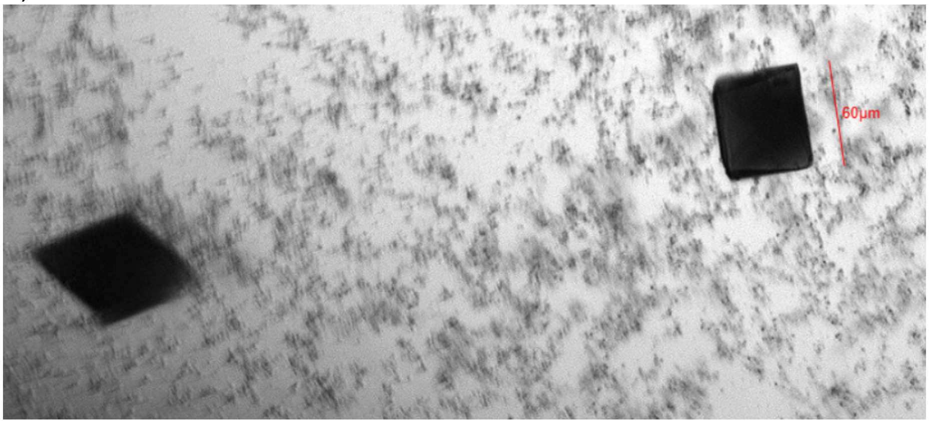


**Fig. S2.** Crystals used for A) RT fresh and B) RT aged structures

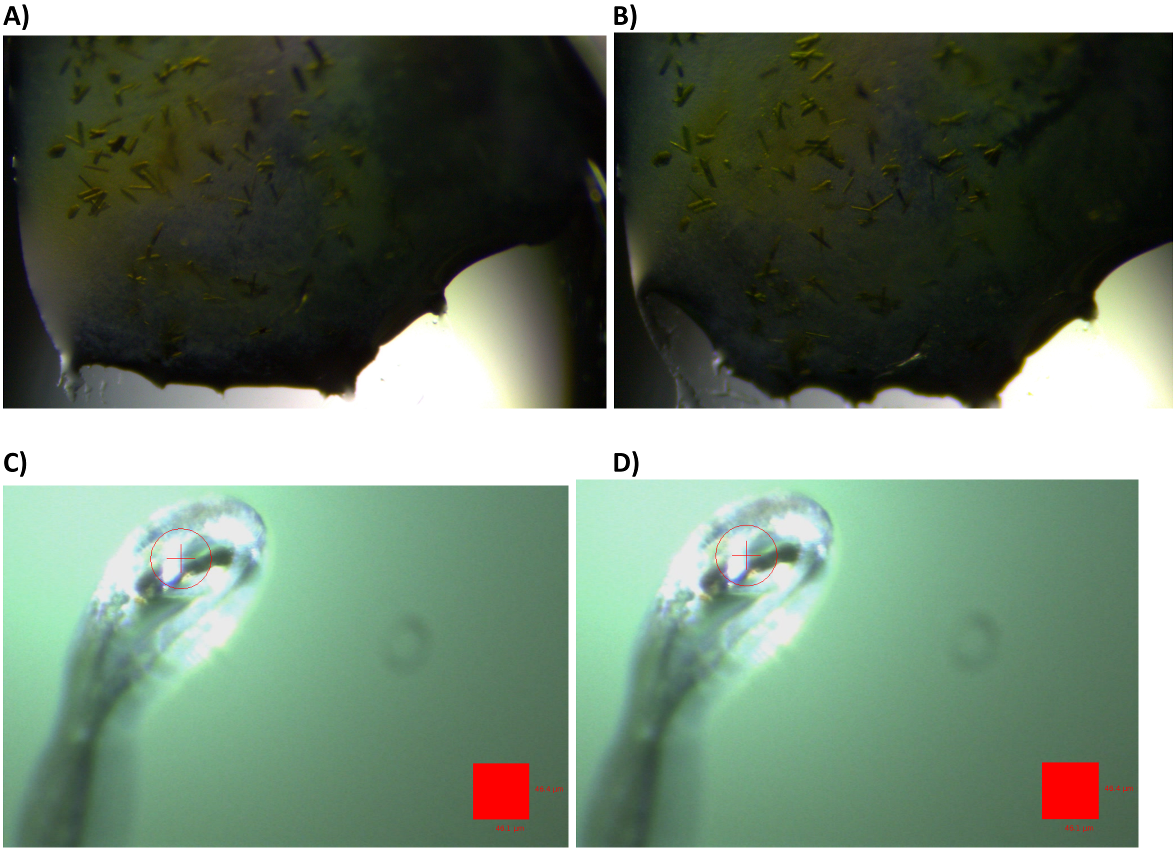


**Fig. S3.** McP460 crystals (200×35×20 µm) in cryoprotectant A) before and B) after addition of sodium dithionite reductant. C) Cryocooled crystal before and D) after collection of the first X-ray diffraction dataset (illuminated by the white light of the microspectrophotometer)

**Table S1** - Heme site parameters forP460 crystal structures (chain A/chain B)

| **Structure** | **Ferric SFX [9hs4]** | **VMXi 1 fresh [9hs9]** | **VMXi2 aged [9hs6]** | **Ferrous (100K) [9hrk]** | **6hiu (100K)** | **NeP460 (2je3)** (100K) |
|---|---|---|---|---|---|---|
| Resolution (Å) | 1.28 | 1.66 | 1.77 | 1.33 | 1.36 | 1.80 |
| Fe–His N (Å) | 2.14/2.12 | 2.11/2.05 | 2.11/2.06 | 2.10/2.10 | 2.13/2.12 | 2.16 |
| Fe–water (Å) | 2.11/2.11 | 2.15/2.11 | 2.06/2.09 | -3.33/3.20 | 2.32/2.37 | 2.62 to Pi |
| Fe-PyrNA (Å) | (2.03,1.99, 2.00,2.00) / (2.02,1.99, 2.01, 2.01) | (2.06, 2.00, 2.03,2.01) / (2.07,1.98, 2.04, 2.98) | (2.04,1.99, 2.04,2.02) / (2.09,2.01, 2.04,2.0) | (2.10,2.09, 2.06,2.12) / (2.10,2.05, 2.00,2.05) | (2.07,2.02,2 .07,2.04) / (2.08,2.01, 2.03,2.08) | 2.12, 2.10, 2.12, 2.12 |
| Lys–CHA (Å) | 1.46/1.50 | 1.40/1.38 | 1.40/1.39 | 1.44/1.41 | (1.35,1.35) / (1.33, 1.37) | 1.63 |
| LysCD-C2A | 1.94/2.11 | 1.58/1.58 | 1.55/1.56 | 3.72/3.71 | (2.17,3.92) / (2.26,3.83) | 3.70 |
| C3A-CMA | 1.43/1.44 | 1.33/1.33 | 1.35/1.33 | 1.53/1.50 | 1.48/ | 1.55 |
| Fe -OOP (Å) | 0.06/0.06 | 0.11/0.13 | 0.08/0.06 | 0.35/0.33 | 0.11/0.10 | 0.24 |

**S.1 Charge determination**

P460 has a unique heme C unit where an additional covalent bond is formed between N atom of Lys78 and meso-CG (CHA) of heme C (Scheme 1, main manuscript). This covalent modification required calculation of the charges of the heme C unit with this unique covalent bond and the coordinated residues. CHARMM forcefield (FF) uses groups to define neutral units within molecules. For the available heme B cofactor all the side chains (propionate, vinyl and methyl) of Heme B unit are defined as groups and sums up to 0.0. To minimize the charge manipulation and fit it with the rest of the charges provided by CHARMM FF the following steps were performed: For the charge manipulation of the modified heme C cofactor in P460, a cluster model consisting of the core heme C, the covalently linked Cys140 and Cys143, the proximal His ligand and the crosslinked Lys78 was considered. All these amino acid residues were truncated at the neutral CA-CB. See Fig. S4 – the brown stars indicate where the side chains of the Heme were cut off and the purple stars indicate the terminal atoms of the linked active site residues. Geometry optimization was carried out in NWCHEM, the CB atoms of His144, Cys140, Cys143 were fixed to their crystallographic coordinates. For Lys78, to keep as close as possible to crystal structure, the CB and CG were fixed. DFT functional B3LYP with D3 dispersion correction was used. The basis set of 631G* was used for all atoms. The geometries were optimized in the spin states of doublet (M = 2), quartet (M = 4) and sextet (M = 6) for ferric state and singlet (M = 1), triplet (M = 3), and quintet (M = 5) for ferric state, respectively. The electrostatic potential (ESP) scheme within NWCHEM was used to derive charges. The lowest energy optimized state was taken for charge determination – which was quartet for ferric and triplet for ferrous states, respectively. It was followed by addition of constraints to the ESP charges so assign equivalency to chemically similar groups and also to minimize disparity with CHARMM FF charges when this cofactor is added within the protein for simulation. The constrains used were:

1. Charges were set to zero to the H atoms that was used to replace the side chains of heme (Figure S1, brown stars)
2. Charges were set to zero to the H atoms which were added for replace the cut made between Ca-Cb atoms for the amino acids (pink stars)
3. Additionally, the charges from CB to CD were summed to zero for the Lys78 side chain (pink stars) to keep parity with CHARMM FF for this amino acid.
4. In Heme C, Cys covalently links to the vinyl (-C=$CH_2$) group of Heme B forming -CH(SCys)-$CH_3$. The charges on this generated -$CH_3$ group were summed to zero (pink stars) and the charge on H-atoms were made equivalent.
5. The charges on His residue were also summed to zero following the His links to Heme B groups in CHARMM FF which remains unaffected in charges.
6. All equivalent atoms were made to have equal charges.
7. For the porphyrin core, the two pyrrole rings next to the methylene bridge (CHA) where Lys78 forms the cross link were kept equivalent to one another. The other two pyrroles were also kept equivalent to each other.
8. Except for the methylene bridge (CHA) where the Lys cross links, the other 3 were made equivalent.
9. Double crosslink (DC) and double crosslink with unsaturated Lys (DCu) were generated by removing the corresponding H-atoms and adding the charges of the removed H- atoms to the C-atom it was bonded to.

The charges so derived are provided in Table S1. The charges that were different in case of DC and DCu are given in in Table S1.

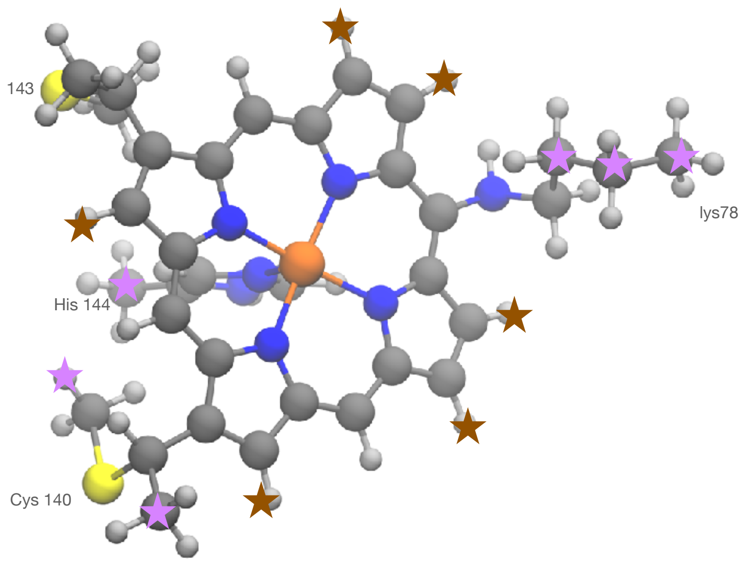


**Fig. S4.** The cluster model used for charge determination. The brown stars indicate where the side chains of the heme were cut off and the purple stars indicate the terminal atoms of the linked active site residues.

**Table S2**. Charges used for ferric and Ferrous states for the heme C unit with the covalent links to Lys78, Cys140 and Cys143 and coordinated His residue. The full set of charges for single crosslink are provided. For double cross link and double cross link with unsaturated Lys the atoms that had different charges are only provided.

| **Residue** | **Atom names** | **Single crosslink (SC)** | | **Double crosslink (DC)** | | **Double crosslink with unsaturated Lys (DCu)** | |
|---|---|---|---|---|---|---|---|
| | | **Ferric** | **Ferrous** | **Ferric** | **Ferrous** | **Ferric** | **Ferrous** |
| Lys-78 | CD* | NC | NC | -0.09 | -0.09 | 0.00 | 0.00 |
| | HD | NC | NC | 0.09 | 0.09 | - | - |
| | CE* | 0.08 | 0.16 | | | 0.23 | 0.24 |
| | HE1 | 0.15 | 0.08 | | | 0.15 | 0.08 |
| | HE2 | 0.15 | 0.08 | | | - | - |
| | NZ | -0.85 | -0.83 | | | | |
| | HZ | 0.40 | 0.41 | | | | |
| Cys-140 | CB | 0.16 | 0.23 | | | | |
| | HB1 | -0.05 | -0.07 | | | | |
| | HB2 | -0.05 | -0.07 | | | | |
| | SG | -0.29 | -0.37 | | | | |
| Cys-143 | CB | 0.16 | 0.23 | | | | |
| | HB1 | -0.05 | -0.07 | | | | |
| | HB2 | -0.05 | -0.07 | | | | |
| | SG | -0.29 | -0.37 | | | | |
| His-144 | CB | 0.09 | -0.04 | | | | |
| | HB1 | 0.06 | 0.08 | | | | |
| | HB2 | 0.06 | 0.08 | | | | |
| | ND1 | -0.12 | -0.14 | | | | |

| | | | | | | | |
|---|---|---|---|---|---|---|---|
| | HD1 | 0.31 | 0.28 | | | | |
| | CG | -0.10 | 0.02 | | | | |
| | CE1 | -0.02 | -0.02 | | | | |
| | HE1 | 0.18 | 0.16 | | | | |
| | NE2 | -0.36 | -0.28 | | | | |
| | CD2 | 0.18 | -0.10 | | | | |
| | HD2 | 0.07 | 0.19 | | | | |
| Heme C | FE | 0.51 | 0.20 | | | | |
| | NA | -0.20 | -0.19 | | | | |
| | NB | -0.27 | -0.10 | | | | |
| | NC | -0.27 | -0.10 | | | | |
| | ND | -0.20 | -0.19 | | | | |
| | C1A | 0.24 | 0.26 | | | | |
| | C2A | -0.12 | -0.17 | | | | |
| | C3A | -0.12 | -0.17 | | | | |
| | C4A | 0.24 | 0.26 | | | | |
| | C1B | 0.34 | 0.18 | | | | |
| | C2B | -0.17 | -0.16 | | | | |
| | C3B | -0.17 | -0.16 | | | | |
| | C4B | 0.34 | 0.18 | | | | |
| | C1C | 0.34 | 0.18 | | | | |
| | C2C | -0.17 | -0.16 | | | | |
| | C3C | -0.17 | -0.16 | | | | |
| | C4C | 0.34 | 0.18 | | | | |
| | C1D | 0.24 | 0.26 | | | | |
| | C2D | -0.12 | -0.17 | | | | |
| | C3D | -0.12 | -0.17 | | | | |
| | C4D | 0.24 | 0.26 | | | | |
| | CHA | 0.28 | 0.16 | | | | |
| | CHB | -0.43 | -0.36 | | | | |
| | HB | 0.22 | 0.17 | | | | |
| | CHC | -0.43 | -0.36 | | | | |
| | HC | 0.22 | 0.17 | | | | |
| | CHD | -0.43 | -0.36 | | | | |
| | HD | 0.22 | 0.17 | | | | |
| | CMA* | NC | NC | -0.18 | -0.18 | -0.18 | -0.18 |
| | HMA1 | NC | NC | 0.09 | 0.09 | 0.09 | 0.09 |
| | HMA2 | NC | NC | 0.09 | 0.09 | 0.09 | 0.09 |
| Cys-140 link | CAB | 0.52 | 0.61 | | | | |
| | HAB | -0.08 | -0.10 | | | | |

| | | | |
|---|---|---|---|
| | CBB | -0.37 | -0.39 |
| | HBB1 | 0.11 | 0.09 |
| | HBB2 | 0.11 | 0.09 |
| | HBB3 | 0.11 | 0.09 |
| Cys-143 link | CAC | 0.52 | 0.61 |
| | HAC | -0.08 | -0.10 |
| | CBC | -0.37 | -0.39 |
| | HBC1 | 0.11 | 0.09 |
| | HBC2 | 0.11 | 0.09 |
| | HBC3 | 0.11 | 0.09 |

NC- this group was not considered in the cluster for charge calculation, CHARMM FF charges were used directly. *DC and DCu involved removal of a H atom bonded to Heme-CMA and Lys-CD, so the charge on H atom that was removed was added to the charge of the C atom to which it was bonded. From DC to DCu involved an unsaturated Lys CD-CE bond, hence required removal of further H-atoms bonded to CD and CE of lysine. So, the charges of these H atoms removed were added to the remaining charge on the CE and CD atoms.

**S.2 Modelling the crosslinks**

To model the cross-link to represent both double and single cross-links, partial optimization of the heme unit around the crosslink was performed. Gas phase cluster calculations were performed where all the heme atoms, except those involved in the crosslink and their two immediate neighbouring atoms were fixed (Fig S5). Both oxidized and reduced forms of Fe were calculated; spin states considered were M=2, 4 and 6 for ferric state and M=1, 3 and 5 for ferrous state. The resulting optimised geometries from all spin states were similar for both oxidised ferric and reduced ferrous systems. The key distance: LysN-CHA (1.36 ± 0.02 Å) remains invariant among all the optimised structures for both SC and DC systems. In addition, the CD-C2A (1.55 Å) and C3A-CMA (1.35 Å) distances for DC are also invariant across all structures. The angles around the linking of NZ of Lys78 to heme averages to: NZ-CHA-C1A (115.69 ± 0.31º and 116.60 ± 0.59 º, for SC and DC, respectively), NZ-CHA-C4D (120.58 ± 0.34º and 121.86 ± 0.32º for SC and DC, respectively). The angles around the second CD-C2A link: CD-C2A-C1A and CD-C2A-C3A for DC averages to 99.24 ± 0.46º and 113.20 ± 0.20º, respectively. The resulting optimised SC and DC heme C unit in the lowest spin state were aligned to the crystal heme C unit of the **ferric SFX** structure. The alignment showed good agreement at the junctions of CA – CB bond to Lys78, Cys140, Cys143 and His144. The heme unit from SFX structure was replaced by these optimised SC and DC heme unit - thus providing starting structure for both SC and DC (Fig S6).

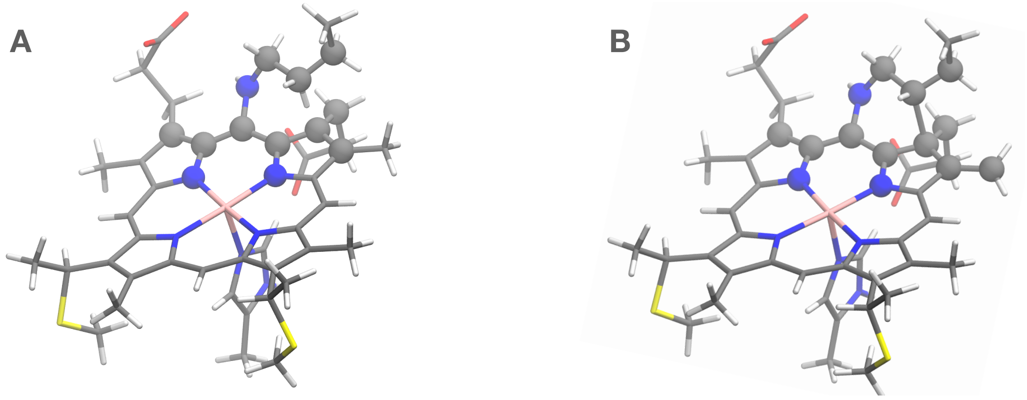


**Fig. S5.** QM models for generating A) SC and B) DC heme units. The atoms in spheres were optimized only, the rest were fixed

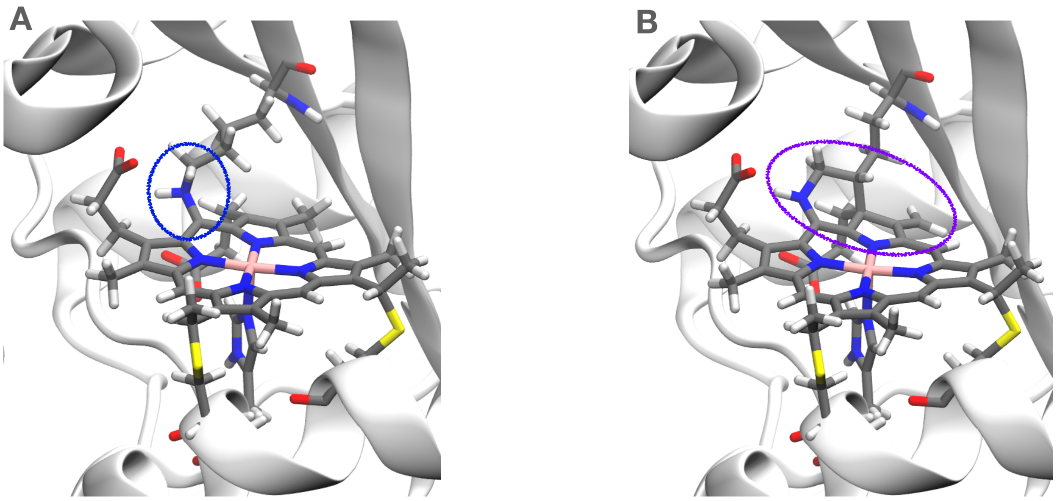


**Fig. S6.** Optimised QM models A) SC and B) DC heme units with crosslinks aligned within the active site of ferric SFX providing the initial structures for Ferric-SC and Ferric-DC.

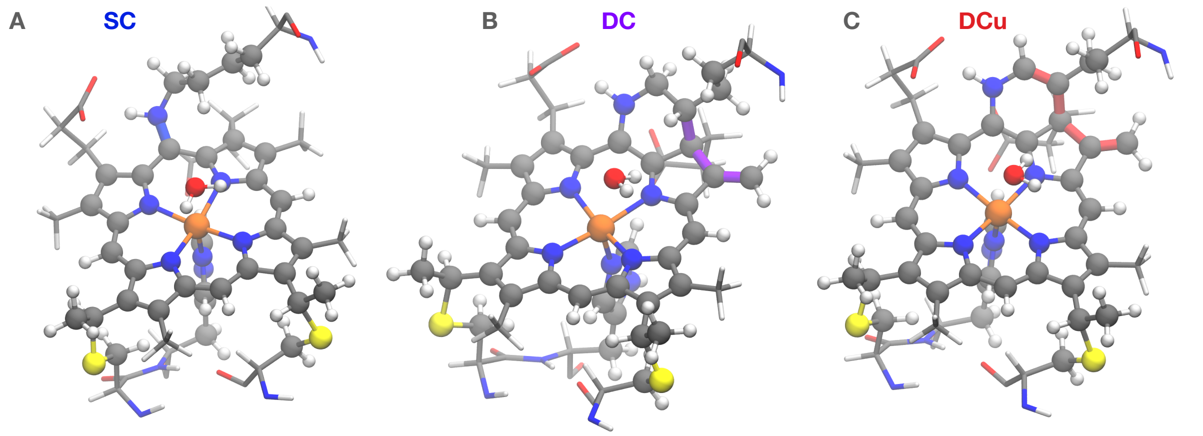


**Fig. S7.** Atoms included in QM regions are shown in spheres for A) SC, B) DC and C) DCu. The changes that occurs due to crosslink are shown in blue for SC, in violet for DC and in red for DCu.

**Table S3A.** Absolute and relative energies of QM/MM optimised Ferric-SC/DC/DCu systems in all three spin states.

| | **Chain A (Hartree)** | **Relative energies wrt lowest energy spin state of same chain (kcal/mol)** | **Chain B (Hartree)** | **Relative energies wrt lowest energy spin state of same chain (kcal/mol)** | **Relative energies wrt lowest energy spin state of chain A (kcal/mol)** |
|---|---|---|---|---|---|
| **Ferric-SC** | | | | | |
| **M=2** | -3840.45 | 7.14 | -3840.46 | 5.27 | 2.77 |
| **M=4** | -3840.46 | 0.65 | **-3840.47** | **0.00** | **-2.50** |
| **M=6** | **-3840.46** | **0.00** | -3840.46 | 4.94 | 2.43 |
| **Ferric-DC** | | | | | |
| **M=2** | -3878.43 | 0.88 | **-3878.44** | **0.00** | **-1.10** |
| **M=4** | -3878.42 | 8.85 | -3878.43 | 4.14 | 3.04 |
| **M=6** | **-3878.44** | **0.00** | -3878.43 | 5.31 | 4.21 |
| **Ferric-DCu** | | | | | |

| | | | | | |
|---|---|---|---|---|---|
| **M=2** | **-3877.11** | **0.00** | -3877.11 | 2.25 | 0.12 |
| **M=4** | -3877.11 | 0.99 | **-3877.12** | **0.00** | **-2.13** |
| **M=6** | -3877.11 | 2.70 | -3877.11 | 2.12 | -0.02 |

**Table S3B.** Absolute and relative energies of QM/MM optimised Ferrous-SC/DC/DCu systems in all three spin states.

| | **Chain A (Hartree)** | **Relative energies wrt lowest energy spin state of same chain (kcal/mol)** | **Chain B (Hartree)** | **Relative energies wrt lowest energy spin state of same chain (kcal/mol)** |
|---|---|---|---|---|
| **Ferrous-SC** | | | | |
| **M=1** | -3839.93 | 8.32 | -3839.98 | 9.86 |
| **M=3** | -3839.94 | 2.14 | -3839.99 | 2.20 |
| **M=5** | **-3839.95** | **0.00** | **-3839.99** | **0.00** |
| **Ferrous-DC** | | | | |
| **M=1** | -3877.84 | 15.77 | -3877.91 | 7.19 |
| **M=3** | -3877.85 | 9.87 | -3877.92 | 1.68 |
| **M=5** | **-3877.87** | **0.00** | **-3877.92** | **0.00** |
| **Ferrous-DCu** | | | | |
| **M=1** | -3876.49 | 4.64 | -3876.55 | 9.01 |
| **M=3** | -3876.49 | 2.28 | -3876.56 | 2.53 |
| **M=5** | **-3876.50** | **0.00** | **-3876.56** | **0.00** |

**Table S3C.** Absolute and relative energies of QM/MM optimised *SimFerrous-SC/DC/DCu* systems in all three spin states.

| | **Chain A (Hartree)** | **Relative energies wrt lowest energy spin state of same chain (kcal/mol)** | **Chain B (Hartree)** | **Relative energies wrt lowest energy spin state of same chain (kcal/mol)** |
|---|---|---|---|---|
| ***SimFerrous-SC*** | | | | |
| **M=1** | -3840.34 | 5.44 | -3840.34 | 1.99 |
| **M=3** | -3840.34 | 1.51 | **-3840.35** | **0.00** |
| **M=5** | **-3840.35** | **0.00** | -3840.35 | 0.20 |
| ***SimFerrous-DC*** | | | | |
| **M=1** | -3878.32 | 1.56 | **-3878.31** | **0.00** |
| **M=3** | -3878.32 | 1.65 | -3878.31 | 2.00 |
| **M=5** | **-3878.32** | **0.00** | -3878.30 | 4.66 |
| ***SimFerrous-DCu*** | | | | |
| **M=1** | -3876.99 | 2.70 | **-3876.99** | **0.00** |
| **M=3** | -3876.99 | 3.98 | -3876.97 | 9.27 |
| **M=5** | **-3876.99** | **0.00** | -3876.97 | 10.92 |

In tables S3A-C the two chains reported are treated as individual systems.

**Table S4.** Heme C site parameters for McP460 structures, including: the distance between proximal His143 N and Fe (Fe-HisN), the distance between the coordinated water to Fe (Fe-water), Fe to porphyrin N of all pyrrole rings (Fe-PyrN), the distance between Lys78 and CHA of heme (LysN-CHA), the distance between CD of Lys78 and C2A of heme in DC systems (LysCD-C2A), the distance between heme C3A and CMA distance (C3A-CMA) that undergoes exocyclic modification in DC systems and Fe out of plane motion (Fe-OOP).

**A) Ferric-SC/DC/DCu in all spin states and both chains**

| | Fe-His N (Å) | | Fe-water N (Å) | | Fe PyrN (Å): NA, NB, NC and ND | | LysN-CHA (Å) | | LysCD-C2A (Å) | | C3A-CMA (Å) | | Fe-OOP (Å)* | |
|---|---|---|---|---|---|---|---|---|---|---|---|---|---|---|
| **Chain** | **A** | **B** | **A** | **B** | **A** | **B** | **A** | **B** | **A** | **B** | **A** | **B** | **A** | **B** |
| **Ferric-SC** | | | | | | | | | | | | | | |
| **M = 2** | 1.96 | 2.00 | 2.16 | 2.30 | 1.99, 2.03, 2.05, 2.00 | 2.01, 2.01, 2.04, 2.04 | 1.46 | 1.41 | - | - | 1.47 | 1.47 | 0.12 | 0.08 |
| **M = 4** | 2.15 | 2.18 | 2.54 | 2.36 | 1.99, 2.03, 2.04, 2.01 | 2.01, 2.01, 2.02, 2.02 | 1.46 | 1.41 | - | - | 1.47 | 1.47 | 0.14 | 0.05 |
| **M = 6** | 2.09 | 2.13 | 2.58 | 2.32 | 2.08, 2.07, 2.09, 2.08 | 2.07, 2.06, 2.08, 2.08 | 1.46 | 1.41 | - | - | 1.47 | 1.47 | 0.24 | 0.10 |
| **Ferric-DC** | | | | | | | | | | | | | | |
| **M = 2** | 1.92 | 1.95 | 2.35 | 2.29 | 2.09, 2.00, 2.01, 2.01 | 2.09, 2.01, 2.02, 2.02 | 1.36 | 1.36 | 1.57 | 1.57 | 1.35 | 1.35 | 0.12 | 0.09 |
| **M = 4** | 2.11 | 2.17 | 2.75 | 2.38 | 2.07, 2.01, 2.01, 2.01 | 2.11, 2.00, 2.00, 2.01 | 1.36 | 1.36 | 1.57 | 1.57 | 1.35 | 1.35 | 0.14 | 0.06 |
| **M = 6** | 2.11 | 2.17 | 2.68 | 2.37 | 2.22, 2.07, 2.06, 2.09 | 2.20, 2.06, 2.05, 2.09 | 1.34 | 1.35 | 1.57 | 1.56 | 1.35 | 1.35 | 0.20 | 0.09 |
| **Ferric-DCu** | | | | | | | | | | | | | | |
| **M = 2** | 1.93 | 1.96 | 2.33 | 2.28 | 2.10, 2.00, 2.01, 2.02 | 2.11, 2.01, 2.03, 2.03 | 1.38 | 1.38 | 1.54 | 1.54 | 1.35 | 1.35 | 0.12 | 0.09 |
| **M = 4** | 2.11 | 2.16 | 2.51 | 2.30 | 2.12, 2.00, 2.00, 2.01 | 2.12, 2.00, 2.01, 2.01 | 1.38 | 1.38 | 1.54 | 1.54 | 1.35 | 1.35 | 0.13 | 0.07 |
| **M = 6** | 2.11 | 2.14 | 2.49 | 2.27 | 2.24, 2.06, 2.04, 2.09 | 2.22, 2.06, 2.03, 2.09 | 1.37 | 1.37 | 1.53 | 1.53 | 1.35 | 1.35 | 0.19 | 0.11 |

**B) Ferrous-SC/DC/DCu in all spin states and both chains**

| | Fe-His N (Å) | | Fe-water N (Å) | | Fe PyrN (Å): NA, NB, NC and ND | | LysN-CHA (Å) | | LysCD-C2A (Å) | | C3A-CMA (Å) | | Fe-OOP (Å)* | |
|---|---|---|---|---|---|---|---|---|---|---|---|---|---|---|
| **Chain** | **A** | **B** | **A** | **B** | **A** | **B** | **A** | **B** | **A** | **B** | **A** | **B** | **A** | **B** |
| **Ferrous-SC** | | | | | | | | | | | | | | |
| **M = 1** | 1.97 | 1.96 | 3.11 | 3.51 | 2.04, 2.02, 2.03, 2.02 | 2.02, 2.02, 2.02, 2.05 | 1.36 | 1.36 | - | - | 1.47 | 1.47 | 0.15 | 0.17 |
| **M = 3** | 2.24 | 2.20 | 3.22 | 3.53 | 2.04, 2.01, 2.04, 2.03 | 2.03, 2.03, 2.03, 2.04 | 1.35 | 1.36 | - | - | 1.47 | 1.47 | 0.13 | 0.16 |
| **M = 5** | 2.11 | 2.08 | 3.38 | 3.68 | 2.14, 2.11, 2.12, 2.10 | 2.12, 2.12, 2.10, 2.14 | 1.35 | 1.35 | - | - | 1.47 | 1.47 | 0.40 | 0.43 |
| **Ferrous-DC** | | | | | | | | | | | | | | |
| **M = 1** | 1.94 | 1.94 | 3.26 | 3.37 | 2.02, 2.03, 2.03, 2.03 | 2.02, 2.03, 2.02, 2.04 | 1.36 | 1.36 | 1.57 | 1.57 | 1.35 | 1.35 | 0.16 | 0.16 |

| | | | | | | | | | | | | | | |
|---|---|---|---|---|---|---|---|---|---|---|---|---|---|---|
| **M = 3** | 2.17 | 2.15 | 3.31 | 3.48 | 2.02, 2.04, 2.03, 2.03 | 2.03, 2.04, 2.02, 2.04 | 1.36 | 1.36 | 1.57 | 1.57 | 1.35 | 1.35 | 0.15 | 0.17 |
| **M = 5** | 2.11 | 2.10 | 3.26 | 3.56 | 2.15, 2.08, 2.11, 2.10 | 2.13, 2.09, 2.11, 2.11 | 1.37 | 1.36 | 1.57 | 1.57 | 1.35 | 1.35 | 0.30 | 0.31 |
| **Ferrous-DCu** | | | | | | | | | | | | | | |
| **M = 1** | 2.01 | 1.95 | 2.28 | 3.37 | 2.04, 2.02, 2.05, 2.05 | 2.04, 2.03, 2.02, 2.04 | 1.39 | 1.38 | 1.54 | 1.54 | 1.35 | 1.35 | 0.06 | 0.16 |
| **M = 3** | 2.19 | 2.16 | 3.14 | 3.46 | 2.04, 2.03, 2.04,2.04 | 2.05, 2.04, 2.03, 2.04 | 1.38 | 1.38 | 1.54 | 1.54 | 1.35 | 1.35 | 0.13 | 1.17 |
| **M = 5** | 2.13 | 2.10 | 3.21 | 3.56 | 2.14, 2.07, 2.12, 2.09 | 2.14, 2.09, 2.12, 2.10 | 1.38 | 1.38 | 1.54 | 1.54 | 1.35 | 1.35 | 0.26 | 0.31 |

#### C) *SimFerrous-SC/DC/DCu* in all spin states and both chains

| | **Fe-His N (Å)** | | **Fe-water N (Å)** | | **Fe PyrN (Å): NA, NB, NC and ND** | | **LysN-CHA (Å)** | | **LysCD-C2A (Å)** | | **C3A-CMA (Å)** | | **Fe-OOP (Å)*** | |
|---|---|---|---|---|---|---|---|---|---|---|---|---|---|---|
| **Chain** | **A** | **B** | **A** | **B** | **A** | **B** | **A** | **B** | **A** | **B** | **A** | **B** | **A** | **B** |
| ***SimFerrous-SC*** | | | | | | | | | | | | | | |
| **M = 1** | 2.03 | 2.00 | 2.27 | 2.33 | 2.00, 2.05, 2.07, 2.03 | 2.04, 2.04, 2.04, 2.03 | 1.41 | 1.41 | - | - | 1.47 | 1.48 | 0.09 | 0.09 |
| **M = 3** | 2.23 | 2.24 | 3.28 | 3.05 | 2.01, 2.05, 2.05, 2.03 | 2.04, 2.04, 2.03, 2.04 | 1.41 | 1.41 | - | - | 1.47 | 1.47 | 0.14 | 0.13 |
| **M = 5** | 2.16 | 2.08 | 3.30 | 2.98 | 2.08, 2.11, 2.14, 2.10 | 2.13, 2.11, 2.12, 2.17 | 1.41 | 1.41 | - | - | 1.47 | 1.47 | 0.29 | 0.44 |
| ***SimFerrous-DC*** | | | | | | | | | | | | | | |
| **M = 1** | 1.99 | 1.97 | 2.28 | 2.30 | 2.03, 2.04, 2.06, 2.05 | 2.04, 2.07, 2.05, 2.03 | 1.37 | 1.37 | 1.57 | 1.57 | 1.35 | 1.35 | 0.09 | 0.1 |
| **M = 3** | 2.16 | 2.16 | 3.32 | 3.25 | 2.04, 2.05, 2.02, 2.04 | 2.05, 2.04, 2.02, 2.04 | 1.37 | 1.37 | 1.57 | 1.57 | 1.35 | 1.35 | 0.16 | 0.16 |
| **M = 5** | 2.15 | 2.05 | 2.61 | 2.94 | 2.11, 2.09, 2.14, 2.10 | 2.19, 2.09, 2.10, 2.16 | 1.37 | 1.37 | 1.56 | 1.57 | 1.35 | 1.35 | 0.21 | 0.46 |
| ***SimFerrous-DCu*** | | | | | | | | | | | | | | |
| **M = 1** | 2.00 | 1.99 | 2.27 | 2.26 | 2.05, 2.04, 2 .05, 2.06 | 2.05, 2.04, 2.06, 2.06 | 1.39 | 1.39 | 1.54 | 1.54 | 1.35 | 1.35 | 0.09 | 0.09 |
| **M = 3** | 2.17 | 2.18 | 3.33 | 2.96 | 2.05, 2.04, 2.03, 2.04 | 2.06, 2.03, 2.02, 2.05 | 1.39 | 1.39 | 1.54 | 1.54 | 1.35 | 1.35 | 0.16 | 0.13 |
| **M = 5** | 2.16 | 2.07 | 2.50 | 2.87 | 2.12, 2.09, 2.13, 2.10 | 2.18, 2.06, 2.10, 2.14 | 1.39 | 1.39 | 1.53 | 1.54 | 1.35 | 1.35 | 0.20 | 0.36 |

**Table S5.** Cluster calculations of Ferric-DC system without any environment

| | **Chain A (Hartree)** | **Relative energies wrt lowest energy spin state of chain A (kcal/mol)** | **Chain B (Hartree)** | **Relative energies wrt lowest energy spin state of chain A (kcal/mol)** |
|---|---|---|---|---|
| **Ferric-DC without environment** | | | | |
| **M=2** | -3875.65 | 2.85 | -3875.65 | 2.66 |
| **M=4** | **-3875.65** | **0.00** | **-3875.65** | **0.00** |
| **M=6** | -3875.64 | 5.65 | -3875.64 | 5.68 |

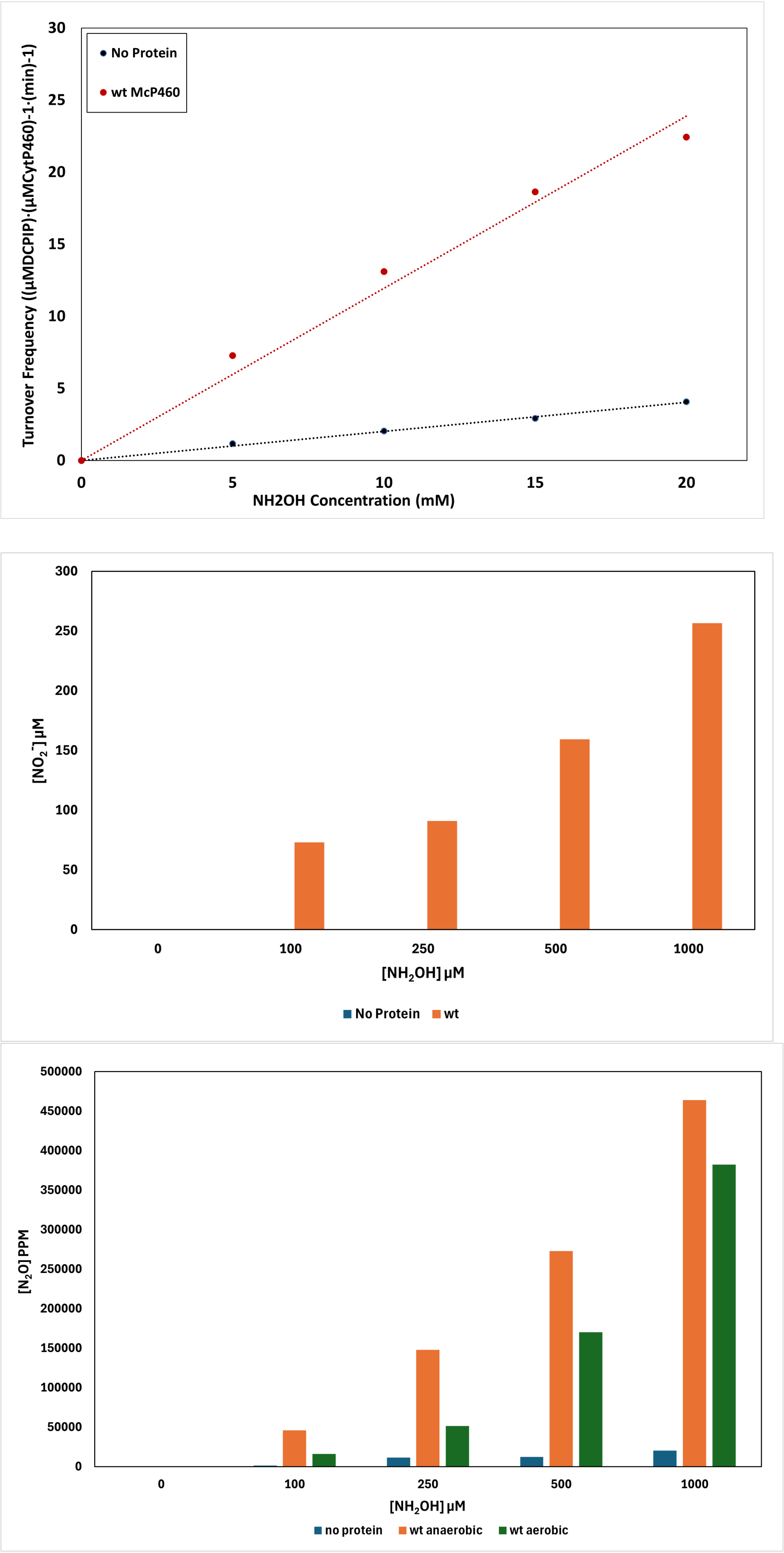


**Fig. S8**: enzyme activity data for the oxidation of hydroxylamine by McP460 (top panel); production of nitrite (middle panel); production of Nitrous oxide under aerobic and anaerobic conditions

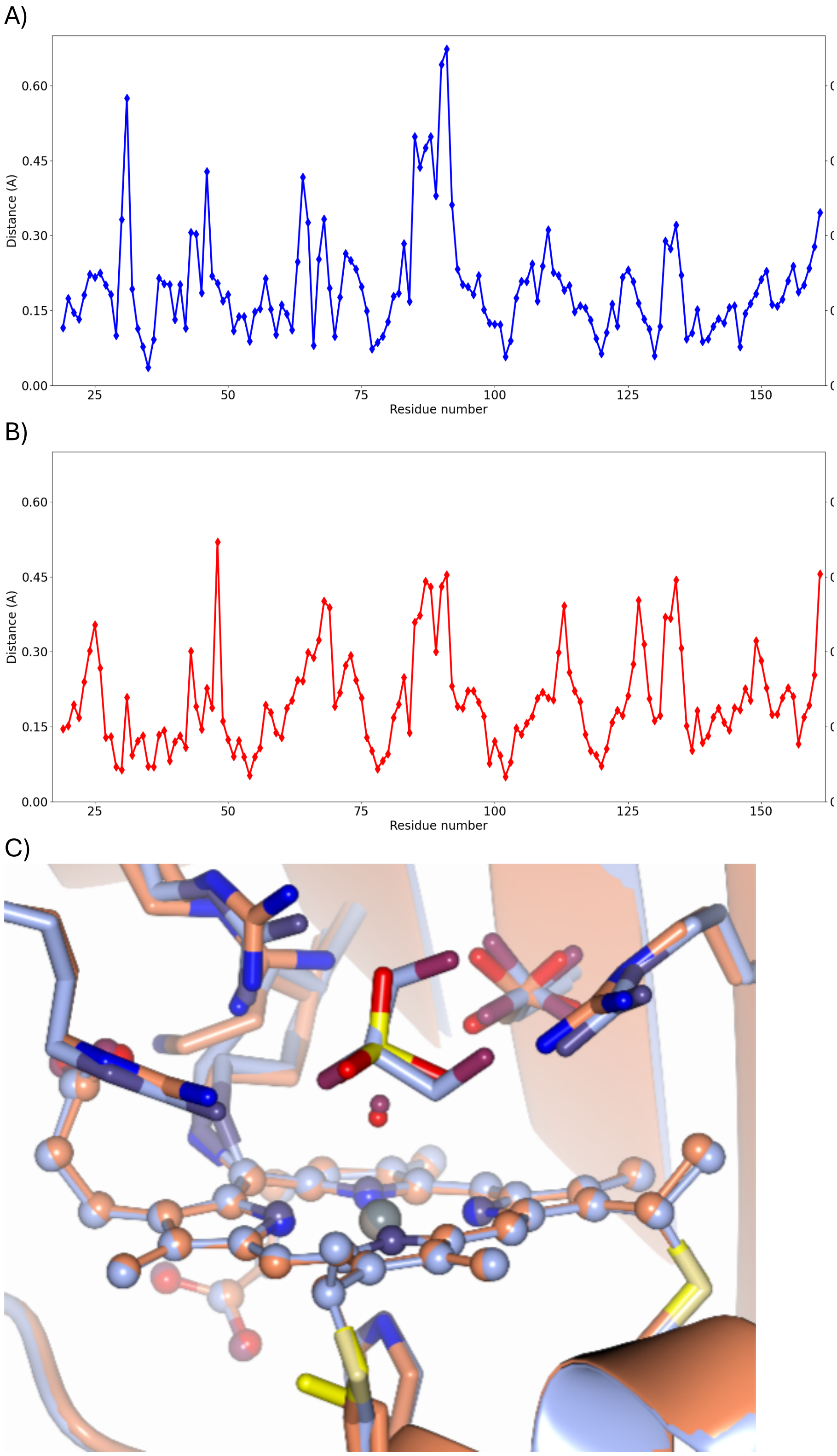


**Fig. S9. A)** RMSD between the Cα atoms of the previously published 100 K structure 6HIU and those of the **SFX** structure for chain A and **B)** for chain B based on their alignment using Gesamt. **C)** Comparison of the active sites in the previously published 100 K structure (PDB 6HIU, blue and cooler colours) and in the damage-free r.t SFX structure (orange and warmer colours). The sulfate (yellow and red) in the SFX structure is instead the site of a cryoprotectant molecule in the 100 K structure.

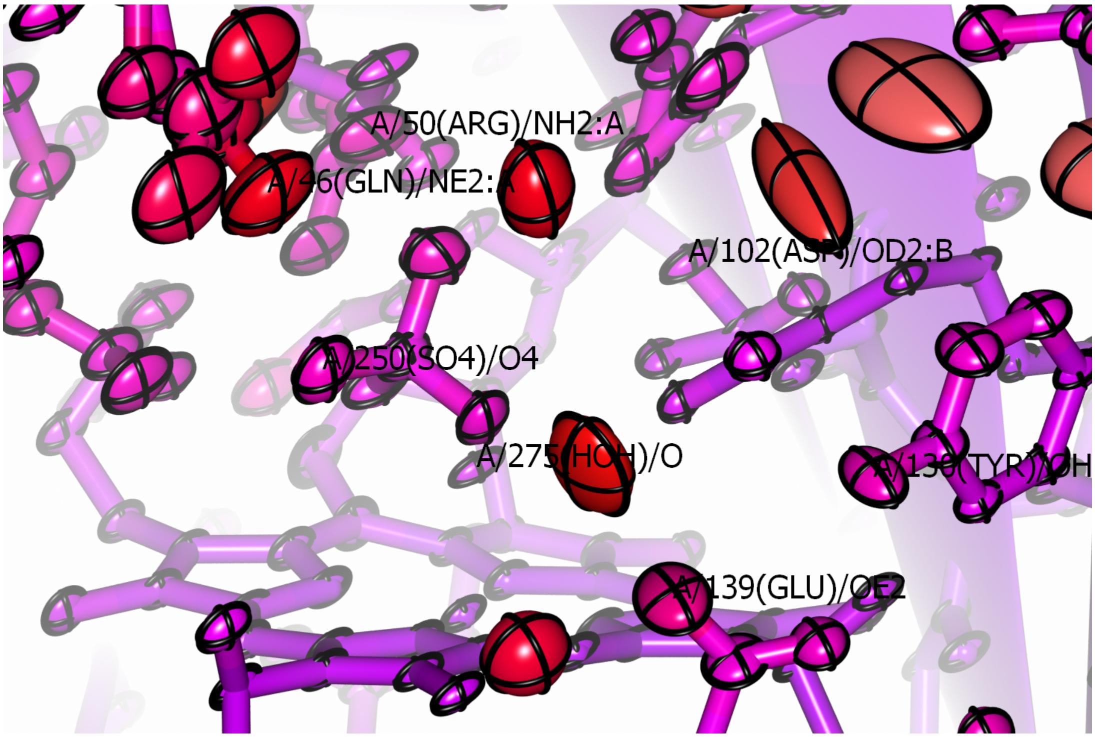


**Fig. S10:** Thermal ellipsoids for the active site of McP460 in the **SFX** structure with bonds represented as cylinders. Color varies from blue to red based on low to high B-factor. Atom labels refer to the atom at their bottom left. Note that Gln46, Arg50 and Asp102 have two conformations each.

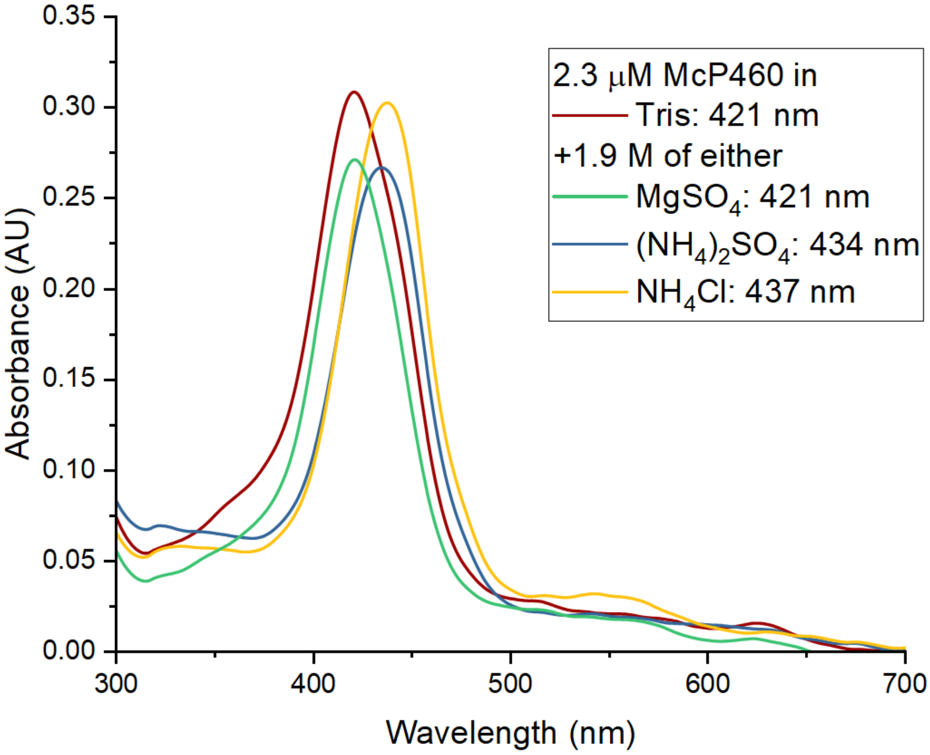


**Fig. S11.** Solution UV-vis spectroscopy of McP460 at pH 8.0, with peak wavelengths in the inset box.

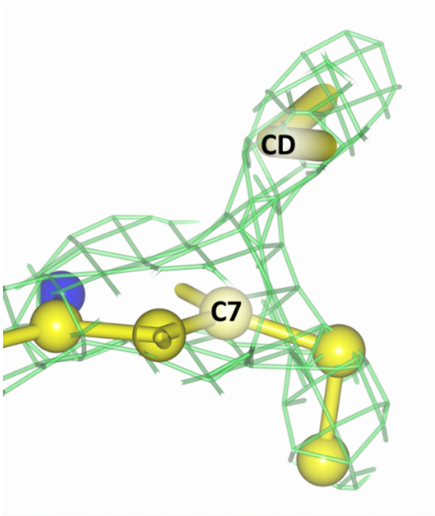


**Fig. S12.** Composite omit electron density map (light green) around the "fresh" crystal structure (yellow) built without new restraints for the double crosslink. It shows planar electron density around the CD carbon of Lys78 that makes a covalent bond to the C7 carbon of the heme.

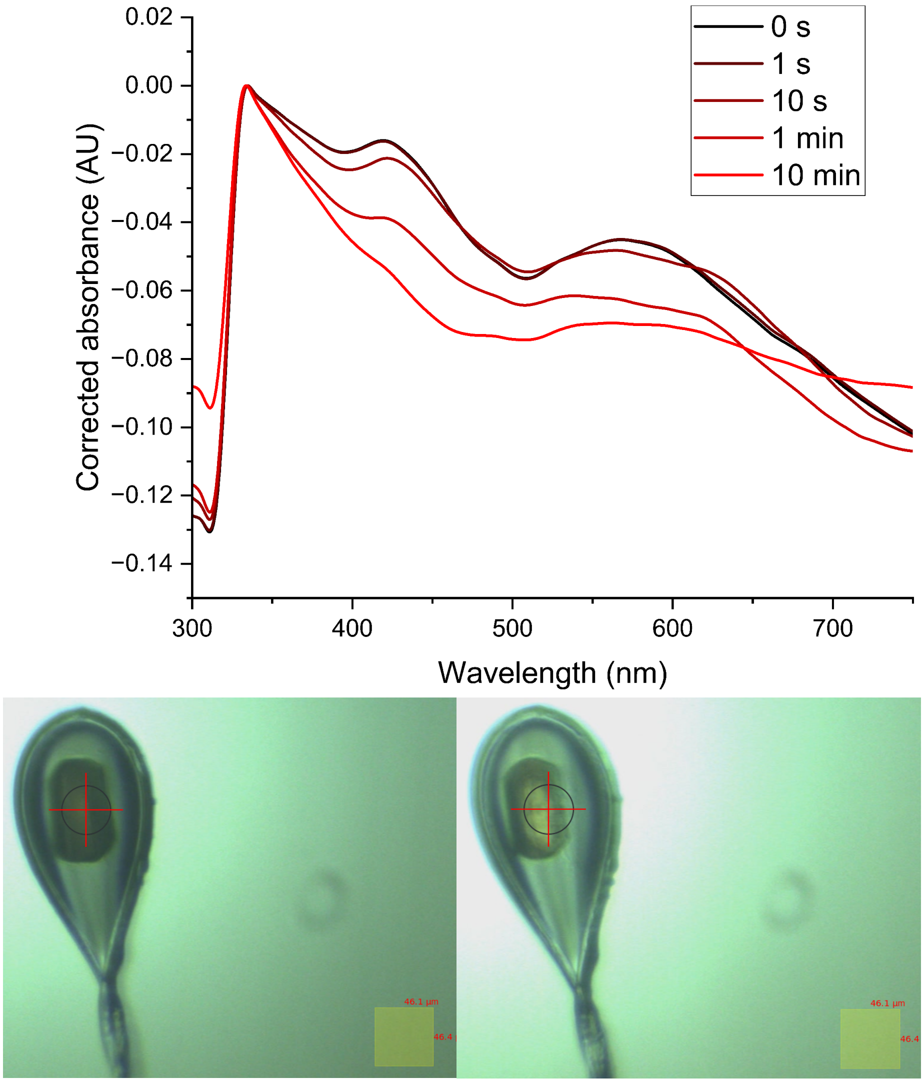


**Fig. S13.** Single-crystal UV-vis spectroscopy (top panel); photographic images of cytochrome P460 crystals illuminated by the white light of the micro spectrophotometer

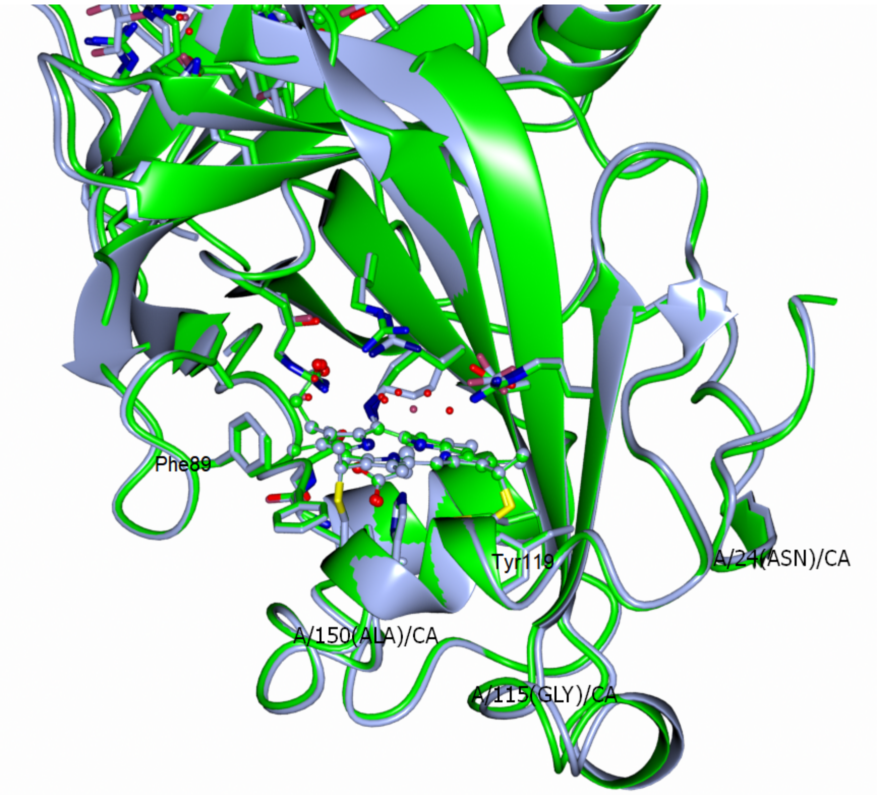


**Fig. S14**. Superposition of reduced 100 K structure and oxidised SFX structure

**Activity Assays**

NH2OH·HCl was equilibrated in an anaerobic environment overnight and then dissolved in 1 mL of deoxygenated water to yield a 1 M solution. This solution was serially diluted 20,000-fold in deoxygenated water and assayed by the method of Frear and Burrell (Frear and Burrell, 1955) for accurate determination of the stock NH2OH concentration. Final concentrations of 50 μM 2,6-dichlorophenolindophenol (DCPIP), 6 μM phenazine methosulfate (PMS), and 1 μM P460 were added to 2 mL of deoxygenated 50 mM sodium phosphate (pH 8.0). The reaction was initiated by adding an appropriate volume of the NH2OH stock solution to the reaction mixture through the septum with a Hamilton syringe. The reaction progress was followed by monitoring the absorption of DCPIP at 605 nm. The rate of the first 10% of the total oxidant consumption was determined through linear regression. This rate was converted to the rate of oxidant consumed by using ε605 nm = 20.6 mM-1 cm-1 (Williamson and Engel, 1984). At least three trials for each NH2OH concentration were averaged and the standard deviations plotted. The initial rates were plotted versus NH2OH concentration, and the resulting plot was fit using linear regression.

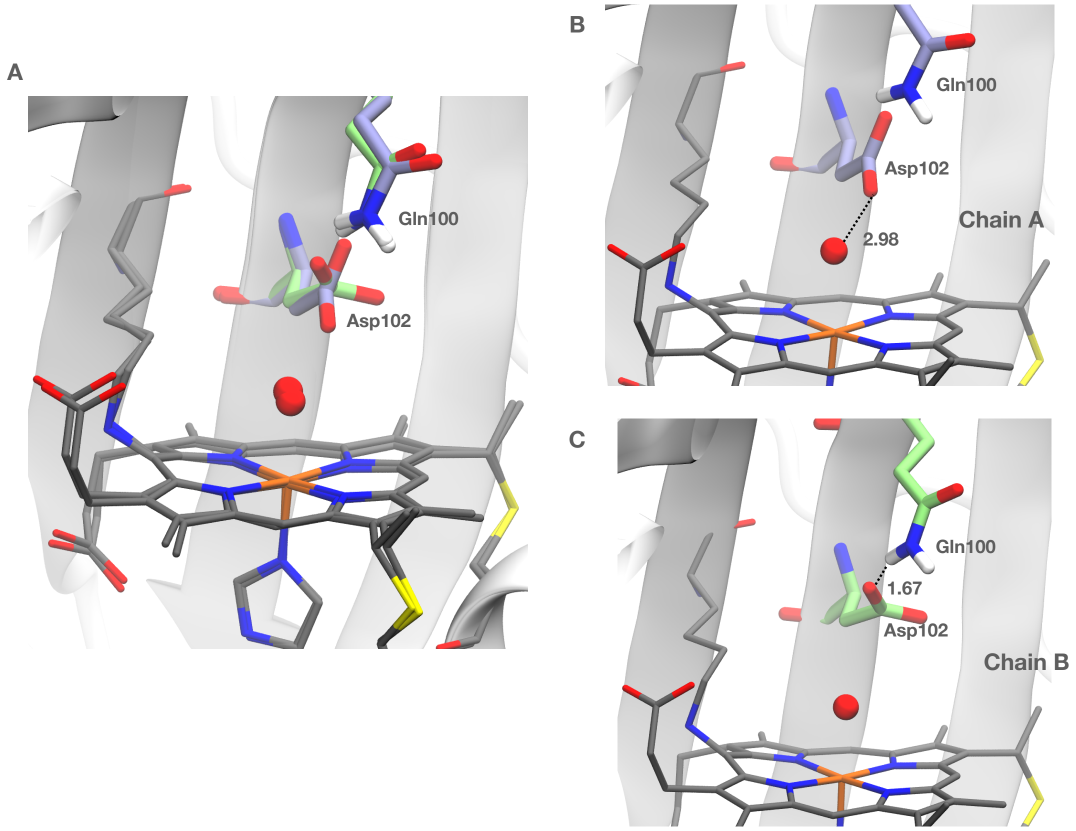


**Fig. S15.** Chain A and B of ferric SFX structure showing A) the two alternating conformations of active site residue Asp102 in chain A (light blue) and chain B (light green). B) Interaction of Asp102 in chain A with the water coordinating to Fe and C) Interaction of Asp102 with another active site residue Gln100. The H-atoms for all residues except the amine group of Gln102 was removed for clarity. Distances are in Å.

**S.3 MD analysis of Ferric-SC/DC systems**

The MD trajectories of Ferric-SC and DC systems were analysed to identify water network patterns in the active site of the protein. The last 150 ns of the 175 ns of the MD was considered for production run and analysis were performed on it. The Ferric **SFX** structure indicated water coordinating to Fe of heme so the MD trajectory was analysed for the presence of a coordinated water by calculating the distance between Fe atom and any water molecule (oxygen atom of the water molecule) within 3.5 Å of the Fe atom. The evolution of the distance of a coordinated water to Fe with time is a given in the top panel of Fig. S16 A(I) for SC and A(II) for DC. Each frame corresponds to 0.1 ns. The average distance of this water from chain A and chain B for SC are 2.73 ± 0.20 Å and 3.24 ± 0.16 Å, respectively. These distances are very similar for DC: 2.8 ± 0.21 Å and 3.25 ± 0.15 Å for chain A and B, respectively. Next, the residence time of water molecule near Fe-atom was calculated as a fraction of time the water molecule was present within 3.5 Å of Fe. To do so, the frequency a water present was binned at intervals of discrete distance + 0.2 Å, normalised to the total time and represented as histograms in the lower panel of Fig. S16 A(I) and A(II) representing SC and DC, respectively. In a similar manner, frequency of H-bond interaction of the water molecule with the pyrrole N atoms of porphyrin ring was calculated as a fraction of the total time, binned with respect to the pyrrole N and plotted as histograms. Same was done for calculating H-bond interaction with the active site residues; frequence of the presence of H-bond interaction was calculated as a fraction of the total time, binned with respect to the active site residue and plotted as histograms.

Furthermore, to verify that the difference in the residence time of the coordinated water in the two chains was a result of the orientation of the Asp102, the MD simulation for both Ferric-SC and Ferric-DC system were repeated with restraining the backbone of the protein with a force constant of 25 kcal/Å$^2$ and allowing the side chains to relax. The same analyses were performed, and the results reveal restoration of similar distribution of water around the active site in both chains (SI Fig. 17) for both SC and DC systems. In case of the SC, the Fe-water distance is 2.85 ± 0.24 Å and 2.80 ± 0.25 Å for chain A and B, respectively and that for double cross link it is 2.63 ± 0.16 Å and 2.62 ± 0.16 Å for chain A and B, respectively. The water has negligible interaction with pyrrole N atoms. The interaction with Asp102 is similar among the two chains but quite different between SC and DC systems. In SC system, water predominantly interacts with Asp102, whereas in double crosslink system this water interacts with both Asp102 and Arg50.

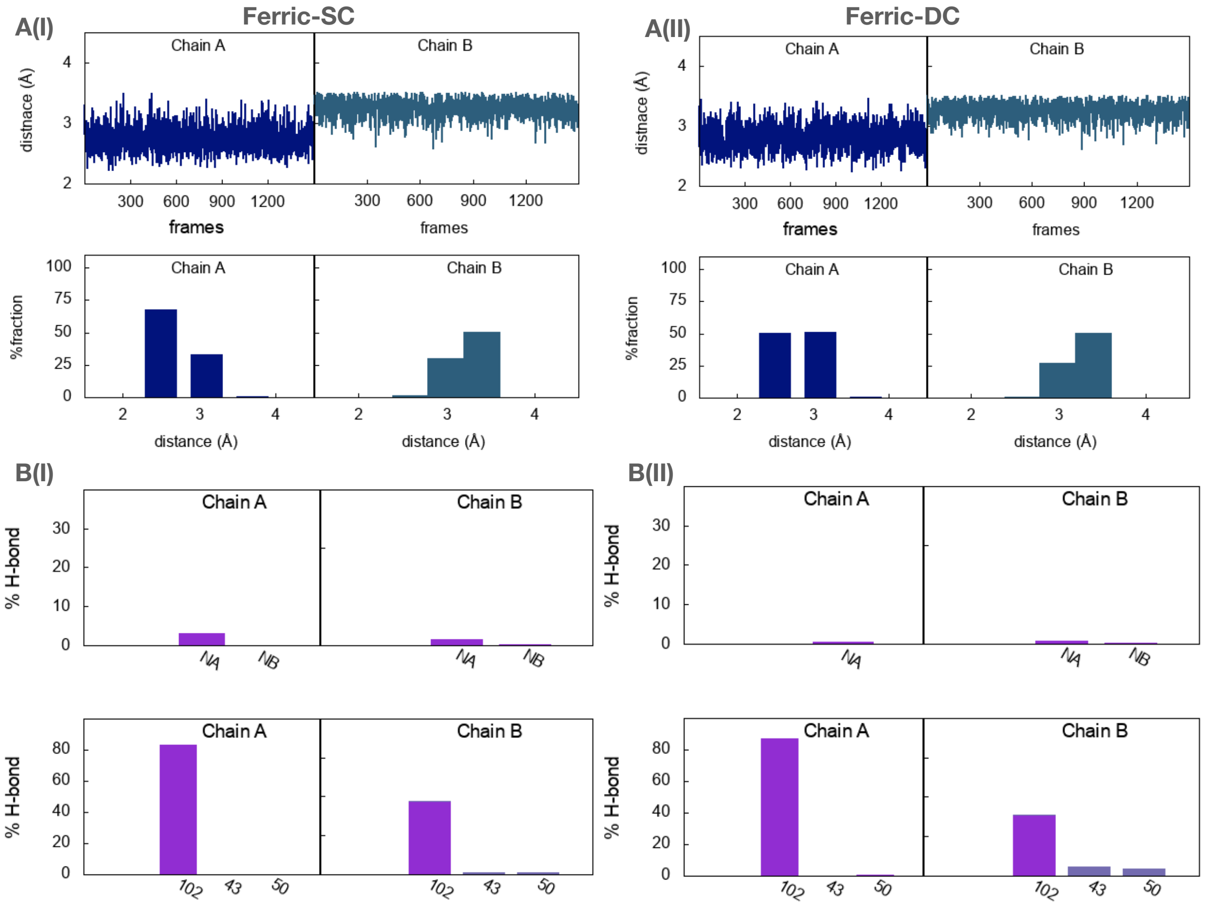


**Fig. S16.** (A) Evolution of the distance of water from Fe along the trajectory (top panel) and residence time of that water at discrete distance intervals (lower panel) is given for (I) for Ferric-SC and (II) for Ferric-DC. (B) Interaction of the above water close to Fe with N-atoms of pyrrole (top panel) and with active site residues Asp102, Arg43 and Arg50 (bottom panel) for (I) Ferric-SC and (II) Ferric-DC. Different water molecules are shown in different colours in the histogram.

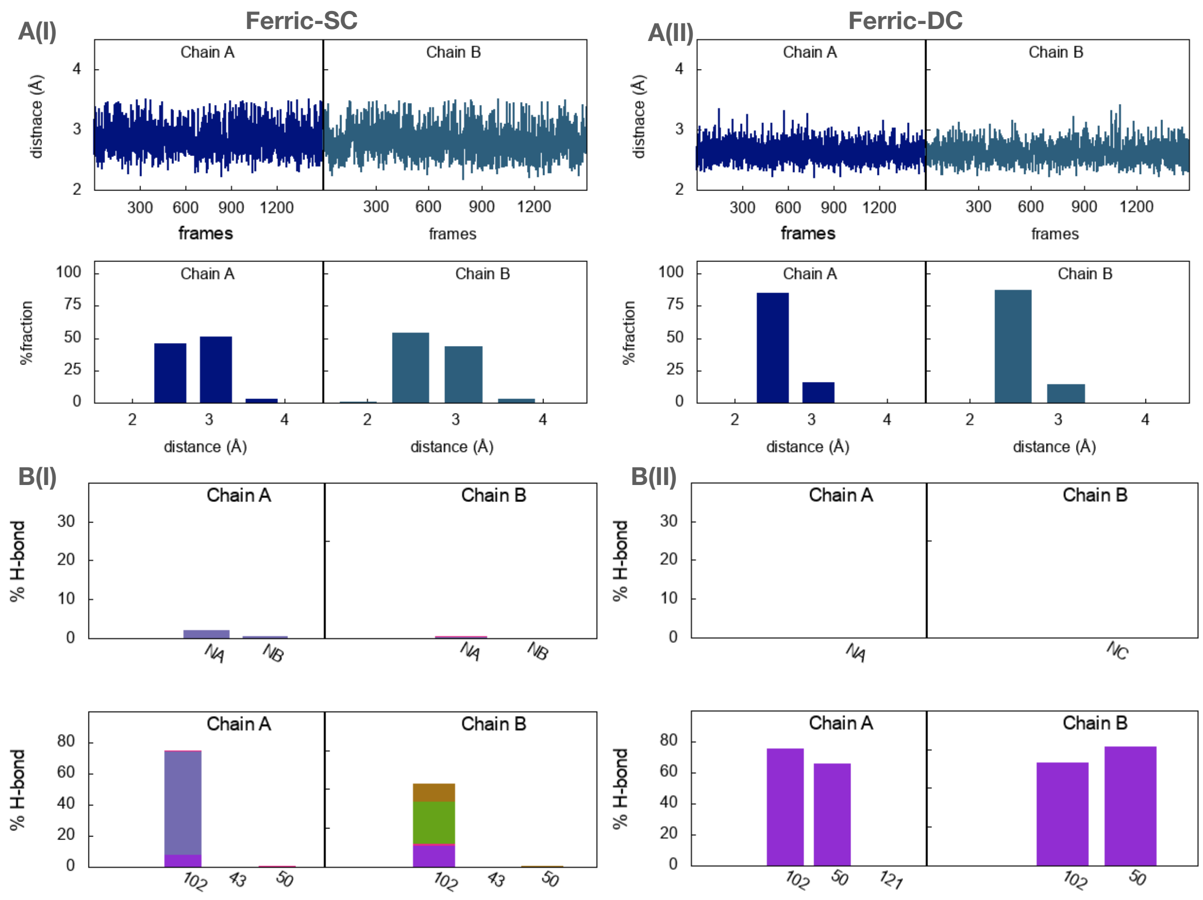


**Fig. S17.** (A) Evolution of the distance of water from Fe along the trajectory (top panel) and residence time of that water at discrete distance intervals (lower panel) is given for (I) for Ferric-SC and (II) for Ferric-DC. (B) Interaction of the above water close to Fe with N-atoms of pyrrole (top panel) and with active site residues Asp102, Arg43 and Arg50 (bottom panel) for (I) Ferric-SC and (II) Ferric-DC. Different water molecules are shown in different colours in the histogram. Here the backbone of the protein was restrained with 25kcal/mol/Å$^2$ force constant and the side chains were allowed to move

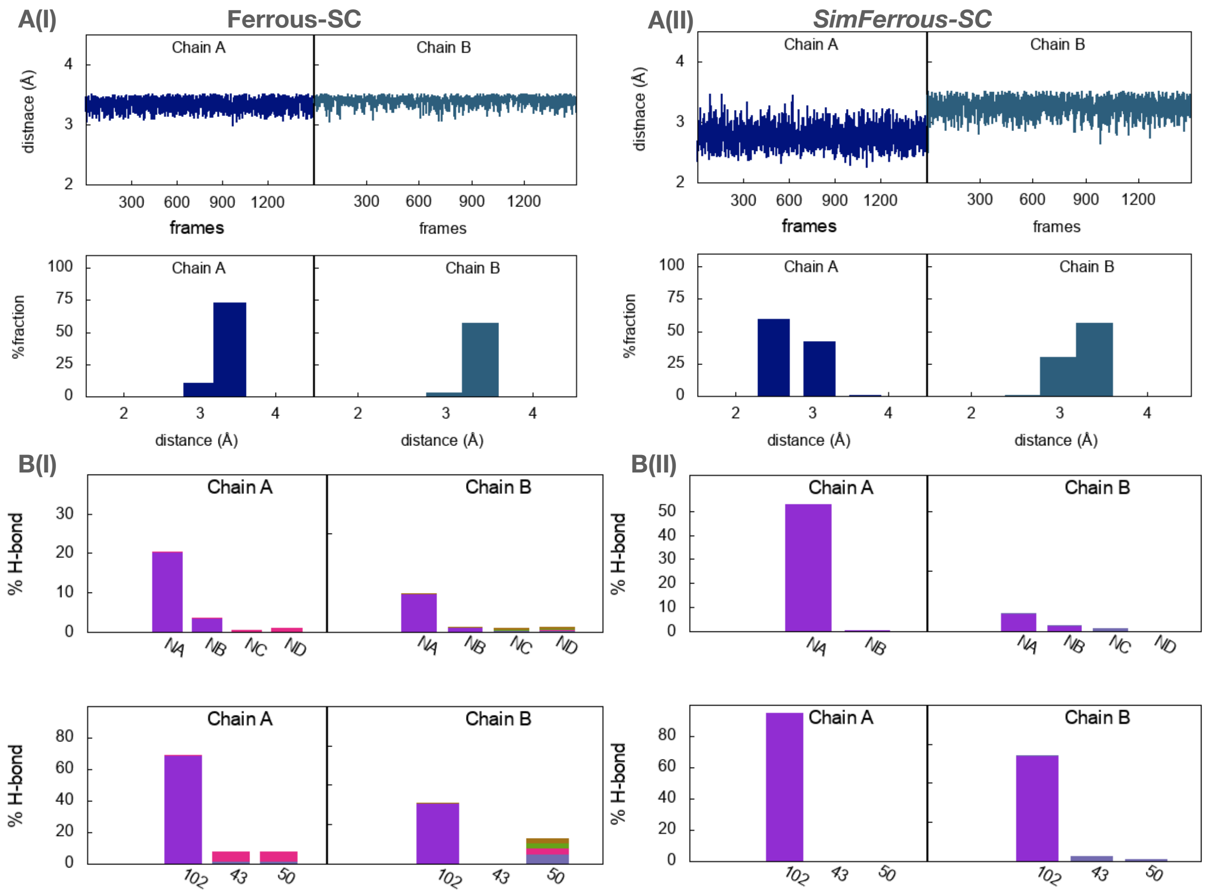


**Fig. S18.** (A) Evolution of the distance of water from Fe along the trajectory (top panel) and residence time of that water at discrete distance intervals (lower panel) is given for (I) Ferrous-SC and (II) *SimFerrous-SC*. (B) Interaction of the above water close to Fe with N-atoms of pyrrole (top panel) and with active site residues Asp102, Arg43 and Arg50 (bottom panel) for (I) Ferrous-SC and (II) *SimFerrous-SC*. Different water molecules are shown in different colours in the histogram.

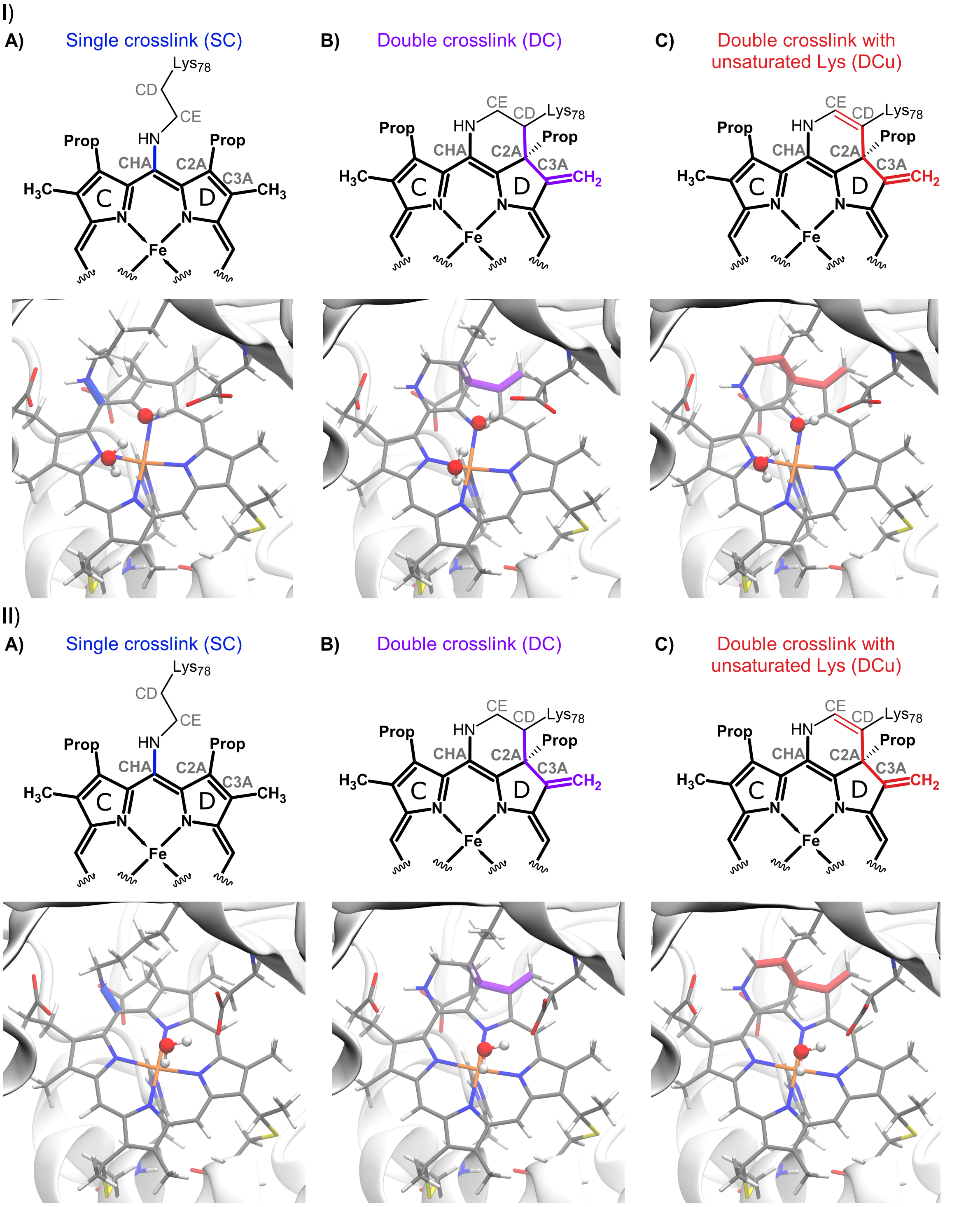


**Fig. 19.** I) QM/MM optimised geometries of A) Ferrous-SC, B) Ferrous-DC and C) Ferrous-DCu. (II) QM/MM optimised geometries of A) *SimFerrous-SC*, B) *SimFerrous-DC* and C) *SimFerrous-DCu*. The heme C unit is shown with sticks and the different crosslinks are shown in the same colour as in Scheme 1 (main manuscript). The 6th site coordinated water is shown in ball and stick. The active site residue Asp102 which interacts with the coordinated water is shown in sticks. Distances are given in Å.

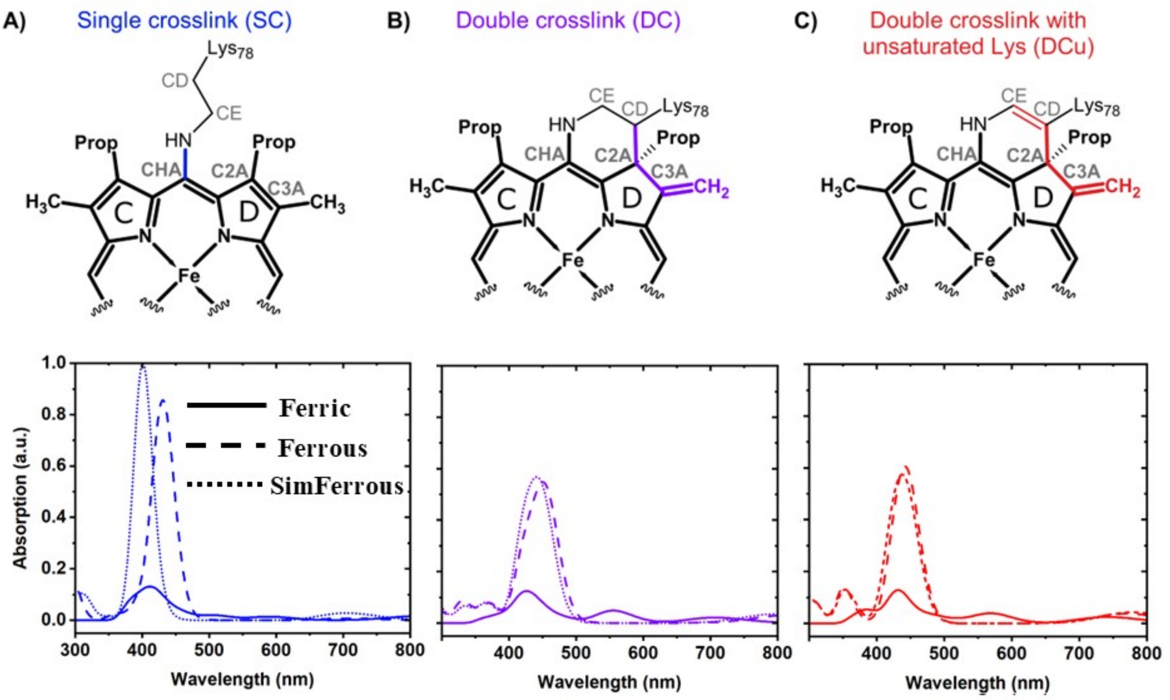


**Fig. S20. Computed absorption spectra for the three crosslink models using ZINDO/S for the lowest spin states.** The Ferrous and *SimFerrous* forms for all models are in a singlet spin state whereas the Ferric forms are in a doublet spin state. ZINDO/S provides good agreement with experiment for the computed Soret band for Ferrous and Ferric forms.

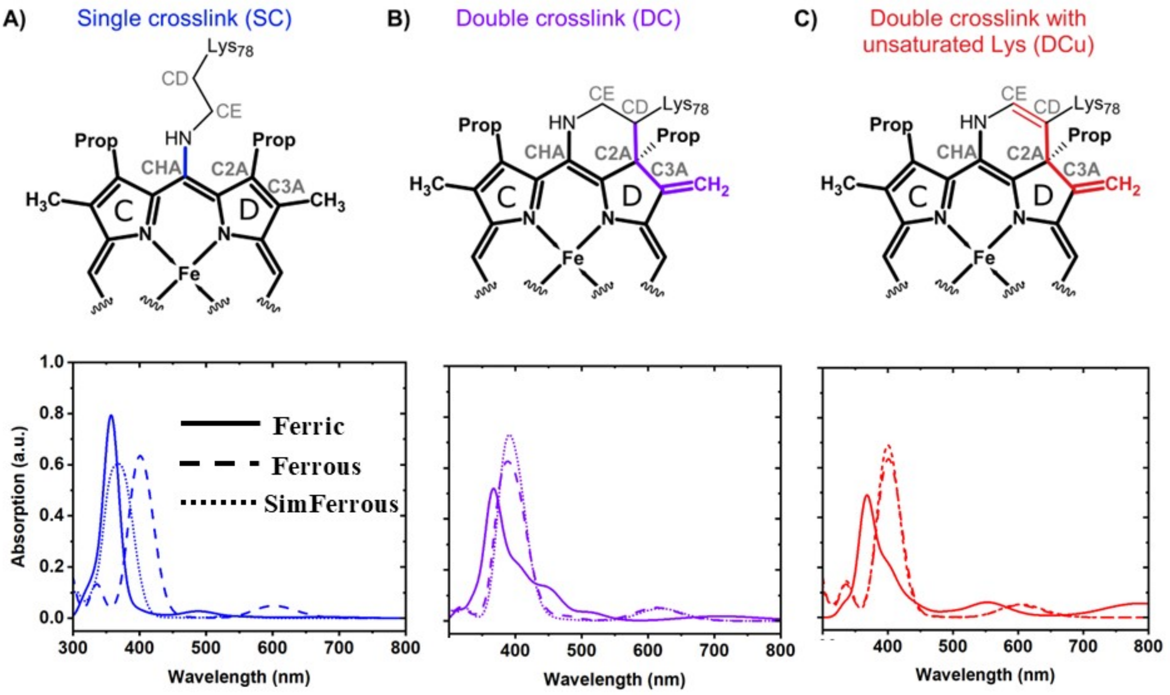


**Fig. S21. Computed absorption spectra for the three crosslink models using CAM-B3LYP (TD-DFT) for the lowest spin states.** The Ferrous and *SimFerrous* forms for all models are in a singlet spin state whereas the Ferric forms are in a doublet spin state.

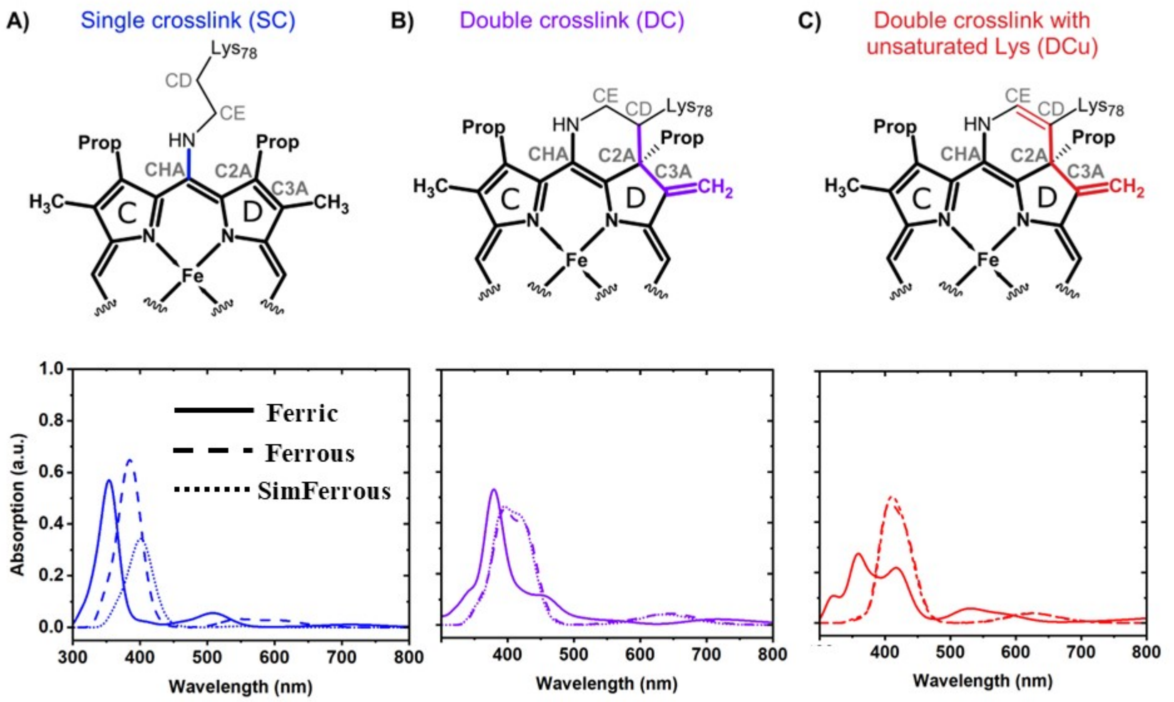


**Fig. S22. Computed absorption spectra for the_three crosslink models using CAM-B3LYP (TD-DFT).** The Ferrous and *SimFerrous* forms for all models are in a triplet spin state whereas the Ferric forms are in a quartet spin state.

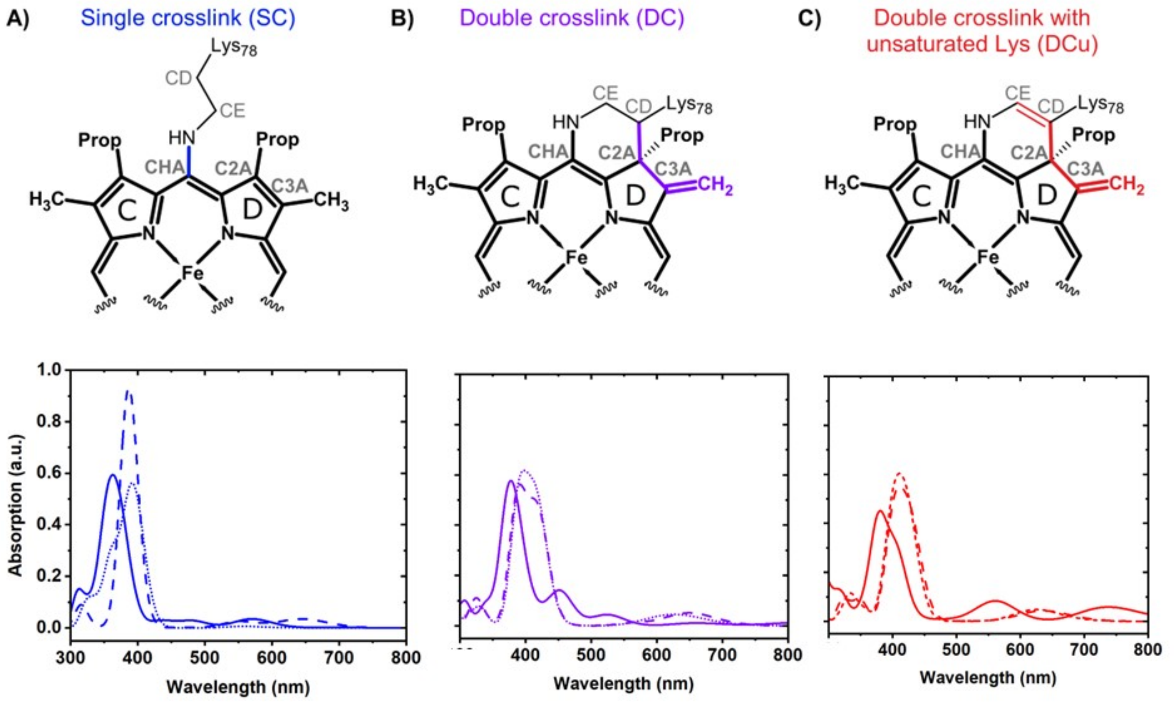


**Fig. S23. Computed absorption spectra for the_three crosslink models using CAM-B3LYP (TD-DFT).** The Ferrous and *SimFerrous* forms for all models are in a quintet spin state whereas the Ferric forms are in a sextet spin state.

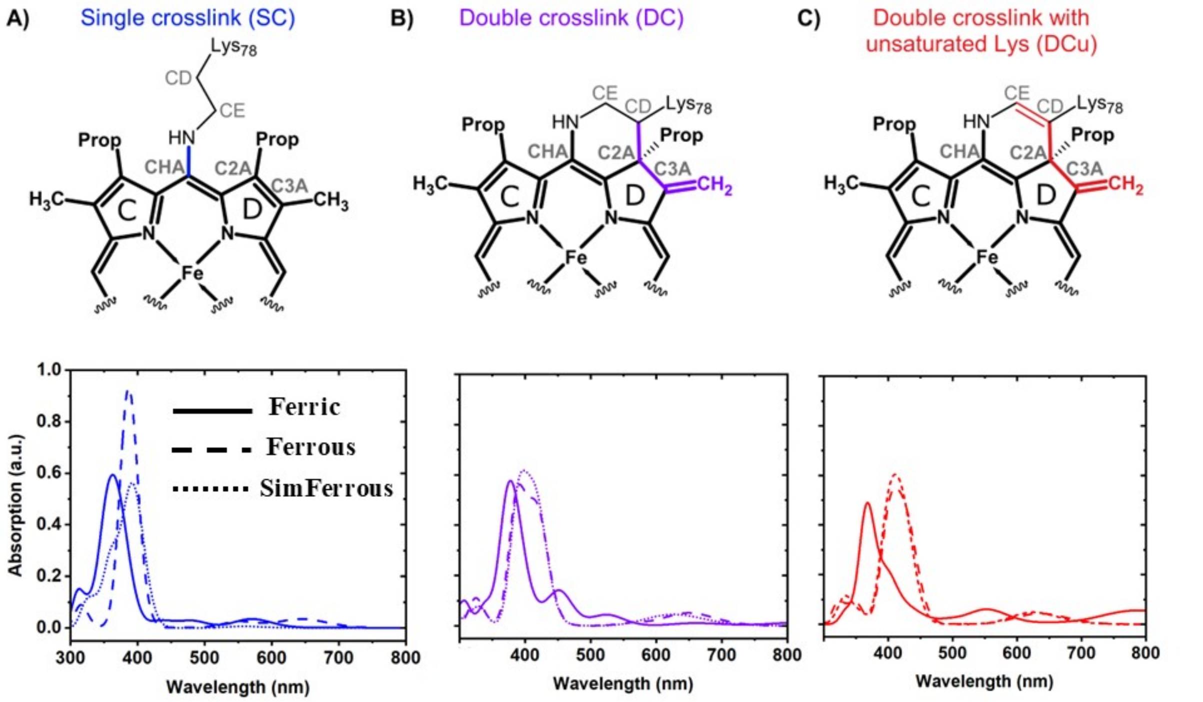


**Fig. S24. Computed absorption spectra for the three crosslink models using CAM-B3LYP (TD-DFT) for the lowest energy spin states.** Ferrous and *SimFerrous* forms for all models are in a quintet spin state whereas the Ferric forms of SC and DC are in a sextet spin state and DCu is in doublet spin state.

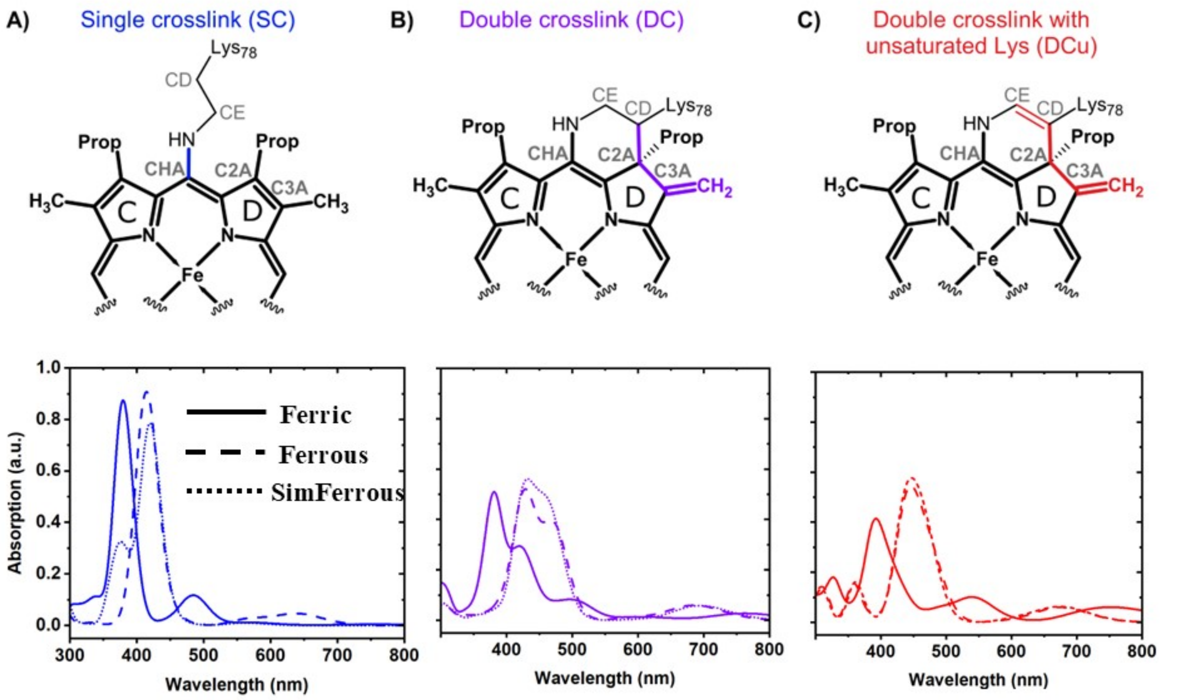


**Fig. S25. Computed absorption spectra for the three crosslink models using CAM-B3LYP (sTD-DFT) for the lowest spin states.** The Ferrous and *SimFerrous* forms for all models are in a singlet spin state whereas the Ferric forms are in a doublet spin state.

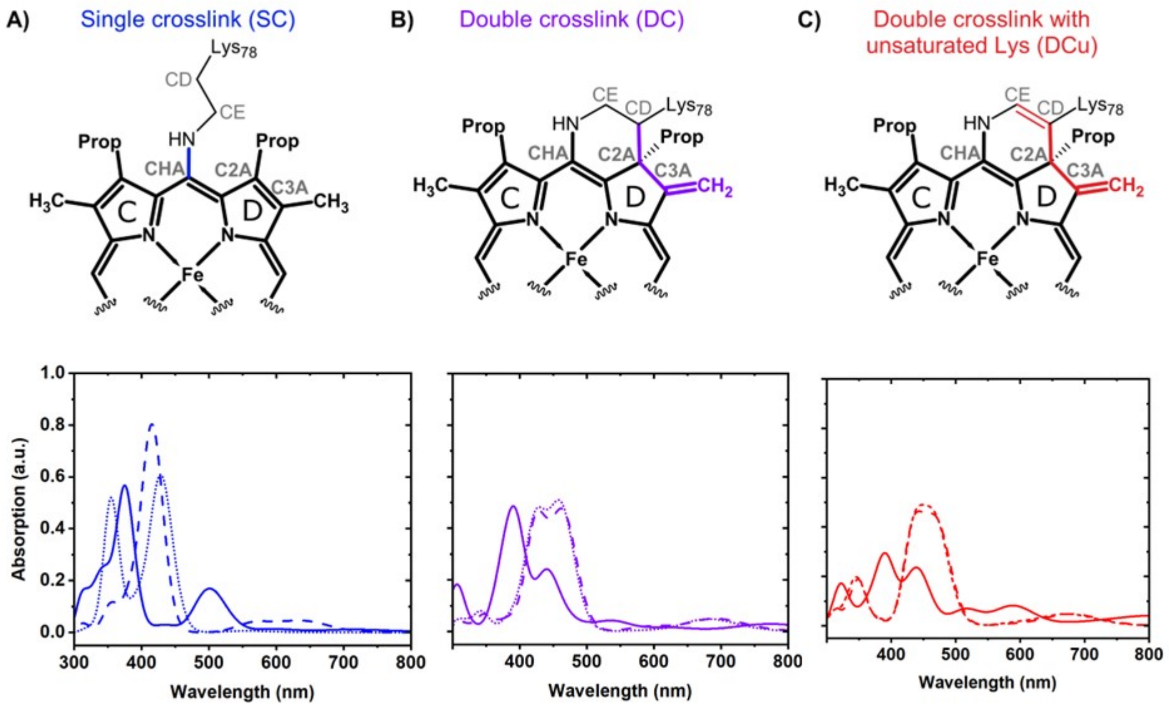


**Fig. S26. Computed absorption spectra for the three crosslink models using CAM-B3LYP (sTD-DFT).** The Ferrous and *SimFerrous* forms for all models are in a triplet spin state whereas the Ferric forms are in a quartet spin state.

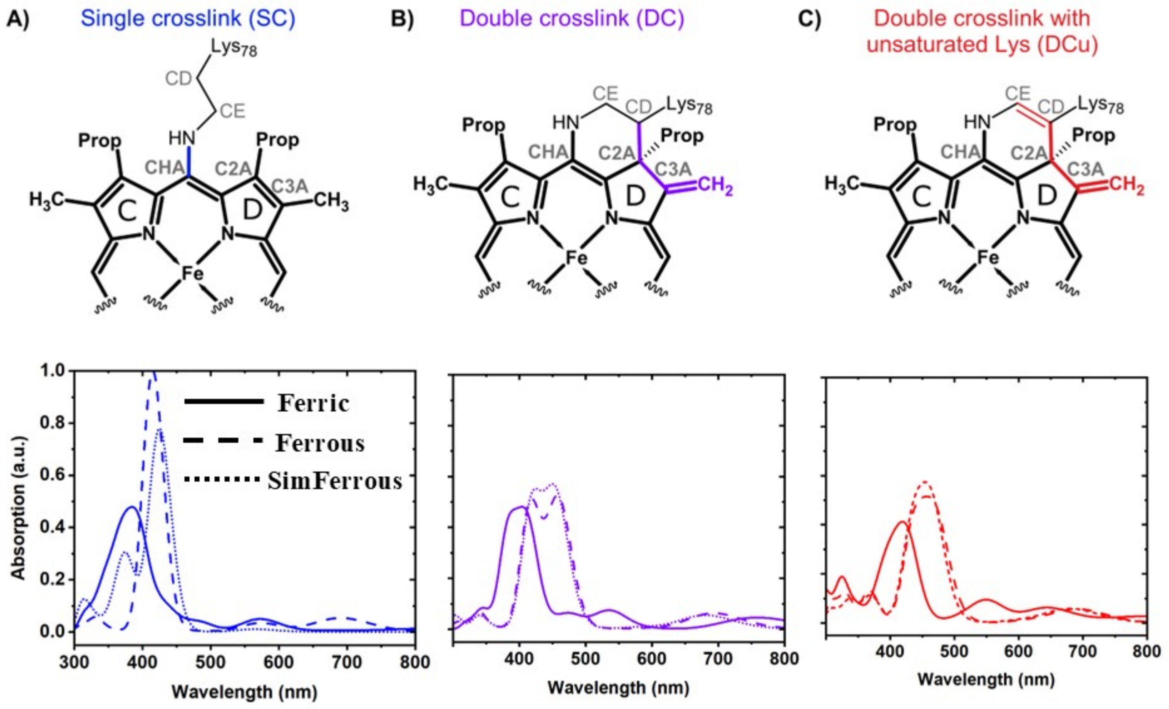


**Fig. S27. Computed absorption spectra for the three crosslink models using CAM-B3LYP (sTD-DFT).** The Ferrous and *SimFerrous* forms for all models are in a quintet spin state whereas the Ferric forms are in a sextet spin state.

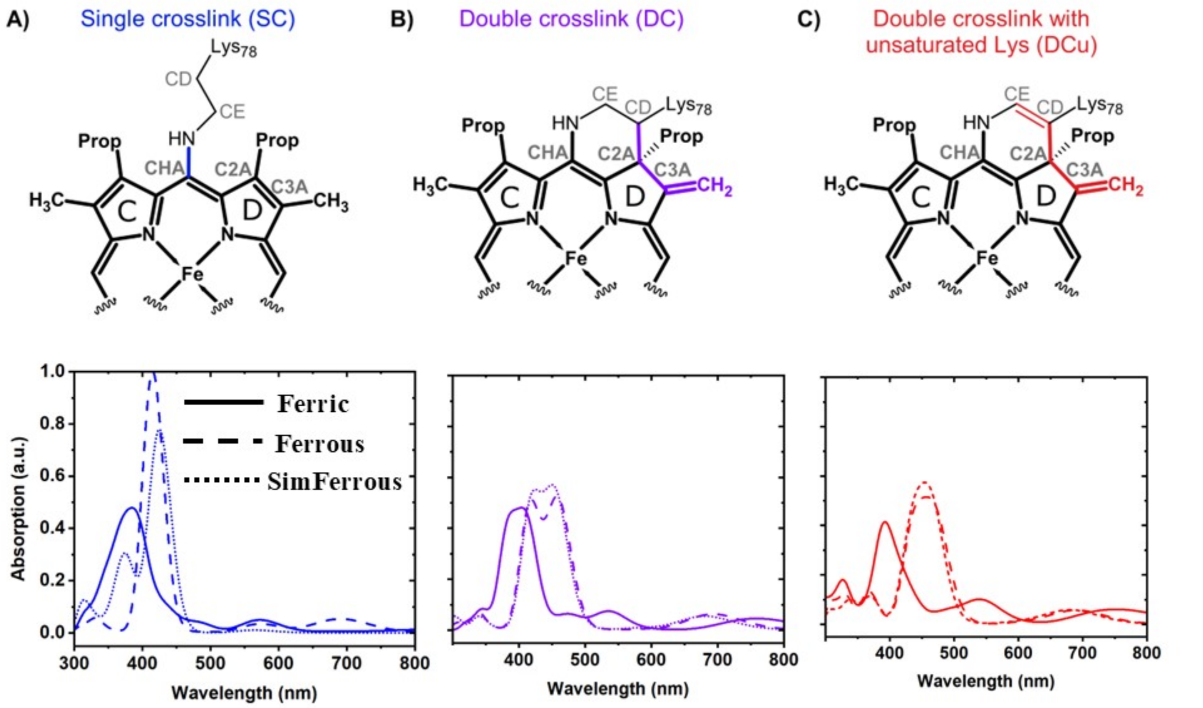


**Fig. S28. Computed absorption spectra for the_three crosslink models using CAM-B3LYP (sTD-DFT) for the lowest energy spin states.** Ferrous and *SimFerrous* forms for all models are in a quintet spin state whereas the Ferric forms of SC and DC are in a sextet spin state and_DCu is in doublet spin state.

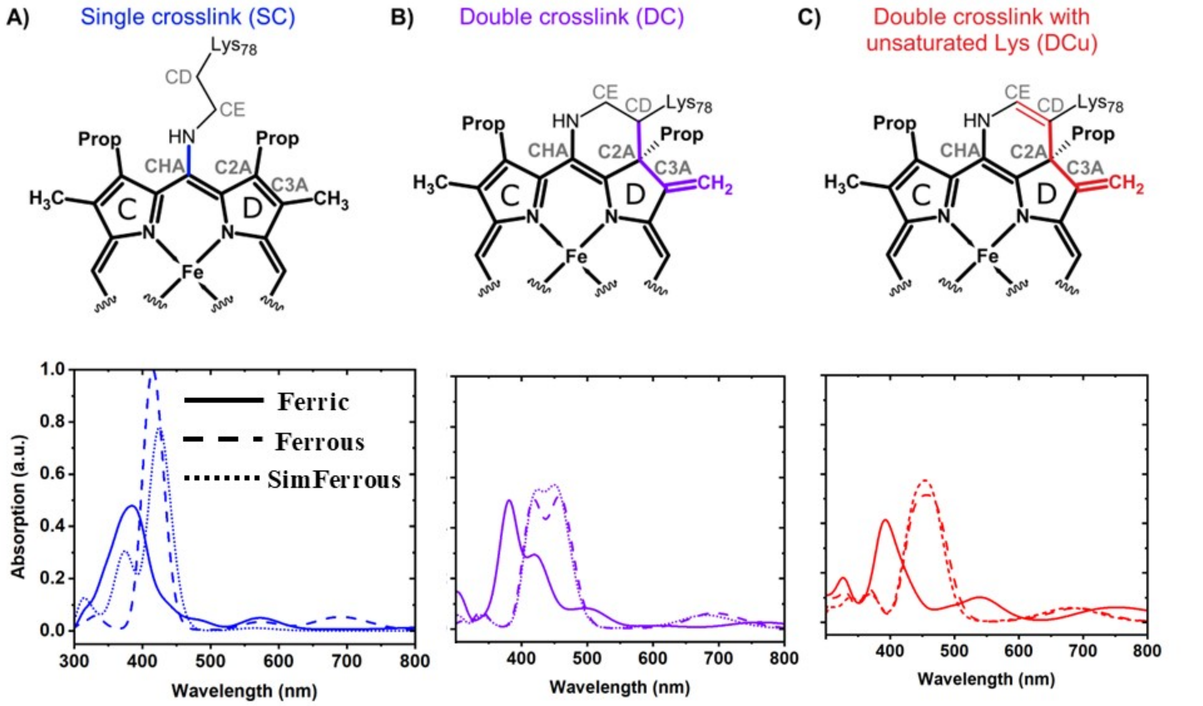


**Fig. S29. Computed absorption spectra for the_three crosslink models using CAM-B3LYP (sTD-DFT).** Ferrous and *SimFerrous* forms for all models are in a quintet spin state whereas the Ferric forms of DC and DCu are in a doublet spin state and SC is in sextet spin state.

**Table S6.** Computed absorption maxima using the ZINDO/S for the lowest spin state for all three models for both Ferrous and Ferric forms. The states correspond to a ππ* transition within the heme moiety. The absorption spectra computed by removing the point charges loses the direction of the shift. However, the DC models still display electronic transitions close to the experiment Soret peak at 460 nm.

| Model | ZINDO/S | | | ZINDO/S without pointcharges | | |
|---|---|---|---|---|---|---|
| | Ferrous<br>nm ($f_{osc}$) | SimFerrous<br>nm ($f_{osc}$) | Ferric<br>Nm ($f_{osc}$) | Ferrous<br>nm ($f_{osc}$) | SimFerrous<br>nm ($f_{osc}$) | Ferric<br>nm ($f_{osc}$) |
| DC | 457.1 (1.33)<br>436.5 (0.43) | 452.3(0.86) | 426.7 (0.19) | 445.9 (1.15)<br>430.5 (0.16) | 447.4 (0.25)<br>440.5 (1.05) | 458.8 (0.15)<br>421.9 (0.11) |
| DCu | 450.3 (1.08)<br>430.5 (0.92) | 449.3 (0.79) | 429.9 (0.31) | 438.4 (1.19) | 443.9 (0.18)<br>437.4 (1.06) | 464.4 (0.12) |
| SC | 434.9 (1.20)<br>428.0 (1.41) | 418.8 (0.15) | 402.5 (1.46) | 436.8 (1.11)<br>424.7 (1.34) | 401.8 (1.46)<br>399.0 (1.46) | 416.6 (0.30) |

**Table S7.** Computed absorption maxima using TD-DFT for all spin states for the three proposed models.

| Model | Ferrous nm ($f_{osc}$) (M=1) | SimFerrous nm ($f_{osc}$) (M=1) | Ferrous nm ($f_{osc}$) (M=3) | SimFerrous nm ($f_{osc}$) (M=3) | Ferrous nm ($f_{osc}$) (M=5) | SimFerrous nm ($f_{osc}$) (M=5) | Ferric nm ($f_{osc}$) (M=2) | Ferric nm ($f_{osc}$) (M=4) | Ferric nm ($f_{osc}$) (M=6) |
|---|---|---|---|---|---|---|---|---|---|
| DC | 405.3 (0.46) 379.0 (0.45) | 406.0 (0.47) 387.6 (0.54) | 429.3 (0.42) | 403.1 (0.52) | 419.3 (0.61) 389.4 (0.46) | 418.0 (0.65) 391.2 (0.60) | 410.4 (0.14) 396.2 (0.14) | 455.6 (0.1) 419.0 (0.1) | 453.7 (0.17) 388.1 (0.33) |
| DCu | 415.3 (0.32) 394.2 (0.56) | 409.9 (0.48) 393.8 (0.65) | 436.1 (0.22) 400.9 (0.45) | 434.6 (0.25) | 430.9 (0.23) 402.0 (0.57) | 425.5 (0.52) 399.9 (0.57) | 397.3 (0.16) | 424.2 (0.16) | 567.7 (0.1) 408.7 (0.2) |
| SC | 377.6 (0.8) 371.4 (0.57) | 389.7 (0.24) | 390.3 (0.64) | 410.2 (0.15) 397.5 (0.21) | 389.4 (0.59) | 401.1 (0.38) 387.4 (0.52) 360.7 (0.37) | 358.7 (0.62) 357.6 (0.46) | 361.8 (0.25) | 386.0 (0.13) 374.3 (0.22) |

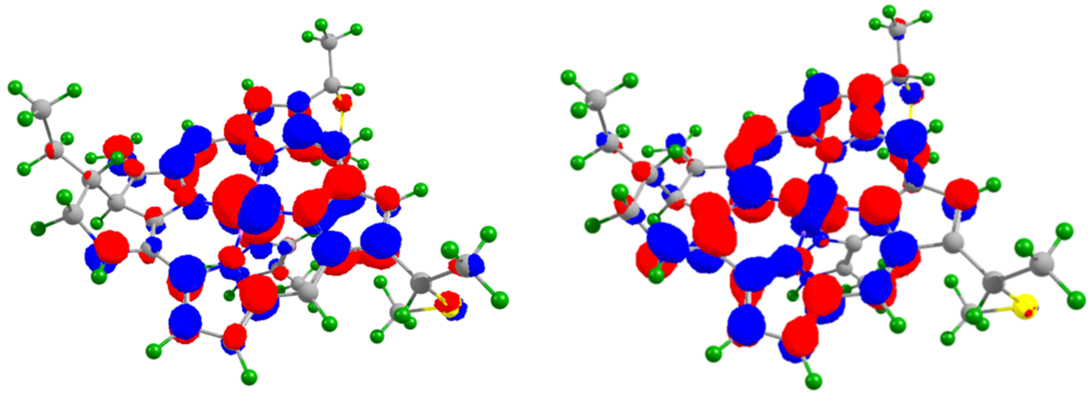

**Fig. S30.** The hole (left) and particle (right) densities for the NTOs corresponding to the Soret transition for the Ferrous-DC model.

Heme restraints

### MEH restraints

Used for SFX and ferrous cryo structure. Require the heme monomer of the PDB file to be renamed from HEC to MEH to apply. The effects of those restraints include allowing more distortion in the plane of the heme.

```
global_
_lib_name         mon_lib
_lib_version      5.39
_lib_update       06/11/2012
# ------------------------------------------------
#
# ---   LIST OF MONOMERS ---
#
data_comp_list
loop_
_chem_comp.id
_chem_comp.three_letter_code
_chem_comp.name
_chem_comp.group
_chem_comp.number_atoms_all
_chem_comp.number_atoms_nh
_chem_comp.desc_level
MEH       MEH 'HEM SINGLE CAB-CBB CAC-CBC          ' .                    77
43 .
No        No  '.                                    ' .                    3
2 .
#
# ---   LIST OF LINKS ---
#
data_link_list
loop_
_chem_link.id
_chem_link.comp_id_1
_chem_link.mod_id_1
_chem_link.group_comp_1
_chem_link.comp_id_2
_chem_link.mod_id_2
_chem_link.group_comp_2
_chem_link.name
CYS-MEHB CYS      CYS-HG   .        MEH      MEH-HABA .
CYS-MEHB
CYS-MEHC CYS      CYS-HG   .        MEH      MEH-HACA .
CYS-MEHC
HIS-MEH  HIS      HIS-HE2  .        MEH      .        .
HIS-MEH
#
# ---   LIST OF MODIFICATIONS ---
#
data_mod_list
loop_
_chem_mod.id
_chem_mod.name
_chem_mod.comp_id
_chem_mod.group_id
CYS-HG   CYSTINE                                          CYS      .
MEH-HABA MEH_DELETED_HABA                                 MEH      .
MEH-HACA MEH_DELETED_HACA                                 MEH      .
HIS-HE2  HISTIDINE-DELETED-HE2                            HIS      .
#
```

```
# --- DESCRIPTION OF MONOMERS ---
#
data_comp_MEH
#
loop_
_chem_comp_atom.comp_id
_chem_comp_atom.atom_id
_chem_comp_atom.type_symbol
_chem_comp_atom.type_energy
_chem_comp_atom.partial_charge
_chem_comp_atom.x
_chem_comp_atom.y
_chem_comp_atom.z
MEH           O2D    O    OC        -0.500       5.649   -3.481   -2.121
MEH           CGD    C    C          0.000       6.133   -3.500   -0.968
MEH           O1D    O    OC        -0.500       6.686   -2.506   -0.450
MEH           CBD    C    CH2        0.000       6.052   -4.778   -0.159
MEH           HBD    H    H          0.000       6.418   -5.607   -0.770
MEH           HBDA   H    H          0.000       6.728   -4.688    0.694
MEH           CAD    C    CH2        0.000       4.649   -5.112    0.368
MEH           HAD    H    H          0.000       4.578   -5.305   -0.704
MEH           HADA   H    H          0.000       4.615   -4.117    0.815
MEH           C3D    C    CR5        0.000       3.174   -4.966    0.026
MEH           C2D    C    CR5        0.000       2.275   -6.011    0.023
MEH           CMD    C    CH3        0.000       2.547   -7.456    0.366
MEH           HMDB   H    H          0.000       3.579   -7.598    0.549
MEH           HMDA   H    H          0.000       2.246   -8.070   -0.444
MEH           HMD    H    H          0.000       1.996   -7.721    1.230
MEH           C4D    C    CR5        0.000       2.419   -3.874   -0.416
MEH           CHA    C    C1         0.000       2.955   -2.503   -0.476
MEH           HHA    H    H          0.000       4.004   -2.385   -0.320
MEH           ND     N    NR5        0.000       1.098   -4.116   -0.520
MEH           C1D    C    CR5        0.000       1.074   -5.453   -0.377
MEH           CHD    C    C1         0.000      -0.154   -6.263   -0.428
MEH           HHD    H    H          0.000      -0.055   -7.314   -0.261
MEH           FE     FE   FE         0.000      -0.441   -2.869   -1.227
MEH           NB     N    NR5        0.000      -1.982   -1.592   -0.509
MEH           C4B    C    CR5        0.000      -3.304   -1.829   -0.323
MEH           C3B    C    CR5        0.000      -4.066   -0.739    0.087
MEH           CAB    C    CH2        0.000      -5.532   -0.710    0.473
MEH           HAB    H    H          0.000      -5.782    0.242    0.944
MEH           CBB    C    CH3        0.000      -6.416   -0.921   -0.740
MEH           HBBA   H    H          0.000      -6.409   -0.043   -1.333
MEH           HBB    H    H          0.000      -7.407   -1.118   -0.422
MEH           C2B    C    CR5        0.000      -3.184    0.311    0.075
MEH           CMB    C    CH3        0.000      -3.451    1.762    0.398
MEH           HMBB   H    H          0.000      -4.481    1.912    0.590
MEH           HMBA   H    H          0.000      -3.160    2.362   -0.425
MEH           HMB    H    H          0.000      -2.890    2.041    1.252
MEH           C1B    C    CR5        0.000      -1.991   -0.245   -0.361
MEH           CHB    C    C1         0.000      -0.796    0.599   -0.442
MEH           HHB    H    H          0.000      -0.907    1.650   -0.279
MEH           NC     N    NR5        0.000      -1.767   -4.384   -0.565
MEH           C4C    C    CR5        0.000      -1.504   -5.690   -0.348
MEH           C3C    C    CR5        0.000      -2.580   -6.464    0.081
MEH           CAC    C    CH2        0.000      -2.590   -7.937    0.441
MEH           HAC    H    H          0.000      -1.762   -8.463   -0.035
MEH           CBC    C    CH3        0.000      -2.522   -8.123    1.944
MEH           HBCA   H    H          0.000      -2.666   -9.147    2.178
MEH           HBC    H    H          0.000      -1.571   -7.814    2.293
MEH           C2C    C    CR5        0.000      -3.637   -5.577    0.123
```

```
MEH           CMC    C    CH3        0.000      -5.089   -5.828    0.476
MEH           HMCB   H    H          0.000      -4.695   -6.606    1.067
MEH           HMCA   H    H          0.000      -5.184   -5.721   -0.574
MEH           HMC    H    H          0.000      -5.754   -6.616    0.233
MEH           C1C    C    CR5        0.000      -3.088   -4.380   -0.320
MEH           CHC    C    C1         0.000      -3.897   -3.168   -0.355
MEH           HHC    H    H          0.000      -4.947   -3.270   -0.163
MEH           NA     N    NR5        0.000       0.799   -1.266   -0.613
MEH           C1A    C    CR5        0.000       2.135   -1.281   -0.410
MEH           C4A    C    CR5        0.000       0.547    0.037   -0.398
MEH           C3A    C    CR5        0.000       1.637    0.810   -0.016
MEH           CMA    C    CH3        0.000       1.673    2.278    0.340
MEH           HMAB   H    H          0.000       0.916    2.800   -0.185
MEH           HMAA   H    H          0.000       1.509    2.386    1.380
MEH           HMA    H    H          0.000       2.616    2.689    0.092
MEH           C2A    C    CR5        0.000       2.692   -0.070   -0.006
MEH           CAA    C    CH2        0.000       4.135    0.260    0.316
MEH           HAA    H    H          0.000       4.164    0.876    1.218
MEH           HAAA   H    H          0.000       4.709   -0.645    0.523
MEH           CBA    C    CH2        0.000       4.782    1.030   -0.830
MEH           HBA    H    H          0.000       4.144    1.857   -1.144
MEH           HBAA   H    H          0.000       4.912    0.365   -1.687
MEH           CGA    C    C          0.000       6.129    1.559   -0.396
MEH           O1A    O    OC        -0.500       6.332    2.791   -0.456
MEH           O2A    O    OC        -0.500       6.989    0.747    0.009
MEH           HABA   H    H          0.000      -5.727   -1.498    1.202
MEH           HBBB   H    H          0.000      -6.062   -1.733   -1.318
MEH           HACA   H    H          0.000      -3.514   -8.386    0.071
MEH           HBCB   H    H          0.000      -3.271   -7.546    2.418
loop_
_chem_comp_tree.comp_id
_chem_comp_tree.atom_id
_chem_comp_tree.atom_back
_chem_comp_tree.atom_forward
_chem_comp_tree.connect_type
MEH      O2D    CGD    .      .
MEH      CGD    CBD    O2D    .
MEH      O1D    CGD    .      .
MEH      CBD    CAD    CGD    .
MEH      HBD    CBD    .      .
MEH      HBDA   CBD    .      .
MEH      CAD    C3D    CBD    .
MEH      HAD    CAD    .      .
MEH      HADA   CAD    .      .
MEH      C3D    C4D    CAD    .
MEH      C2D    C1D    CMD    .
MEH      CMD    C2D    HMDB   .
MEH      HMDB   CMD    .      END
MEH      HMDA   CMD    .      .
MEH      HMD    CMD    .      .
MEH      C4D    ND     C3D    .
MEH      CHA    C1A    HHA    .
MEH      HHA    CHA    .      .
MEH      ND     FE     C4D    .
MEH      C1D    CHD    C2D    .
MEH      CHD    C4C    C1D    .
MEH      HHD    CHD    .      .
MEH      FE     NA     NC     .
MEH      NB     C1B    .      .
MEH      C4B    CHC    .      .
MEH      C3B    C2B    CAB    .
```

```
MEH      CAB    C3B    CBB     .
MEH      HAB    CAB    .       .
MEH      CBB    CAB    HBBA    .
MEH      HBBA   CBB    .       .
MEH      HBB    CBB    .       .
MEH      C2B    C1B    C3B     .
MEH      CMB    C2B    HMBB    .
MEH      HMBB   CMB    .       .
MEH      HMBA   CMB    .       .
MEH      HMB    CMB    .       .
MEH      C1B    CHB    NB      .
MEH      CHB    C4A    C1B     .
MEH      HHB    CHB    .       .
MEH      NC     FE     C1C     .
MEH      C4C    C3C    CHD     .
MEH      C3C    C2C    C4C     .
MEH      CAC    C3C    CBC     .
MEH      HAC    CAC    .       .
MEH      CBC    CAC    HBCA    .
MEH      HBCA   CBC    .       .
MEH      HBC    CBC    .       .
MEH      C2C    C1C    C3C     .
MEH      CMC    C2C    HMCB    .
MEH      HMCB   CMC    .       .
MEH      HMCA   CMC    .       .
MEH      HMC    CMC    .       .
MEH      C1C    NC     C2C     .
MEH      CHC    C1C    C4B     .
MEH      HHC    CHC    .       .
MEH      NA     C4A    FE      .
MEH      C1A    C2A    CHA     .
MEH      C4A    C3A    NA      .
MEH      C3A    C2A    C4A     .
MEH      CMA    C3A    HMAB    .
MEH      HMAB   CMA    .       .
MEH      HMAA   CMA    .       .
MEH      HMA    CMA    .       .
MEH      C2A    CAA    C3A     .
MEH      CAA    CBA    C2A     .
MEH      HAA    CAA    .       .
MEH      HAAA   CAA    .       .
MEH      CBA    CGA    CAA     .
MEH      HBA    CBA    .       .
MEH      HBAA   CBA    .       .
MEH      CGA    O2A    CBA     .
MEH      O1A    CGA    .       .
MEH      O2A    n/a    CGA     START
MEH      HABA   CAB    .       .
MEH      HBBB   CBB    .       .
MEH      HACA   CAC    .       .
MEH      HBCB   CBC    .       .
MEH      C3D    C2D    .    ADD
MEH      C4D    CHA    .    ADD
MEH      ND     C1D    .    ADD
MEH      FE     NB     .    ADD
MEH      NB     C4B    .    ADD
MEH      C4B    C3B    .    ADD
MEH      NC     C4C    .    ADD
MEH      NA     C1A    .    ADD
loop_
_chem_comp_bond.comp_id
```

```
_chem_comp_bond.atom_id_1
_chem_comp_bond.atom_id_2
_chem_comp_bond.type
_chem_comp_bond.value_dist
_chem_comp_bond.value_dist_esd
MEH      C1A    CHA       deloc       1.393    0.020
MEH      CHA    C4D       deloc       1.393    0.020
MEH      C1B    CHB       deloc       1.393    0.020
MEH      CHB    C4A       deloc       1.393    0.020
MEH      C1C    CHC       deloc       1.393    0.020
MEH      CHC    C4B       deloc       1.393    0.020
MEH      C1D    CHD       deloc       1.393    0.020
MEH      CHD    C4C       deloc       1.393    0.020
MEH      CHA    HHA       single      1.077    0.020
MEH      CHB    HHB       single      1.077    0.020
MEH      CHC    HHC       single      1.077    0.020
MEH      CHD    HHD       single      1.077    0.020
MEH      CGD    O2D       deloc       1.250    0.020
MEH      CGD    O1D       deloc       1.250    0.020
MEH      CGD    CBD       single      1.510    0.020
MEH      CBD    HBD       single      1.092    0.020
MEH      CBD    HBDA      single      1.092    0.020
MEH      CBD    CAD       single      1.524    0.020
MEH      CAD    HAD       single      1.092    0.020
MEH      CAD    HADA      single      1.092    0.020
MEH      CAD    C3D       single      1.510    0.020
MEH      C3D    C2D       aromatic    1.390    0.020
MEH      C3D    C4D       aromatic    1.390    0.020
MEH      C2D    CMD       single      1.506    0.020
MEH      C2D    C1D       aromatic    1.390    0.020
MEH      CMD    HMDB      single      1.059    0.020
MEH      CMD    HMDA      single      1.059    0.020
MEH      CMD    HMD       single      1.059    0.020
MEH      C4D    ND        aromatic    1.337    0.020
MEH      ND     C1D       aromatic    1.337    0.020
MEH      NB     C4B       aromatic    1.337    0.020
MEH      NB     C1B       aromatic    1.337    0.020
MEH      C4B    C3B       aromatic    1.390    0.020
MEH      C3B    CAB       single      1.510    0.020
MEH      C3B    C2B       aromatic    1.390    0.020
MEH      CAB    HAB       single      1.092    0.020
MEH      CAB    CBB       single      1.513    0.020
MEH      CAB    HABA      single      1.092    0.020
MEH      CBB    HBBA      single      1.059    0.020
MEH      CBB    HBB       single      1.059    0.020
MEH      CBB    HBBB      single      1.059    0.020
MEH      C2B    CMB       single      1.506    0.020
MEH      C2B    C1B       aromatic    1.390    0.020
MEH      CMB    HMBB      single      1.059    0.020
MEH      CMB    HMBA      single      1.059    0.020
MEH      CMB    HMB       single      1.059    0.020
MEH      NC     C4C       aromatic    1.337    0.020
MEH      NC     C1C       aromatic    1.337    0.020
MEH      C4C    C3C       aromatic    1.390    0.020
MEH      C3C    CAC       single      1.510    0.020
MEH      C3C    C2C       aromatic    1.390    0.020
MEH      CAC    HAC       single      1.092    0.020
MEH      CAC    CBC       single      1.513    0.020
MEH      CAC    HACA      single      1.092    0.020
MEH      CBC    HBCA      single      1.059    0.020
MEH      CBC    HBC       single      1.059    0.020
```

```
MEH      CBC    HBCB      single      1.059    0.020
MEH      C2C    CMC       single      1.506    0.020
MEH      C2C    C1C       aromatic    1.390    0.020
MEH      CMC    HMCB      single      1.059    0.020
MEH      CMC    HMCA      single      1.059    0.020
MEH      CMC    HMC       single      1.059    0.020
MEH      NA     C1A       aromatic    1.337    0.020
MEH      NA     C4A       aromatic    1.337    0.020
MEH      C1A    C2A       aromatic    1.390    0.020
MEH      C4A    C3A       aromatic    1.390    0.020
MEH      C3A    CMA       single      1.506    0.020
MEH      C3A    C2A       aromatic    1.390    0.020
MEH      CMA    HMAB      single      1.059    0.020
MEH      CMA    HMAA      single      1.059    0.020
MEH      CMA    HMA       single      1.059    0.020
MEH      C2A    CAA       single      1.510    0.020
MEH      CAA    HAA       single      1.092    0.020
MEH      CAA    HAAA      single      1.092    0.020
MEH      CAA    CBA       single      1.524    0.020
MEH      CBA    HBA       single      1.092    0.020
MEH      CBA    HBAA      single      1.092    0.020
MEH      CBA    CGA       single      1.510    0.020
MEH      CGA    O1A       deloc       1.250    0.020
MEH      CGA    O2A       deloc       1.250    0.020
MEH      FE     NA        metal       2.090    0.100
MEH      FE     NB        metal       2.090    0.100
MEH      FE     NC        metal       2.090    0.100
MEH      FE     ND        metal       2.090    0.100
loop_
_chem_comp_angle.comp_id
_chem_comp_angle.atom_id_1
_chem_comp_angle.atom_id_2
_chem_comp_angle.atom_id_3
_chem_comp_angle.value_angle
_chem_comp_angle.value_angle_esd
MEH      O2D    CGD    O1D     123.000    3.000
MEH      O2D    CGD    CBD     118.500    3.000
MEH      CGD    CBD    HBD     109.470    3.000
MEH      CGD    CBD    HBDA    109.470    3.000
MEH      CGD    CBD    CAD     109.470    3.000
MEH      O1D    CGD    CBD     118.500    3.000
MEH      CBD    CAD    HAD     109.470    3.000
MEH      CBD    CAD    HADA    109.470    3.000
MEH      CBD    CAD    C3D     109.470    3.000
MEH      HBD    CBD    HBDA    107.900    3.000
MEH      HBD    CBD    CAD     109.470    3.000
MEH      HBDA   CBD    CAD     109.470    3.000
MEH      CAD    C3D    C2D     126.000    3.000
MEH      CAD    C3D    C4D     126.000    3.000
MEH      HAD    CAD    HADA    107.900    3.000
MEH      HAD    CAD    C3D     109.470    3.000
MEH      HADA   CAD    C3D     109.470    3.000
MEH      C3D    C2D    CMD     126.000    3.000
MEH      C3D    C2D    C1D     108.000    3.000
MEH      C3D    C4D    CHA     117.000    3.000
MEH      C3D    C4D    ND      108.000    3.000
MEH      C2D    C3D    C4D     108.000    3.000
MEH      C2D    CMD    HMDB    109.470    3.000
MEH      C2D    CMD    HMDA    109.470    3.000
MEH      C2D    CMD    HMD     109.470    3.000
MEH      C2D    C1D    ND      108.000    3.000
```

| | | | | | |
|---|---|---|---|---|---|
| MEH | C2D | C1D | CHD | 117.000 | 3.000 |
| MEH | CMD | C2D | C1D | 126.000 | 3.000 |
| MEH | HMDB | CMD | HMDA | 109.470 | 3.000 |
| MEH | HMDB | CMD | HMD | 109.470 | 3.000 |
| MEH | HMDA | CMD | HMD | 109.470 | 3.000 |
| MEH | C4D | CHA | HHA | 120.000 | 3.000 |
| MEH | C4D | CHA | C1A | 120.000 | 3.000 |
| MEH | C4D | ND | C1D | 108.000 | 3.000 |
| MEH | CHA | C4D | ND | 108.000 | 3.000 |
| MEH | CHA | C1A | NA | 108.000 | 3.000 |
| MEH | CHA | C1A | C2A | 117.000 | 3.000 |
| MEH | HHA | CHA | C1A | 120.000 | 3.000 |
| MEH | ND | C1D | CHD | 108.000 | 3.000 |
| MEH | C1D | CHD | HHD | 120.000 | 3.000 |
| MEH | C1D | CHD | C4C | 120.000 | 3.000 |
| MEH | CHD | C4C | NC | 108.000 | 3.000 |
| MEH | CHD | C4C | C3C | 117.000 | 3.000 |
| MEH | HHD | CHD | C4C | 120.000 | 3.000 |
| MEH | NB | C4B | C3B | 108.000 | 3.000 |
| MEH | NB | C4B | CHC | 108.000 | 3.000 |
| MEH | NB | C1B | C2B | 108.000 | 3.000 |
| MEH | NB | C1B | CHB | 108.000 | 3.000 |
| MEH | C4B | NB | C1B | 108.000 | 3.000 |
| MEH | C4B | C3B | CAB | 126.000 | 3.000 |
| MEH | C4B | C3B | C2B | 108.000 | 3.000 |
| MEH | C4B | CHC | C1C | 120.000 | 3.000 |
| MEH | C4B | CHC | HHC | 120.000 | 3.000 |
| MEH | C3B | C4B | CHC | 117.000 | 3.000 |
| MEH | C3B | CAB | HAB | 109.470 | 3.000 |
| MEH | C3B | CAB | CBB | 109.470 | 3.000 |
| MEH | C3B | CAB | HABA | 109.470 | 3.000 |
| MEH | C3B | C2B | CMB | 126.000 | 3.000 |
| MEH | C3B | C2B | C1B | 108.000 | 3.000 |
| MEH | CAB | C3B | C2B | 126.000 | 3.000 |
| MEH | CAB | CBB | HBBA | 109.470 | 3.000 |
| MEH | CAB | CBB | HBB | 109.470 | 3.000 |
| MEH | CAB | CBB | HBBB | 109.470 | 3.000 |
| MEH | HAB | CAB | CBB | 109.470 | 3.000 |
| MEH | HAB | CAB | HABA | 107.900 | 3.000 |
| MEH | CBB | CAB | HABA | 109.470 | 3.000 |
| MEH | HBBA | CBB | HBB | 109.470 | 3.000 |
| MEH | HBBA | CBB | HBBB | 109.470 | 3.000 |
| MEH | HBB | CBB | HBBB | 109.470 | 3.000 |
| MEH | C2B | CMB | HMBB | 109.470 | 3.000 |
| MEH | C2B | CMB | HMBA | 109.470 | 3.000 |
| MEH | C2B | CMB | HMB | 109.470 | 3.000 |
| MEH | C2B | C1B | CHB | 117.000 | 3.000 |
| MEH | CMB | C2B | C1B | 126.000 | 3.000 |
| MEH | HMBB | CMB | HMBA | 109.470 | 3.000 |
| MEH | HMBB | CMB | HMB | 109.470 | 3.000 |
| MEH | HMBA | CMB | HMB | 109.470 | 3.000 |
| MEH | C1B | CHB | HHB | 120.000 | 3.000 |
| MEH | C1B | CHB | C4A | 120.000 | 3.000 |
| MEH | CHB | C4A | NA | 108.000 | 3.000 |
| MEH | CHB | C4A | C3A | 117.000 | 3.000 |
| MEH | HHB | CHB | C4A | 120.000 | 3.000 |
| MEH | NC | C4C | C3C | 108.000 | 3.000 |
| MEH | NC | C1C | C2C | 108.000 | 3.000 |
| MEH | NC | C1C | CHC | 108.000 | 3.000 |
| MEH | C4C | NC | C1C | 108.000 | 3.000 |
| MEH | C4C | C3C | CAC | 126.000 | 3.000 |

| | | | | | |
|---|---|---|---|---|---|
| MEH | C4C | C3C | C2C | 108.000 | 3.000 |
| MEH | C3C | CAC | HAC | 109.470 | 3.000 |
| MEH | C3C | CAC | CBC | 109.470 | 3.000 |
| MEH | C3C | CAC | HACA | 109.470 | 3.000 |
| MEH | C3C | C2C | CMC | 126.000 | 3.000 |
| MEH | C3C | C2C | C1C | 108.000 | 3.000 |
| MEH | CAC | C3C | C2C | 126.000 | 3.000 |
| MEH | CAC | CBC | HBCA | 109.470 | 3.000 |
| MEH | CAC | CBC | HBC | 109.470 | 3.000 |
| MEH | CAC | CBC | HBCB | 109.470 | 3.000 |
| MEH | HAC | CAC | CBC | 109.470 | 3.000 |
| MEH | HAC | CAC | HACA | 107.900 | 3.000 |
| MEH | CBC | CAC | HACA | 109.470 | 3.000 |
| MEH | HBCA | CBC | HBC | 109.470 | 3.000 |
| MEH | HBCA | CBC | HBCB | 109.470 | 3.000 |
| MEH | HBC | CBC | HBCB | 109.470 | 3.000 |
| MEH | C2C | CMC | HMCB | 109.470 | 3.000 |
| MEH | C2C | CMC | HMCA | 109.470 | 3.000 |
| MEH | C2C | CMC | HMC | 109.470 | 3.000 |
| MEH | C2C | C1C | CHC | 117.000 | 3.000 |
| MEH | CMC | C2C | C1C | 126.000 | 3.000 |
| MEH | HMCB | CMC | HMCA | 109.470 | 3.000 |
| MEH | HMCB | CMC | HMC | 109.470 | 3.000 |
| MEH | HMCA | CMC | HMC | 109.470 | 3.000 |
| MEH | C1C | CHC | HHC | 120.000 | 3.000 |
| MEH | NA | C1A | C2A | 108.000 | 3.000 |
| MEH | NA | C4A | C3A | 108.000 | 3.000 |
| MEH | C1A | NA | C4A | 108.000 | 3.000 |
| MEH | C1A | C2A | C3A | 108.000 | 3.000 |
| MEH | C1A | C2A | CAA | 126.000 | 3.000 |
| MEH | C4A | C3A | CMA | 126.000 | 3.000 |
| MEH | C4A | C3A | C2A | 108.000 | 3.000 |
| MEH | C3A | CMA | HMAB | 109.470 | 3.000 |
| MEH | C3A | CMA | HMAA | 109.470 | 3.000 |
| MEH | C3A | CMA | HMA | 109.470 | 3.000 |
| MEH | C3A | C2A | CAA | 126.000 | 3.000 |
| MEH | CMA | C3A | C2A | 126.000 | 3.000 |
| MEH | HMAB | CMA | HMAA | 109.470 | 3.000 |
| MEH | HMAB | CMA | HMA | 109.470 | 3.000 |
| MEH | HMAA | CMA | HMA | 109.470 | 3.000 |
| MEH | C2A | CAA | HAA | 109.470 | 3.000 |
| MEH | C2A | CAA | HAAA | 109.470 | 3.000 |
| MEH | C2A | CAA | CBA | 109.470 | 3.000 |
| MEH | CAA | CBA | HBA | 109.470 | 3.000 |
| MEH | CAA | CBA | HBAA | 109.470 | 3.000 |
| MEH | CAA | CBA | CGA | 109.470 | 3.000 |
| MEH | HAA | CAA | HAAA | 107.900 | 3.000 |
| MEH | HAA | CAA | CBA | 109.470 | 3.000 |
| MEH | HAAA | CAA | CBA | 109.470 | 3.000 |
| MEH | CBA | CGA | O1A | 118.500 | 3.000 |
| MEH | CBA | CGA | O2A | 118.500 | 3.000 |
| MEH | HBA | CBA | HBAA | 107.900 | 3.000 |
| MEH | HBA | CBA | CGA | 109.470 | 3.000 |
| MEH | HBAA | CBA | CGA | 109.470 | 3.000 |
| MEH | O1A | CGA | O2A | 123.000 | 3.000 |
| MEH | FE | NA | C1A | 126.000 | 7.500 |
| MEH | FE | NA | C4A | 126.000 | 7.500 |
| MEH | FE | NB | C1B | 126.000 | 7.500 |
| MEH | FE | NB | C4B | 126.000 | 7.500 |
| MEH | FE | NC | C1C | 126.000 | 7.500 |
| MEH | FE | NC | C4C | 126.000 | 7.500 |

```
MEH      FE     ND     C1D     126.000    7.500
MEH      FE     ND     C4D     126.000    7.500
MEH      NA     FE     NB       90.000    7.500
MEH      NB     FE     NC       90.000    7.500
MEH      NC     FE     ND       90.000    7.500
MEH      ND     FE     NA       90.000    7.500
loop_
_chem_comp_plane_atom.comp_id
_chem_comp_plane_atom.plane_id
_chem_comp_plane_atom.atom_id
_chem_comp_plane_atom.dist_esd
MEH      plan-1A          C4D       0.020
MEH      plan-1A          CHA       0.020
MEH      plan-1A          HHA       0.020
MEH      plan-1A          C1A       0.020
MEH      plan-2A          CHA       0.020
MEH      plan-2A          C1A       0.020
MEH      plan-2A          C2A       0.020
MEH      plan-2A          NA        0.020
MEH      plan-3A          NA        0.020
MEH      plan-3A          C1A       0.020
MEH      plan-3A          C2A       0.020
MEH      plan-3A          C3A       0.020
MEH      plan-3A          C4A       0.020
MEH      plan-4A          C1A       0.020
MEH      plan-4A          C2A       0.020
MEH      plan-4A          CAA       0.020
MEH      plan-4A          C3A       0.020
MEH      plan-5A          C2A       0.020
MEH      plan-5A          C3A       0.020
MEH      plan-5A          CMA       0.020
MEH      plan-5A          C4A       0.020
MEH      plan-6A          CBA       0.020
MEH      plan-6A          CGA       0.020
MEH      plan-6A          O1A       0.020
MEH      plan-6A          O2A       0.020
MEH      plan-7A          NA        0.020
MEH      plan-7A          C4A       0.020
MEH      plan-7A          C3A       0.020
MEH      plan-7A          CHB       0.020
MEH      plan-1B          C4A       0.020
MEH      plan-1B          CHB       0.020
MEH      plan-1B          HHB       0.020
MEH      plan-1B          C1B       0.020
MEH      plan-2B          CHB       0.020
MEH      plan-2B          C1B       0.020
MEH      plan-2B          C2B       0.020
MEH      plan-2B          NB        0.020
MEH      plan-3B          NB        0.020
MEH      plan-3B          C1B       0.020
MEH      plan-3B          C2B       0.020
MEH      plan-3B          C3B       0.020
MEH      plan-3B          C4B       0.020
MEH      plan-4B          C1B       0.020
MEH      plan-4B          C2B       0.020
MEH      plan-4B          CMB       0.020
MEH      plan-4B          C3B       0.020
MEH      plan-5B          C2B       0.020
MEH      plan-5B          C3B       0.020
MEH      plan-5B          CAB       0.020
MEH      plan-5B          C4B       0.020
```

```
MEH      plan-7B          NB        0.020
MEH      plan-7B          C4B       0.020
MEH      plan-7B          C3B       0.020
MEH      plan-7B          CHC       0.020
MEH      plan-1C          C4B       0.020
MEH      plan-1C          CHC       0.020
MEH      plan-1C          HHC       0.020
MEH      plan-1C          C1C       0.020
MEH      plan-2C          CHC       0.020
MEH      plan-2C          C1C       0.020
MEH      plan-2C          C2C       0.020
MEH      plan-2C          NC        0.020
MEH      plan-3C          NC        0.020
MEH      plan-3C          C1C       0.020
MEH      plan-3C          C2C       0.020
MEH      plan-3C          C3C       0.020
MEH      plan-3C          C4C       0.020
MEH      plan-4C          C1C       0.020
MEH      plan-4C          C2C       0.020
MEH      plan-4C          CMC       0.020
MEH      plan-4C          C3C       0.020
MEH      plan-5C          C2C       0.020
MEH      plan-5C          C3C       0.020
MEH      plan-5C          CAC       0.020
MEH      plan-5C          C4C       0.020
MEH      plan-7C          NC        0.020
MEH      plan-7C          C4C       0.020
MEH      plan-7C          C3C       0.020
MEH      plan-7C          CHD       0.020
MEH      plan-1D          C4C       0.020
MEH      plan-1D          CHD       0.020
MEH      plan-1D          HHD       0.020
MEH      plan-1D          C1D       0.020
MEH      plan-2D          CHD       0.020
MEH      plan-2D          C1D       0.020
MEH      plan-2D          C2D       0.020
MEH      plan-2D          ND        0.020
MEH      plan-3D          ND        0.020
MEH      plan-3D          C1D       0.020
MEH      plan-3D          C2D       0.020
MEH      plan-3D          C3D       0.020
MEH      plan-3D          C4D       0.020
MEH      plan-4D          C1D       0.020
MEH      plan-4D          C2D       0.020
MEH      plan-4D          CMD       0.020
MEH      plan-4D          C3D       0.020
MEH      plan-5D          C2D       0.020
MEH      plan-5D          C3D       0.020
MEH      plan-5D          CAD       0.020
MEH      plan-5D          C4D       0.020
MEH      plan-6D          CBD       0.020
MEH      plan-6D          CGD       0.020
MEH      plan-6D          O1D       0.020
MEH      plan-6D          O2D       0.020
MEH      plan-7D          ND        0.020
MEH      plan-7D          C4D       0.020
MEH      plan-7D          C3D       0.020
MEH      plan-7D          CHA       0.020
#
data_comp_No
#
```

```
loop_
_chem_comp_atom.comp_id
_chem_comp_atom.atom_id
_chem_comp_atom.type_symbol
_chem_comp_atom.type_energy
_chem_comp_atom.partial_charge
No             N      N    N         0.000
No             O      O    O         0.000
loop_
_chem_comp_tree.comp_id
_chem_comp_tree.atom_id
_chem_comp_tree.atom_back
_chem_comp_tree.atom_forward
_chem_comp_tree.connect_type
No       N      n/a    O      START
No       O      N      .      END
loop_
_chem_comp_bond.comp_id
_chem_comp_bond.atom_id_1
_chem_comp_bond.atom_id_2
_chem_comp_bond.type
_chem_comp_bond.value_dist
_chem_comp_bond.value_dist_esd
No       O      N         triple      1.15    0.020
#loop_
# ------------------------------------------------------
#
# --- DESCRIPTION OF MODIFICATIONS ---
#
data_mod_CYS-HG
#
loop_
_chem_mod_atom.mod_id
_chem_mod_atom.function
_chem_mod_atom.atom_id
_chem_mod_atom.new_atom_id
_chem_mod_atom.new_type_symbol
_chem_mod_atom.new_type_energy
_chem_mod_atom.new_partial_charge
CYS-HG   change   SG     .      .    S2      0.000
CYS-HG   delete   HG     .      .    .       0.000
#
data_mod_MEH-HABA
#
loop_
_chem_mod_atom.mod_id
_chem_mod_atom.function
_chem_mod_atom.atom_id
_chem_mod_atom.new_atom_id
_chem_mod_atom.new_type_symbol
_chem_mod_atom.new_type_energy
_chem_mod_atom.new_partial_charge
MEH-HABA change   CAB    .      .    CH1     0.000
MEH-HABA delete   HABA   .      .    .       0.000
loop_
_chem_mod_bond.mod_id
_chem_mod_bond.function
_chem_mod_bond.atom_id_1
_chem_mod_bond.atom_id_2
_chem_mod_bond.new_type
_chem_mod_bond.new_value_dist
```

```
_chem_mod_bond.new_value_dist_esd
MEH-HABA change   C3B     CAB       .           1.480    0.020
#
data_mod_MEH-HACA
#
loop_
_chem_mod_atom.mod_id
_chem_mod_atom.function
_chem_mod_atom.atom_id
_chem_mod_atom.new_atom_id
_chem_mod_atom.new_type_symbol
_chem_mod_atom.new_type_energy
_chem_mod_atom.new_partial_charge
MEH-HACA change   CAC    .      .    CH1     0.000
MEH-HACA delete   HACA   .      .    .       0.000
loop_
_chem_mod_bond.mod_id
_chem_mod_bond.function
_chem_mod_bond.atom_id_1
_chem_mod_bond.atom_id_2
_chem_mod_bond.new_type
_chem_mod_bond.new_value_dist
_chem_mod_bond.new_value_dist_esd
MEH-HACA change   C3C     CAC       .           1.480    0.020
#
data_mod_HIS-HE2
#
loop_
_chem_mod_atom.mod_id
_chem_mod_atom.function
_chem_mod_atom.atom_id
_chem_mod_atom.new_atom_id
_chem_mod_atom.new_type_symbol
_chem_mod_atom.new_type_energy
_chem_mod_atom.new_partial_charge
HIS-HE2  delete   HE2    .      .    .       0.000
# ------------------------------------------------------
#
# --- DESCRIPTION OF LINKS ---
#
data_link_CYS-MEHB
#
loop_
_chem_link_bond.link_id
_chem_link_bond.atom_1_comp_id
_chem_link_bond.atom_id_1
_chem_link_bond.atom_2_comp_id
_chem_link_bond.atom_id_2
_chem_link_bond.type
_chem_link_bond.value_dist
_chem_link_bond.value_dist_esd
CYS-MEHB 1 SG      2 CAB       single      1.765    0.020
loop_
_chem_link_angle.link_id
_chem_link_angle.atom_1_comp_id
_chem_link_angle.atom_id_1
_chem_link_angle.atom_2_comp_id
_chem_link_angle.atom_id_2
_chem_link_angle.atom_3_comp_id
_chem_link_angle.atom_id_3
_chem_link_angle.value_angle
```

```
_chem_link_angle.value_angle_esd
CYS-MEHB 2 CAB     1 SG      1 CB      109.470    3.000
CYS-MEHB 2 HAB     2 CAB     1 SG      109.500    3.000
CYS-MEHB 2 CBB     2 CAB     1 SG      109.500    3.000
CYS-MEHB 2 C3B     2 CAB     1 SG      109.500    3.000
loop_
_chem_link_chir.link_id
_chem_link_chir.atom_centre_comp_id
_chem_link_chir.atom_id_centre
_chem_link_chir.atom_1_comp_id
_chem_link_chir.atom_id_1
_chem_link_chir.atom_2_comp_id
_chem_link_chir.atom_id_2
_chem_link_chir.atom_3_comp_id
_chem_link_chir.atom_id_3
_chem_link_chir.volume_sign
CYS-MEHB   2 CAB     2 C3B     2 CBB     1 SG      both
#
data_link_CYS-MEHC
#
loop_
_chem_link_bond.link_id
_chem_link_bond.atom_1_comp_id
_chem_link_bond.atom_id_1
_chem_link_bond.atom_2_comp_id
_chem_link_bond.atom_id_2
_chem_link_bond.type
_chem_link_bond.value_dist
_chem_link_bond.value_dist_esd
CYS-MEHC 1 SG      2 CAC       single      1.765    0.020
loop_
_chem_link_angle.link_id
_chem_link_angle.atom_1_comp_id
_chem_link_angle.atom_id_1
_chem_link_angle.atom_2_comp_id
_chem_link_angle.atom_id_2
_chem_link_angle.atom_3_comp_id
_chem_link_angle.atom_id_3
_chem_link_angle.value_angle
_chem_link_angle.value_angle_esd
CYS-MEHC 2 CAC     1 SG      1 CB      109.470    3.000
CYS-MEHC 2 HAC     2 CAC     1 SG      109.500    3.000
CYS-MEHC 2 CBC     2 CAC     1 SG      109.500    3.000
CYS-MEHC 2 C3C     2 CAC     1 SG      109.500    3.000
loop_
_chem_link_chir.link_id
_chem_link_chir.atom_centre_comp_id
_chem_link_chir.atom_id_centre
_chem_link_chir.atom_1_comp_id
_chem_link_chir.atom_id_1
_chem_link_chir.atom_2_comp_id
_chem_link_chir.atom_id_2
_chem_link_chir.atom_3_comp_id
_chem_link_chir.atom_id_3
_chem_link_chir.volume_sign
CYS-MEHC   2 CAC     2 C3C     2 CBC     1 SG      both
#
data_link_HIS-MEH
#
loop_
_chem_link_bond.link_id
```

```
_chem_link_bond.atom_1_comp_id
_chem_link_bond.atom_id_1
_chem_link_bond.atom_2_comp_id
_chem_link_bond.atom_id_2
_chem_link_bond.type
_chem_link_bond.value_dist
_chem_link_bond.value_dist_esd
HIS-MEH  1 NE2     2 FE        metal       1.935    0.120
# ------------------------------------------------------
```

### HX2 restraints

Restraints used for the RT fresh and RT aged structures. The restraints were made in JLigand based on the SMILES string of the proposed heme-lysine structure. The heme and Lys78 monomers were deleted in Coot and replaced by the HX2 monomer. The iron atom (Fe201) was added back in.

```
data_xyz

loop_
_struct_conn.id
_struct_conn.conn_type_id
_struct_conn.ptnr1_label_asym_id
_struct_conn.ptnr1_label_seq_id
_struct_conn.ptnr1_label_comp_id
_struct_conn.ptnr1_label_atom_id
_struct_conn.ptnr2_label_asym_id
_struct_conn.ptnr2_label_seq_id
_struct_conn.ptnr2_label_comp_id
_struct_conn.ptnr2_label_atom_id
_struct_conn.ccp4_link_id
   1 covalent A    3 HX2 .     A    5 CYS .     HX2-CYS
   2 covalent A    9 CYS .     A    11 HX2 .     CYS-HX2

loop_
_atom_site.id
_atom_site.type_symbol
_atom_site.label_atom_id
_atom_site.label_alt_id
_atom_site.label_comp_id
_atom_site.label_asym_id
_atom_site.auth_seq_id
_atom_site.Cartn_x
_atom_site.Cartn_y
_atom_site.Cartn_z
_atom_site.occupancy
_atom_site.B_iso_or_equiv
   1 C   CBC   . HX2 A   1    -6.641    -7.573     0.719  1.00 .
   2 C   CAC   . HX2 A   1    -7.081    -8.936     0.230  1.00 .
   3 S   S1    . HX2 A   1    -8.068    -9.810     1.479  1.00 .
   4 C   C3C   . HX2 A   1    -5.909    -9.768    -0.200  1.00 .
   5 C   C4C   . HX2 A   1    -5.630   -11.153     0.091  1.00 .
   6 N   NC    . HX2 A   1    -4.437   -11.454    -0.515  1.00 .
   7 C   C1C   . HX2 A   1    -3.950   -10.365    -1.191  1.00 .
   8 C   CHC   . HX2 A   1    -2.748   -10.327    -1.875  1.00 .
   9 C   C4B   . HX2 A   1    -1.533   -10.269    -1.260  1.00 .
  10 N   NB    . HX2 A   1    -0.458   -11.076    -1.501  1.00 .
  11 C   C1B   . HX2 A   1     0.117   -11.139    -0.261  1.00 .
  12 C   CHB   . HX2 A   1     0.649   -12.275     0.323  1.00 .
  13 C   C4A   . HX2 A   1    -0.030   -13.482     0.375  1.00 .
```

| | | | | | | | | | | | |
|---|---|---|---|---|---|---|---|---|---|---|---|
| 14 | N | NA | . | HX2 | A | 1 | -1.298 | -13.719 | 0.914 | 1.00 | . |
| 15 | C | C1A | . | HX2 | A | 1 | -1.933 | -14.576 | 0.071 | 1.00 | . |
| 16 | C | CHA | . | HX2 | A | 1 | -3.236 | -14.657 | -0.431 | 1.00 | . |
| 17 | N | NZ | . | HX2 | A | 1 | -3.262 | -14.679 | -1.782 | 1.00 | . |
| 18 | C | CE | . | HX2 | A | 1 | -2.139 | -14.930 | -2.493 | 1.00 | . |
| 19 | C | CD | . | HX2 | A | 1 | -1.145 | -15.674 | -1.924 | 1.00 | . |
| 20 | C | C2A | . | HX2 | A | 1 | -0.855 | -15.513 | -0.446 | 1.00 | . |
| 21 | C | CAA | . | HX2 | A | 1 | -0.808 | -16.870 | 0.292 | 1.00 | . |
| 22 | C | CBA | . | HX2 | A | 1 | 0.572 | -17.295 | 0.774 | 1.00 | . |
| 23 | C | CGA | . | HX2 | A | 1 | 1.609 | -17.386 | -0.331 | 1.00 | . |
| 24 | O | O2A | . | HX2 | A | 1 | 2.377 | -18.370 | -0.338 | 1.00 | . |
| 25 | O | O1A | . | HX2 | A | 1 | 1.648 | -16.472 | -1.181 | 1.00 | . |
| 26 | C | C3A | . | HX2 | A | 1 | 0.398 | -14.697 | -0.132 | 1.00 | . |
| 27 | C | CMA | . | HX2 | A | 1 | 1.641 | -15.100 | -0.319 | 1.00 | . |
| 28 | C | C4D | . | HX2 | A | 1 | -4.485 | -14.675 | 0.334 | 1.00 | . |
| 29 | N | ND | . | HX2 | A | 1 | -5.643 | -14.166 | -0.175 | 1.00 | . |
| 30 | C | C1D | . | HX2 | A | 1 | -6.043 | -13.391 | 0.876 | 1.00 | . |
| 31 | C | CHD | . | HX2 | A | 1 | -6.366 | -12.067 | 0.831 | 1.00 | . |
| 32 | C | C2D | . | HX2 | A | 1 | -5.823 | -14.207 | 2.072 | 1.00 | . |
| 33 | C | CMD | . | HX2 | A | 1 | -6.525 | -14.092 | 3.394 | 1.00 | . |
| 34 | C | C3D | . | HX2 | A | 1 | -4.846 | -15.078 | 1.697 | 1.00 | . |
| 35 | C | CAD | . | HX2 | A | 1 | -4.292 | -16.224 | 2.484 | 1.00 | . |
| 36 | C | CBD | . | HX2 | A | 1 | -4.746 | -17.656 | 2.253 | 1.00 | . |
| 37 | C | CGD | . | HX2 | A | 1 | -3.670 | -18.676 | 2.566 | 1.00 | . |
| 38 | O | O1D | . | HX2 | A | 1 | -3.193 | -18.691 | 3.720 | 1.00 | . |
| 39 | O | O2D | . | HX2 | A | 1 | -3.314 | -19.452 | 1.654 | 1.00 | . |
| 40 | C | C2B | . | HX2 | A | 1 | -0.044 | -9.818 | 0.324 | 1.00 | . |
| 41 | C | CMB | . | HX2 | A | 1 | 0.900 | -9.109 | 1.250 | 1.00 | . |
| 42 | C | C3B | . | HX2 | A | 1 | -1.225 | -9.379 | -0.160 | 1.00 | . |
| 43 | C | CAB | . | HX2 | A | 1 | -2.097 | -8.254 | 0.325 | 1.00 | . |
| 44 | C | CBB | . | HX2 | A | 1 | -1.883 | -7.984 | 1.797 | 1.00 | . |
| 45 | S | S2 | . | HX2 | A | 1 | -3.854 | -8.546 | -0.033 | 1.00 | . |
| 46 | C | C2C | . | HX2 | A | 1 | -4.874 | -9.290 | -0.977 | 1.00 | . |
| 47 | C | CMC | . | HX2 | A | 1 | -4.720 | -7.900 | -1.520 | 1.00 | . |
| 48 | H | H1 | . | HX2 | A | 1 | -5.862 | -7.294 | 0.231 | 1.00 | . |
| 49 | H | H21 | . | HX2 | A | 1 | -7.349 | -6.940 | 0.584 | 1.00 | . |
| 50 | H | H31 | . | HX2 | A | 1 | -6.430 | -7.623 | 1.655 | 1.00 | . |
| 51 | H | H4 | . | HX2 | A | 1 | -7.662 | -8.800 | -0.561 | 1.00 | . |
| 52 | H | H5 | . | HX2 | A | 1 | -7.279 | -9.790 | 2.399 | 1.00 | . |
| 53 | H | H6 | . | HX2 | A | 1 | -4.048 | -12.241 | -0.490 | 1.00 | . |
| 54 | H | H7 | . | HX2 | A | 1 | -2.764 | -10.326 | -2.827 | 1.00 | . |
| 55 | H | H8 | . | HX2 | A | 1 | 1.524 | -12.219 | 0.701 | 1.00 | . |
| 56 | H | H9 | . | HX2 | A | 1 | -1.633 | -13.385 | 1.654 | 1.00 | . |
| 57 | H | H10 | . | HX2 | A | 1 | -4.029 | -14.526 | -2.207 | 1.00 | . |
| 58 | H | H11 | . | HX2 | A | 1 | -2.033 | -14.601 | -3.369 | 1.00 | . |
| 59 | H | H13 | . | HX2 | A | 1 | -1.158 | -17.563 | -0.307 | 1.00 | . |
| 60 | H | H14 | . | HX2 | A | 1 | -1.409 | -16.826 | 1.065 | 1.00 | . |
| 61 | H | H15 | . | HX2 | A | 1 | 0.883 | -16.651 | 1.445 | 1.00 | . |
| 62 | H | H16 | . | HX2 | A | 1 | 0.497 | -18.172 | 1.207 | 1.00 | . |
| 63 | H | H17 | . | HX2 | A | 1 | 2.362 | -14.530 | -0.095 | 1.00 | . |
| 64 | H | H18 | . | HX2 | A | 1 | 1.810 | -15.958 | -0.676 | 1.00 | . |
| 65 | H | H19 | . | HX2 | A | 1 | -7.112 | -11.768 | 1.345 | 1.00 | . |
| 66 | H | H20 | . | HX2 | A | 1 | -7.472 | -13.941 | 3.247 | 1.00 | . |
| 67 | H | H211 | . | HX2 | A | 1 | -6.405 | -14.911 | 3.899 | 1.00 | . |
| 68 | H | H22 | . | HX2 | A | 1 | -6.156 | -13.350 | 3.896 | 1.00 | . |
| 69 | H | H23 | . | HX2 | A | 1 | -3.999 | -16.415 | 1.571 | 1.00 | . |
| 70 | H | H24 | . | HX2 | A | 1 | -5.235 | -16.211 | 2.740 | 1.00 | . |
| 71 | H | H25 | . | HX2 | A | 1 | -5.529 | -17.853 | 2.811 | 1.00 | . |
| 72 | H | H26 | . | HX2 | A | 1 | -5.020 | -17.773 | 1.319 | 1.00 | . |
| 73 | H | H27 | . | HX2 | A | 1 | 0.420 | -8.425 | 1.741 | 1.00 | . |
| 74 | H | H28 | . | HX2 | A | 1 | 1.284 | -9.744 | 1.873 | 1.00 | . |

| | | | | | | | | | | | |
|---|---|---|---|---|---|---|---|---|---|---|---|
| 75 | H | H29 | . | HX2 | A | 1 | 1.610 | -8.695 | 0.735 | 1.00 | . |
| 76 | H | H30 | . | HX2 | A | 1 | -1.829 | -7.443 | -0.175 | 1.00 | . |
| 77 | H | H311 | . | HX2 | A | 1 | -1.654 | -7.170 | 2.278 | 1.00 | . |
| 78 | H | H32 | . | HX2 | A | 1 | -2.699 | -7.474 | 1.655 | 1.00 | . |
| 79 | H | H33 | . | HX2 | A | 1 | -0.975 | -7.853 | 1.474 | 1.00 | . |
| 80 | H | H34 | . | HX2 | A | 1 | -4.093 | -9.342 | 0.852 | 1.00 | . |
| 81 | H | H35 | . | HX2 | A | 1 | -5.512 | -7.379 | -1.313 | 1.00 | . |
| 82 | H | H36 | . | HX2 | A | 1 | -3.943 | -7.478 | -1.118 | 1.00 | . |
| 83 | H | H37 | . | HX2 | A | 1 | -4.602 | -7.938 | -2.483 | 1.00 | . |
| 84 | N | N | . | HX2 | A | 1 | -2.370 | -19.884 | -2.666 | 1.00 | . |
| 85 | C | CA | . | HX2 | A | 1 | -1.083 | -19.140 | -2.702 | 1.00 | . |
| 86 | C | C | . | HX2 | A | 1 | -0.528 | -19.017 | -1.274 | 1.00 | . |
| 87 | O | O | . | HX2 | A | 1 | 0.650 | -19.394 | -1.084 | 1.00 | . |
| 88 | C | CB | . | HX2 | A | 1 | -1.283 | -17.760 | -3.329 | 1.00 | . |
| 89 | C | CG | . | HX2 | A | 1 | -0.379 | -16.685 | -2.741 | 1.00 | . |
| 90 | O | OXT | . | HX2 | A | 1 | -1.293 | -18.549 | -0.402 | 1.00 | . |
| 91 | H | H | . | HX2 | A | 1 | -2.463 | -20.297 | -1.869 | 1.00 | . |
| 92 | H | H2 | . | HX2 | A | 1 | -2.386 | -20.503 | -3.321 | 1.00 | . |
| 93 | H | H3 | . | HX2 | A | 1 | -3.060 | -19.313 | -2.783 | 1.00 | . |
| 94 | H | HA | . | HX2 | A | 1 | -0.440 | -19.656 | -3.251 | 1.00 | . |
| 95 | H | HB3 | . | HX2 | A | 1 | -2.217 | -17.491 | -3.201 | 1.00 | . |
| 96 | H | HB2 | . | HX2 | A | 1 | -1.116 | -17.825 | -4.292 | 1.00 | . |
| 97 | H | HG3 | . | HX2 | A | 1 | 0.300 | -17.114 | -2.168 | 1.00 | . |
| 98 | H | HG2 | . | HX2 | A | 1 | 0.085 | -16.218 | -3.477 | 1.00 | . |
| 99 | C | CBC | . | HX2 | A | 3 | -16.842 | 2.710 | -1.317 | 1.00 | . |
| 100 | C | CAC | . | HX2 | A | 3 | -17.335 | 1.730 | -0.258 | 1.00 | . |
| 101 | C | C3C | . | HX2 | A | 3 | -17.246 | 0.305 | -0.729 | 1.00 | . |
| 102 | C | C4C | . | HX2 | A | 3 | -16.130 | -0.225 | -1.456 | 1.00 | . |
| 103 | N | NC | . | HX2 | A | 3 | -16.399 | -1.550 | -1.693 | 1.00 | . |
| 104 | C | C1C | . | HX2 | A | 3 | -17.628 | -1.888 | -1.180 | 1.00 | . |
| 105 | C | CHC | . | HX2 | A | 3 | -18.199 | -3.152 | -1.259 | 1.00 | . |
| 106 | C | C4B | . | HX2 | A | 3 | -17.870 | -4.129 | -2.156 | 1.00 | . |
| 107 | N | NB | . | HX2 | A | 3 | -16.983 | -5.140 | -1.936 | 1.00 | . |
| 108 | C | C1B | . | HX2 | A | 3 | -16.232 | -5.002 | -3.070 | 1.00 | . |
| 109 | C | CHB | . | HX2 | A | 3 | -14.851 | -4.947 | -3.112 | 1.00 | . |
| 110 | C | C4A | . | HX2 | A | 3 | -14.106 | -4.606 | -1.999 | 1.00 | . |
| 111 | N | NA | . | HX2 | A | 3 | -12.824 | -4.046 | -1.955 | 1.00 | . |
| 112 | C | C1A | . | HX2 | A | 3 | -12.621 | -3.617 | -0.682 | 1.00 | . |
| 113 | C | CHA | . | HX2 | A | 3 | -12.048 | -2.497 | -0.074 | 1.00 | . |
| 114 | N | NZ | . | HX2 | A | 3 | -11.426 | -2.799 | 1.085 | 1.00 | . |
| 115 | C | CE | . | HX2 | A | 3 | -12.176 | -3.446 | 2.004 | 1.00 | . |
| 116 | C | CD | . | HX2 | A | 3 | -13.361 | -3.967 | 1.561 | 1.00 | . |
| 117 | C | C2A | . | HX2 | A | 3 | -13.317 | -4.638 | 0.199 | 1.00 | . |
| 118 | C | CAA | . | HX2 | A | 3 | -12.589 | -6.000 | 0.246 | 1.00 | . |
| 119 | C | CBA | . | HX2 | A | 3 | -12.183 | -6.559 | -1.109 | 1.00 | . |
| 120 | C | CGA | . | HX2 | A | 3 | -10.866 | -6.015 | -1.629 | 1.00 | . |
| 121 | O | O2A | . | HX2 | A | 3 | -9.808 | -6.469 | -1.146 | 1.00 | . |
| 122 | O | O1A | . | HX2 | A | 3 | -10.900 | -5.137 | -2.517 | 1.00 | . |
| 123 | C | C3A | . | HX2 | A | 3 | -14.556 | -4.752 | -0.691 | 1.00 | . |
| 124 | C | CMA | . | HX2 | A | 3 | -15.806 | -4.969 | -0.321 | 1.00 | . |
| 125 | C | C4D | . | HX2 | A | 3 | -12.269 | -1.125 | -0.529 | 1.00 | . |
| 126 | N | ND | . | HX2 | A | 3 | -12.847 | -0.787 | -1.720 | 1.00 | . |
| 127 | C | C1D | . | HX2 | A | 3 | -13.713 | 0.214 | -1.355 | 1.00 | . |
| 128 | C | CHD | . | HX2 | A | 3 | -14.975 | 0.459 | -1.812 | 1.00 | . |
| 129 | C | C2D | . | HX2 | A | 3 | -13.078 | 0.946 | -0.255 | 1.00 | . |
| 130 | C | CMD | . | HX2 | A | 3 | -13.488 | 2.282 | 0.293 | 1.00 | . |
| 131 | C | C3D | . | HX2 | A | 3 | -12.087 | 0.132 | 0.194 | 1.00 | . |
| 132 | C | CAD | . | HX2 | A | 3 | -11.075 | 0.477 | 1.241 | 1.00 | . |
| 133 | C | CBD | . | HX2 | A | 3 | -11.691 | 0.711 | 2.612 | 1.00 | . |
| 134 | C | CGD | . | HX2 | A | 3 | -13.048 | 1.380 | 2.535 | 1.00 | . |
| 135 | O | O1D | . | HX2 | A | 3 | -13.190 | 2.488 | 3.094 | 1.00 | . |

| | | | | | | | | | | | |
|---|---|---|---|---|---|---|---|---|---|---|---|
| 136 | O | O2D | . | HX2 | A | 3 | -13.958 | 0.791 | 1.914 | 1.00 | . |
| 137 | C | C2B | . | HX2 | A | 3 | -17.183 | -4.795 | -4.148 | 1.00 | . |
| 138 | C | CMB | . | HX2 | A | 3 | -17.003 | -5.140 | -5.598 | 1.00 | . |
| 139 | C | C3B | . | HX2 | A | 3 | -18.254 | -4.224 | -3.549 | 1.00 | . |
| 140 | C | CAB | . | HX2 | A | 3 | -19.554 | -3.794 | -4.168 | 1.00 | . |
| 141 | C | CBB | . | HX2 | A | 3 | -19.315 | -2.955 | -5.403 | 1.00 | . |
| 142 | S | S2 | . | HX2 | A | 3 | -20.641 | -2.944 | -2.986 | 1.00 | . |
| 143 | C | C2C | . | HX2 | A | 3 | -18.162 | -0.709 | -0.560 | 1.00 | . |
| 144 | C | CMC | . | HX2 | A | 3 | -19.488 | -0.615 | 0.136 | 1.00 | . |
| 145 | H | H1 | . | HX2 | A | 3 | -16.008 | 2.399 | -1.677 | 1.00 | . |
| 146 | H | H21 | . | HX2 | A | 3 | -17.491 | 2.772 | -2.022 | 1.00 | . |
| 147 | H | H31 | . | HX2 | A | 3 | -16.717 | 3.575 | -0.920 | 1.00 | . |
| 148 | H | H4 | . | HX2 | A | 3 | -18.289 | 1.931 | -0.070 | 1.00 | . |
| 149 | H | H6 | . | HX2 | A | 3 | -15.867 | -2.101 | -2.126 | 1.00 | . |
| 150 | H | H7 | . | HX2 | A | 3 | -18.857 | -3.373 | -0.605 | 1.00 | . |
| 151 | H | H8 | . | HX2 | A | 3 | -14.408 | -5.128 | -3.940 | 1.00 | . |
| 152 | H | H9 | . | HX2 | A | 3 | -12.252 | -3.979 | -2.617 | 1.00 | . |
| 153 | H | H10 | . | HX2 | A | 3 | -10.573 | -2.594 | 1.228 | 1.00 | . |
| 154 | H | H11 | . | HX2 | A | 3 | -11.917 | -3.544 | 2.904 | 1.00 | . |
| 155 | H | H13 | . | HX2 | A | 3 | -13.172 | -6.651 | 0.692 | 1.00 | . |
| 156 | H | H14 | . | HX2 | A | 3 | -11.784 | -5.906 | 0.800 | 1.00 | . |
| 157 | H | H15 | . | HX2 | A | 3 | -12.114 | -7.536 | -1.039 | 1.00 | . |
| 158 | H | H16 | . | HX2 | A | 3 | -12.887 | -6.352 | -1.759 | 1.00 | . |
| 159 | H | H17 | . | HX2 | A | 3 | -16.486 | -5.030 | -0.975 | 1.00 | . |
| 160 | H | H18 | . | HX2 | A | 3 | -16.018 | -5.061 | 0.594 | 1.00 | . |
| 161 | H | H19 | . | HX2 | A | 3 | -15.087 | 1.218 | -2.377 | 1.00 | . |
| 162 | H | H20 | . | HX2 | A | 3 | -12.705 | 2.842 | 0.406 | 1.00 | . |
| 163 | H | H211 | . | HX2 | A | 3 | -14.104 | 2.710 | -0.320 | 1.00 | . |
| 164 | H | H22 | . | HX2 | A | 3 | -13.922 | 2.161 | 1.152 | 1.00 | . |
| 165 | H | H23 | . | HX2 | A | 3 | -10.603 | 1.289 | 0.968 | 1.00 | . |
| 166 | H | H24 | . | HX2 | A | 3 | -10.417 | -0.242 | 1.305 | 1.00 | . |
| 167 | H | H25 | . | HX2 | A | 3 | -11.796 | -0.143 | 3.083 | 1.00 | . |
| 168 | H | H26 | . | HX2 | A | 3 | -11.100 | 1.273 | 3.155 | 1.00 | . |
| 169 | H | H27 | . | HX2 | A | 3 | -17.757 | -5.666 | -5.903 | 1.00 | . |
| 170 | H | H28 | . | HX2 | A | 3 | -16.187 | -5.651 | -5.711 | 1.00 | . |
| 171 | H | H29 | . | HX2 | A | 3 | -16.947 | -4.325 | -6.121 | 1.00 | . |
| 172 | H | H30 | . | HX2 | A | 3 | -20.025 | -4.618 | -4.450 | 1.00 | . |
| 173 | H | H311 | . | HX2 | A | 3 | -18.739 | -3.441 | -6.019 | 1.00 | . |
| 174 | H | H32 | . | HX2 | A | 3 | -18.887 | -2.118 | -5.150 | 1.00 | . |
| 175 | H | H33 | . | HX2 | A | 3 | -20.165 | -2.766 | -5.838 | 1.00 | . |
| 176 | H | H34 | . | HX2 | A | 3 | -20.741 | -1.875 | -3.550 | 1.00 | . |
| 177 | H | H35 | . | HX2 | A | 3 | -19.486 | 0.143 | 0.741 | 1.00 | . |
| 178 | H | H36 | . | HX2 | A | 3 | -20.193 | -0.500 | -0.522 | 1.00 | . |
| 179 | H | H37 | . | HX2 | A | 3 | -19.649 | -1.430 | 0.639 | 1.00 | . |
| 180 | N | N | . | HX2 | A | 3 | -12.938 | -5.131 | 4.811 | 1.00 | . |
| 181 | C | CA | . | HX2 | A | 3 | -14.316 | -4.576 | 4.781 | 1.00 | . |
| 182 | C | C | . | HX2 | A | 3 | -15.301 | -5.680 | 4.361 | 1.00 | . |
| 183 | O | O | . | HX2 | A | 3 | -14.956 | -6.863 | 4.572 | 1.00 | . |
| 184 | C | CB | . | HX2 | A | 3 | -14.390 | -3.396 | 3.815 | 1.00 | . |
| 185 | C | CG | . | HX2 | A | 3 | -14.628 | -3.818 | 2.372 | 1.00 | . |
| 186 | O | OXT | . | HX2 | A | 3 | -16.377 | -5.316 | 3.838 | 1.00 | . |
| 187 | H | H | . | HX2 | A | 3 | -12.693 | -5.391 | 3.982 | 1.00 | . |
| 188 | H | H2 | . | HX2 | A | 3 | -12.905 | -5.844 | 5.363 | 1.00 | . |
| 189 | H | H3 | . | HX2 | A | 3 | -12.359 | -4.506 | 5.104 | 1.00 | . |
| 190 | H | HA | . | HX2 | A | 3 | -14.550 | -4.268 | 5.694 | 1.00 | . |
| 191 | H | HB3 | . | HX2 | A | 3 | -13.550 | -2.893 | 3.865 | 1.00 | . |
| 192 | H | HB2 | . | HX2 | A | 3 | -15.117 | -2.801 | 4.097 | 1.00 | . |
| 193 | H | HG3 | . | HX2 | A | 3 | -15.206 | -3.148 | 1.936 | 1.00 | . |
| 194 | H | HG2 | . | HX2 | A | 3 | -15.111 | -4.679 | 2.369 | 1.00 | . |
| 195 | N | N | . | CYS | A | 5 | -19.811 | 3.740 | 2.578 | 1.00 | . |
| 196 | C | CA | . | CYS | A | 5 | -18.962 | 2.811 | 1.785 | 1.00 | . |

| | | | | | | | | | | | |
|---|---|---|---|---|---|---|---|---|---|---|---|
| 197 | C | C | . | CYS | A | 5 | -19.729 | 1.504 | 1.526 | 1.00 | . |
| 198 | O | O | . | CYS | A | 5 | -19.804 | 0.689 | 2.472 | 1.00 | . |
| 199 | C | CB | . | CYS | A | 5 | -17.613 | 2.541 | 2.453 | 1.00 | . |
| 200 | S | SG | . | CYS | A | 5 | -16.397 | 2.024 | 1.264 | 1.00 | . |
| 201 | O | OXT | . | CYS | A | 5 | -20.223 | 1.349 | 0.387 | 1.00 | . |
| 202 | H | H | . | CYS | A | 5 | -20.270 | 3.277 | 3.200 | 1.00 | . |
| 203 | H | H2 | . | CYS | A | 5 | -19.292 | 4.352 | 2.988 | 1.00 | . |
| 204 | H | H3 | . | CYS | A | 5 | -20.393 | 4.162 | 2.032 | 1.00 | . |
| 205 | H | HA | . | CYS | A | 5 | -18.768 | 3.241 | 0.914 | 1.00 | . |
| 206 | H | HB3 | . | CYS | A | 5 | -17.316 | 3.364 | 2.895 | 1.00 | . |
| 207 | H | HB2 | . | CYS | A | 5 | -17.737 | 1.840 | 3.127 | 1.00 | . |
| 208 | H | HG | . | CYS | A | 5 | -15.567 | 2.908 | 1.090 | 1.00 | . |
| 209 | H | HSG1 | . | CYS | A | 5 | -15.851 | 0.991 | 1.634 | 1.00 | . |
| 210 | N | N | . | CYS | A | 9 | 7.013 | 1.935 | -2.367 | 1.00 | . |
| 211 | C | CA | . | CYS | A | 9 | 7.311 | 1.474 | -0.985 | 1.00 | . |
| 212 | C | C | . | CYS | A | 9 | 6.245 | 2.016 | -0.016 | 1.00 | . |
| 213 | O | O | . | CYS | A | 9 | 5.639 | 1.183 | 0.694 | 1.00 | . |
| 214 | C | CB | . | CYS | A | 9 | 8.720 | 1.856 | -0.529 | 1.00 | . |
| 215 | S | SG | . | CYS | A | 9 | 9.210 | 0.966 | 0.930 | 1.00 | . |
| 216 | O | OXT | . | CYS | A | 9 | 6.060 | 3.254 | -0.009 | 1.00 | . |
| 217 | H | H | . | CYS | A | 9 | 7.009 | 2.837 | -2.394 | 1.00 | . |
| 218 | H | H2 | . | CYS | A | 9 | 6.206 | 1.627 | -2.625 | 1.00 | . |
| 219 | H | H3 | . | CYS | A | 9 | 7.642 | 1.628 | -2.935 | 1.00 | . |
| 220 | H | HA | . | CYS | A | 9 | 7.267 | 0.484 | -0.977 | 1.00 | . |
| 221 | H | HB3 | . | CYS | A | 9 | 8.733 | 2.818 | -0.343 | 1.00 | . |
| 222 | H | HB2 | . | CYS | A | 9 | 9.342 | 1.670 | -1.264 | 1.00 | . |
| 223 | H | HG | . | CYS | A | 9 | 8.599 | 1.428 | 1.887 | 1.00 | . |
| 224 | H | HSG1 | . | CYS | A | 9 | 8.951 | -0.223 | 0.844 | 1.00 | . |
| 225 | C | CBC | . | HX2 | A | 11 | 5.922 | -3.937 | -0.076 | 1.00 | . |
| 226 | C | CAC | . | HX2 | A | 11 | 6.886 | -4.073 | -1.237 | 1.00 | . |
| 227 | S | S1 | . | HX2 | A | 11 | 7.220 | -5.811 | -1.652 | 1.00 | . |
| 228 | C | C3C | . | HX2 | A | 11 | 8.168 | -3.316 | -1.017 | 1.00 | . |
| 229 | C | C4C | . | HX2 | A | 11 | 9.150 | -3.607 | -0.002 | 1.00 | . |
| 230 | N | NC | . | HX2 | A | 11 | 10.160 | -2.690 | -0.147 | 1.00 | . |
| 231 | C | C1C | . | HX2 | A | 11 | 9.890 | -1.824 | -1.176 | 1.00 | . |
| 232 | C | CHC | . | HX2 | A | 11 | 10.719 | -0.773 | -1.562 | 1.00 | . |
| 233 | C | C4B | . | HX2 | A | 11 | 11.795 | -0.249 | -0.886 | 1.00 | . |
| 234 | N | NB | . | HX2 | A | 11 | 13.063 | -0.174 | -1.378 | 1.00 | . |
| 235 | C | C1B | . | HX2 | A | 11 | 13.766 | -0.715 | -0.336 | 1.00 | . |
| 236 | C | CHB | . | HX2 | A | 11 | 14.889 | -1.512 | -0.482 | 1.00 | . |
| 237 | C | C4A | . | HX2 | A | 11 | 15.119 | -2.757 | 0.089 | 1.00 | . |
| 238 | N | NA | . | HX2 | A | 11 | 14.243 | -3.521 | 0.864 | 1.00 | . |
| 239 | C | C1A | . | HX2 | A | 11 | 14.401 | -4.822 | 0.508 | 1.00 | . |
| 240 | C | CHA | . | HX2 | A | 11 | 13.490 | -5.891 | 0.432 | 1.00 | . |
| 241 | N | NZ | . | HX2 | A | 11 | 13.832 | -6.860 | -0.454 | 1.00 | . |
| 242 | C | CE | . | HX2 | A | 11 | 14.773 | -6.666 | -1.423 | 1.00 | . |
| 243 | C | CD | . | HX2 | A | 11 | 15.829 | -5.780 | -1.231 | 1.00 | . |
| 244 | C | C2A | . | HX2 | A | 11 | 15.863 | -4.990 | 0.075 | 1.00 | . |
| 245 | C | CAA | . | HX2 | A | 11 | 16.709 | -5.774 | 1.113 | 1.00 | . |
| 246 | C | CBA | . | HX2 | A | 11 | 16.856 | -5.208 | 2.533 | 1.00 | . |
| 247 | C | CGA | . | HX2 | A | 11 | 18.221 | -4.622 | 2.855 | 1.00 | . |
| 248 | O | O2A | . | HX2 | A | 11 | 18.320 | -3.385 | 2.997 | 1.00 | . |
| 249 | O | O1A | . | HX2 | A | 11 | 19.186 | -5.409 | 2.962 | 1.00 | . |
| 250 | C | C3A | . | HX2 | A | 11 | 16.277 | -3.514 | -0.004 | 1.00 | . |
| 251 | C | CMA | . | HX2 | A | 11 | 17.513 | -3.054 | -0.099 | 1.00 | . |
| 252 | C | C4D | . | HX2 | A | 11 | 12.247 | -5.935 | 1.209 | 1.00 | . |
| 253 | N | ND | . | HX2 | A | 11 | 11.522 | -4.815 | 1.510 | 1.00 | . |
| 254 | C | C1D | . | HX2 | A | 11 | 10.229 | -5.266 | 1.468 | 1.00 | . |
| 255 | C | CHD | . | HX2 | A | 11 | 9.135 | -4.629 | 0.943 | 1.00 | . |
| 256 | C | C2D | . | HX2 | A | 11 | 10.251 | -6.626 | 2.009 | 1.00 | . |
| 257 | C | CMD | . | HX2 | A | 11 | 9.122 | -7.340 | 2.695 | 1.00 | . |

| | | | | | | | | | | | |
|---|---|---|---|---|---|---|---|---|---|---|---|
| 258 | C | C3D | . | HX2 | A | 11 | 11.510 | -7.080 | 1.755 | 1.00 | . |
| 259 | C | CAD | . | HX2 | A | 11 | 12.033 | -8.464 | 1.976 | 1.00 | . |
| 260 | C | CBD | . | HX2 | A | 11 | 11.396 | -9.500 | 1.066 | 1.00 | . |
| 261 | C | CGD | . | HX2 | A | 11 | 11.834 | -10.913 | 1.388 | 1.00 | . |
| 262 | O | O1D | . | HX2 | A | 11 | 11.312 | -11.478 | 2.372 | 1.00 | . |
| 263 | O | O2D | . | HX2 | A | 11 | 12.696 | -11.445 | 0.655 | 1.00 | . |
| 264 | C | C2B | . | HX2 | A | 11 | 13.095 | -0.320 | 0.897 | 1.00 | . |
| 265 | C | CMB | . | HX2 | A | 11 | 13.635 | -0.513 | 2.286 | 1.00 | . |
| 266 | C | C3B | . | HX2 | A | 11 | 11.920 | 0.235 | 0.478 | 1.00 | . |
| 267 | C | CAB | . | HX2 | A | 11 | 11.000 | 1.250 | 1.119 | 1.00 | . |
| 268 | C | CBB | . | HX2 | A | 11 | 11.198 | 1.387 | 2.631 | 1.00 | . |
| 269 | C | C2C | . | HX2 | A | 11 | 8.623 | -2.218 | -1.730 | 1.00 | . |
| 270 | C | CMC | . | HX2 | A | 11 | 7.933 | -1.548 | -2.885 | 1.00 | . |
| 271 | H | H1 | . | HX2 | A | 11 | 6.250 | -3.286 | 0.549 | 1.00 | . |
| 272 | H | H21 | . | HX2 | A | 11 | 5.832 | -4.785 | 0.367 | 1.00 | . |
| 273 | H | H31 | . | HX2 | A | 11 | 5.064 | -3.658 | -0.405 | 1.00 | . |
| 274 | H | H4 | . | HX2 | A | 11 | 6.440 | -3.693 | -2.032 | 1.00 | . |
| 275 | H | H5 | . | HX2 | A | 11 | 8.299 | -5.678 | -2.188 | 1.00 | . |
| 276 | H | H6 | . | HX2 | A | 11 | 10.879 | -2.662 | 0.358 | 1.00 | . |
| 277 | H | H7 | . | HX2 | A | 11 | 10.536 | -0.391 | -2.416 | 1.00 | . |
| 278 | H | H8 | . | HX2 | A | 11 | 15.568 | -1.169 | -1.060 | 1.00 | . |
| 279 | H | H9 | . | HX2 | A | 11 | 13.682 | -3.232 | 1.473 | 1.00 | . |
| 280 | H | H10 | . | HX2 | A | 11 | 13.426 | -7.650 | -0.403 | 1.00 | . |
| 281 | H | H11 | . | HX2 | A | 11 | 14.697 | -7.128 | -2.239 | 1.00 | . |
| 282 | H | H13 | . | HX2 | A | 11 | 17.608 | -5.890 | 0.738 | 1.00 | . |
| 283 | H | H14 | . | HX2 | A | 11 | 16.325 | -6.672 | 1.191 | 1.00 | . |
| 284 | H | H15 | . | HX2 | A | 11 | 16.673 | -5.929 | 3.173 | 1.00 | . |
| 285 | H | H16 | . | HX2 | A | 11 | 16.179 | -4.514 | 2.678 | 1.00 | . |
| 286 | H | H17 | . | HX2 | A | 11 | 18.242 | -3.657 | -0.111 | 1.00 | . |
| 287 | H | H18 | . | HX2 | A | 11 | 17.668 | -2.123 | -0.153 | 1.00 | . |
| 288 | H | H19 | . | HX2 | A | 11 | 8.281 | -4.933 | 1.235 | 1.00 | . |
| 289 | H | H20 | . | HX2 | A | 11 | 9.453 | -8.147 | 3.116 | 1.00 | . |
| 290 | H | H211 | . | HX2 | A | 11 | 8.442 | -7.573 | 2.045 | 1.00 | . |
| 291 | H | H22 | . | HX2 | A | 11 | 8.735 | -6.762 | 3.370 | 1.00 | . |
| 292 | H | H23 | . | HX2 | A | 11 | 11.873 | -8.720 | 2.906 | 1.00 | . |
| 293 | H | H24 | . | HX2 | A | 11 | 13.001 | -8.466 | 1.829 | 1.00 | . |
| 294 | H | H25 | . | HX2 | A | 11 | 10.419 | -9.455 | 1.140 | 1.00 | . |
| 295 | H | H26 | . | HX2 | A | 11 | 11.625 | -9.309 | 0.131 | 1.00 | . |
| 296 | H | H27 | . | HX2 | A | 11 | 13.904 | 0.341 | 2.654 | 1.00 | . |
| 297 | H | H28 | . | HX2 | A | 11 | 14.405 | -1.099 | 2.264 | 1.00 | . |
| 298 | H | H29 | . | HX2 | A | 11 | 12.954 | -0.907 | 2.850 | 1.00 | . |
| 299 | H | H30 | . | HX2 | A | 11 | 11.197 | 2.135 | 0.714 | 1.00 | . |
| 300 | H | H311 | . | HX2 | A | 11 | 10.530 | 1.995 | 2.993 | 1.00 | . |
| 301 | H | H32 | . | HX2 | A | 11 | 11.104 | 0.517 | 3.053 | 1.00 | . |
| 302 | H | H33 | . | HX2 | A | 11 | 12.084 | 1.746 | 2.812 | 1.00 | . |
| 303 | H | H35 | . | HX2 | A | 11 | 6.980 | -1.490 | -2.704 | 1.00 | . |
| 304 | H | H36 | . | HX2 | A | 11 | 8.076 | -2.064 | -3.694 | 1.00 | . |
| 305 | H | H37 | . | HX2 | A | 11 | 8.288 | -0.654 | -3.007 | 1.00 | . |
| 306 | N | N | . | HX2 | A | 11 | 19.366 | -8.396 | -2.799 | 1.00 | . |
| 307 | C | CA | . | HX2 | A | 11 | 18.228 | -7.785 | -2.058 | 1.00 | . |
| 308 | C | C | . | HX2 | A | 11 | 17.347 | -8.905 | -1.473 | 1.00 | . |
| 309 | O | O | . | HX2 | A | 11 | 16.954 | -9.795 | -2.260 | 1.00 | . |
| 310 | C | CB | . | HX2 | A | 11 | 17.437 | -6.852 | -2.979 | 1.00 | . |
| 311 | C | CG | . | HX2 | A | 11 | 16.898 | -5.597 | -2.290 | 1.00 | . |
| 312 | O | OXT | . | HX2 | A | 11 | 17.085 | -8.847 | -0.251 | 1.00 | . |
| 313 | H | H | . | HX2 | A | 11 | 19.061 | -8.821 | -3.533 | 1.00 | . |
| 314 | H | H2 | . | HX2 | A | 11 | 19.801 | -8.988 | -2.276 | 1.00 | . |
| 315 | H | H3 | . | HX2 | A | 11 | 19.949 | -7.756 | -3.052 | 1.00 | . |
| 316 | H | HA | . | HX2 | A | 11 | 18.600 | -7.255 | -1.308 | 1.00 | . |
| 317 | H | HB3 | . | HX2 | A | 11 | 18.020 | -6.573 | -3.716 | 1.00 | . |
| 318 | H | HB2 | . | HX2 | A | 11 | 16.691 | -7.348 | -3.377 | 1.00 | . |

```
319 H  HG3  . HX2 A   11    16.545    -4.990    -2.984  1.00 .
320 H  HG2  . HX2 A   11    17.666    -5.127    -1.888  1.00 .

data_comp_list

loop_
_chem_comp.id
_chem_comp.three_letter_code
_chem_comp.name
_chem_comp.group
_chem_comp.number_atoms_all
_chem_comp.number_atoms_nh
_chem_comp.desc_level
HX2      HX2 .                                      peptide            98
54 .

data_mod_list

loop_
_chem_mod.id
_chem_mod.name
_chem_mod.comp_id
_chem_mod.group_id
HX2mod1  .                                              HX2      .
CYSmod1  .                                              CYS      .
CYSmod2  .                                              CYS      .
HX2mod2  .                                              HX2      .

data_link_list

loop_
_chem_link.id
_chem_link.comp_id_1
_chem_link.mod_id_1
_chem_link.group_comp_1
_chem_link.comp_id_2
_chem_link.mod_id_2
_chem_link.group_comp_2
_chem_link.name
HX2-CYS  HX2      HX2mod1  .         CYS      CYSmod1  .
.
CYS-HX2  CYS      CYSmod2  .         HX2      HX2mod2  .
.

data_comp_HX2

loop_
_chem_comp_atom.comp_id
_chem_comp_atom.atom_id
_chem_comp_atom.type_symbol
_chem_comp_atom.type_energy
_chem_comp_atom.charge
_chem_comp_atom.x
_chem_comp_atom.y
_chem_comp_atom.z
HX2           CBC    C    CH3      0     -6.641   -7.573    0.719
HX2           CAC    C    CH1      0     -7.081   -8.936    0.230
HX2           S1     S    SH1      0     -8.068   -9.810    1.479
HX2           C3C    C    CR5      0     -5.909   -9.768   -0.200
```

| | | | | | | | |
|---|---|---|---|---|---|---|---|
| HX2 | C4C | C | CR5 | 0 | -5.630 | -11.153 | 0.091 |
| HX2 | NC | N | NR15 | 0 | -4.437 | -11.454 | -0.515 |
| HX2 | C1C | C | CR5 | 0 | -3.950 | -10.365 | -1.191 |
| HX2 | CHC | C | C1 | 0 | -2.748 | -10.327 | -1.875 |
| HX2 | C4B | C | CR5 | 0 | -1.533 | -10.269 | -1.260 |
| HX2 | NB | N | NRD5 | 0 | -0.458 | -11.076 | -1.501 |
| HX2 | C1B | C | CR5 | 0 | 0.117 | -11.139 | -0.261 |
| HX2 | CHB | C | C1 | 0 | 0.649 | -12.275 | 0.323 |
| HX2 | C4A | C | CR5 | 0 | -0.030 | -13.482 | 0.375 |
| HX2 | NA | N | NR15 | 0 | -1.298 | -13.719 | 0.914 |
| HX2 | C1A | C | CR56 | 0 | -1.933 | -14.576 | 0.071 |
| HX2 | CHA | C | CR6 | 0 | -3.236 | -14.657 | -0.431 |
| HX2 | NZ | N | NR16 | 0 | -3.262 | -14.679 | -1.782 |
| HX2 | CE | C | CR16 | 0 | -2.139 | -14.930 | -2.493 |
| HX2 | CD | C | CR6 | 0 | -1.145 | -15.674 | -1.924 |
| HX2 | C2A | C | CT | 0 | -0.855 | -15.513 | -0.446 |
| HX2 | CAA | C | CH2 | 0 | -0.808 | -16.870 | 0.292 |
| HX2 | CBA | C | CH2 | 0 | 0.572 | -17.295 | 0.774 |
| HX2 | CGA | C | C | 0 | 1.609 | -17.386 | -0.331 |
| HX2 | O2A | O | O | 0 | 2.377 | -18.370 | -0.338 |
| HX2 | O1A | O | OC | -1 | 1.648 | -16.472 | -1.181 |
| HX2 | C3A | C | CR5 | 0 | 0.398 | -14.697 | -0.132 |
| HX2 | CMA | C | C2 | 0 | 1.641 | -15.100 | -0.319 |
| HX2 | C4D | C | CR5 | 0 | -4.485 | -14.675 | 0.334 |
| HX2 | ND | N | NRD5 | 0 | -5.643 | -14.166 | -0.175 |
| HX2 | C1D | C | CR5 | 0 | -6.043 | -13.391 | 0.876 |
| HX2 | CHD | C | C1 | 0 | -6.366 | -12.067 | 0.831 |
| HX2 | C2D | C | CR5 | 0 | -5.823 | -14.207 | 2.072 |
| HX2 | CMD | C | CH3 | 0 | -6.525 | -14.092 | 3.394 |
| HX2 | C3D | C | CR5 | 0 | -4.846 | -15.078 | 1.697 |
| HX2 | CAD | C | CH2 | 0 | -4.292 | -16.224 | 2.484 |
| HX2 | CBD | C | CH2 | 0 | -4.746 | -17.656 | 2.253 |
| HX2 | CGD | C | C | 0 | -3.670 | -18.676 | 2.566 |
| HX2 | O1D | O | O | 0 | -3.193 | -18.691 | 3.720 |
| HX2 | O2D | O | OC | -1 | -3.314 | -19.452 | 1.654 |
| HX2 | C2B | C | CR5 | 0 | -0.044 | -9.818 | 0.324 |
| HX2 | CMB | C | CH3 | 0 | 0.900 | -9.109 | 1.250 |
| HX2 | C3B | C | CR5 | 0 | -1.225 | -9.379 | -0.160 |
| HX2 | CAB | C | CH1 | 0 | -2.097 | -8.254 | 0.325 |
| HX2 | CBB | C | CH3 | 0 | -1.883 | -7.984 | 1.797 |
| HX2 | S2 | S | SH1 | 0 | -3.854 | -8.546 | -0.033 |
| HX2 | C2C | C | CR5 | 0 | -4.874 | -9.290 | -0.977 |
| HX2 | CMC | C | CH3 | 0 | -4.720 | -7.900 | -1.520 |
| HX2 | H1 | H | H | 0 | -5.862 | -7.294 | 0.231 |
| HX2 | H21 | H | H | 0 | -7.349 | -6.940 | 0.584 |
| HX2 | H31 | H | H | 0 | -6.430 | -7.623 | 1.655 |
| HX2 | H4 | H | H | 0 | -7.662 | -8.800 | -0.561 |
| HX2 | H5 | H | HSH1 | 0 | -7.279 | -9.790 | 2.399 |
| HX2 | H6 | H | H | 0 | -4.048 | -12.241 | -0.490 |
| HX2 | H7 | H | H | 0 | -2.764 | -10.326 | -2.827 |
| HX2 | H8 | H | H | 0 | 1.524 | -12.219 | 0.701 |
| HX2 | H9 | H | H | 0 | -1.633 | -13.385 | 1.654 |
| HX2 | H10 | H | H | 0 | -4.029 | -14.526 | -2.207 |
| HX2 | H11 | H | H | 0 | -2.033 | -14.601 | -3.369 |
| HX2 | H13 | H | H | 0 | -1.158 | -17.563 | -0.307 |
| HX2 | H14 | H | H | 0 | -1.409 | -16.826 | 1.065 |
| HX2 | H15 | H | H | 0 | 0.883 | -16.651 | 1.445 |
| HX2 | H16 | H | H | 0 | 0.497 | -18.172 | 1.207 |
| HX2 | H17 | H | H | 0 | 2.362 | -14.530 | -0.095 |
| HX2 | H18 | H | H | 0 | 1.810 | -15.958 | -0.676 |
| HX2 | H19 | H | H | 0 | -7.112 | -11.768 | 1.345 |

```
HX2          H20    H    H        0     -7.472  -13.941    3.247
HX2          H211   H    H        0     -6.405  -14.911    3.899
HX2          H22    H    H        0     -6.156  -13.350    3.896
HX2          H23    H    H        0     -3.999  -16.415    1.571
HX2          H24    H    H        0     -5.235  -16.211    2.740
HX2          H25    H    H        0     -5.529  -17.853    2.811
HX2          H26    H    H        0     -5.020  -17.773    1.319
HX2          H27    H    H        0      0.420   -8.425    1.741
HX2          H28    H    H        0      1.284   -9.744    1.873
HX2          H29    H    H        0      1.610   -8.695    0.735
HX2          H30    H    H        0     -1.829   -7.443   -0.175
HX2          H311   H    H        0     -1.654   -7.170    2.278
HX2          H32    H    H        0     -2.699   -7.474    1.655
HX2          H33    H    H        0     -0.975   -7.853    1.474
HX2          H34    H    HSH1     0     -4.093   -9.342    0.852
HX2          H35    H    H        0     -5.512   -7.379   -1.313
HX2          H36    H    H        0     -3.943   -7.478   -1.118
HX2          H37    H    H        0     -4.602   -7.938   -2.483
HX2          N      N    NT3      1     -2.370  -19.884   -2.666
HX2          CA     C    CH1      0     -1.083  -19.140   -2.702
HX2          C      C    C        0     -0.528  -19.017   -1.274
HX2          O      O    O        0      0.650  -19.394   -1.084
HX2          CB     C    CH2      0     -1.283  -17.760   -3.329
HX2          CG     C    CH2      0     -0.379  -16.685   -2.741
HX2          OXT    O    OC      -1     -1.293  -18.549   -0.402
HX2          H      H    H        0     -2.463  -20.297   -1.869
HX2          H2     H    H        0     -2.386  -20.503   -3.321
HX2          H3     H    H        0     -3.060  -19.313   -2.783
HX2          HA     H    H        0     -0.440  -19.656   -3.251
HX2          HB3    H    H        0     -2.217  -17.491   -3.201
HX2          HB2    H    H        0     -1.116  -17.825   -4.292
HX2          HG3    H    H        0      0.300  -17.114   -2.168
HX2          HG2    H    H        0      0.085  -16.218   -3.477

loop_
_chem_comp_bond.comp_id
_chem_comp_bond.atom_id_1
_chem_comp_bond.atom_id_2
_chem_comp_bond.type
_chem_comp_bond.aromatic
_chem_comp_bond.value_dist
_chem_comp_bond.value_dist_esd
HX2      N      CA        single   n    1.487    0.010
HX2      CA     C         single   n    1.538    0.011
HX2      C      O         double   n    1.251    0.018
HX2      C      OXT       single   n    1.251    0.018
HX2      CA     CB        single   n    1.529    0.010
HX2      CB     CG        single   n    1.523    0.019
HX2      CD     CG        single   n    1.511    0.010
HX2      CBC    CAC       single   n    1.513    0.012
HX2      CAC    C3C       single   n    1.503    0.010
HX2      CAC    S1        single   n    1.817    0.016
HX2      C3C    C4C       double   y    1.442    0.017
HX2      C3C    C2C       single   y    1.375    0.020
HX2      C4C    NC        single   y    1.373    0.010
HX2      NC     C1C       single   y    1.373    0.010
HX2      C1C    CHC       single   n    1.392    0.019
HX2      CHC    C4B       double   n    1.372    0.020
HX2      C4B    C3B       single   n    1.451    0.017
HX2      C4B    NB        single   n    1.365    0.010
HX2      NB     C1B       double   n    1.367    0.012
```

| | | | | | | |
|---|---|---|---|---|---|---|
| HX2 | C1B | C2B | single | n | 1.453 | 0.010 |
| HX2 | C1B | CHB | single | n | 1.384 | 0.011 |
| HX2 | CHB | C4A | double | n | 1.382 | 0.020 |
| HX2 | C4A | NA | single | n | 1.396 | 0.010 |
| HX2 | NA | C1A | single | n | 1.363 | 0.020 |
| HX2 | C1A | CHA | double | n | 1.396 | 0.020 |
| HX2 | CHA | NZ | single | n | 1.351 | 0.013 |
| HX2 | NZ | CE | single | n | 1.354 | 0.016 |
| HX2 | CE | CD | double | n | 1.368 | 0.020 |
| HX2 | C1A | C2A | single | n | 1.525 | 0.017 |
| HX2 | CD | C2A | single | n | 1.515 | 0.013 |
| HX2 | C2A | C3A | single | n | 1.525 | 0.017 |
| HX2 | C2A | CAA | single | n | 1.547 | 0.014 |
| HX2 | CAA | CBA | single | n | 1.521 | 0.019 |
| HX2 | CBA | CGA | single | n | 1.518 | 0.014 |
| HX2 | CGA | O2A | double | n | 1.249 | 0.016 |
| HX2 | CGA | O1A | single | n | 1.249 | 0.016 |
| HX2 | C4A | C3A | single | n | 1.381 | 0.020 |
| HX2 | C3A | CMA | double | n | 1.321 | 0.010 |
| HX2 | CHA | C4D | single | n | 1.464 | 0.015 |
| HX2 | C4D | ND | double | n | 1.366 | 0.020 |
| HX2 | ND | C1D | single | n | 1.374 | 0.016 |
| HX2 | C4C | CHD | single | n | 1.392 | 0.019 |
| HX2 | C1D | CHD | double | n | 1.372 | 0.020 |
| HX2 | C1D | C2D | single | n | 1.467 | 0.010 |
| HX2 | C2D | CMD | single | n | 1.501 | 0.010 |
| HX2 | C4D | C3D | single | n | 1.458 | 0.018 |
| HX2 | C2D | C3D | double | n | 1.360 | 0.011 |
| HX2 | C3D | CAD | single | n | 1.497 | 0.010 |
| HX2 | CAD | CBD | single | n | 1.519 | 0.020 |
| HX2 | CBD | CGD | single | n | 1.515 | 0.012 |
| HX2 | CGD | O2D | single | n | 1.249 | 0.016 |
| HX2 | CGD | O1D | double | n | 1.249 | 0.016 |
| HX2 | C2B | CMB | single | n | 1.501 | 0.010 |
| HX2 | C2B | C3B | double | n | 1.354 | 0.013 |
| HX2 | C3B | CAB | single | n | 1.505 | 0.014 |
| HX2 | CAB | CBB | single | n | 1.513 | 0.012 |
| HX2 | CAB | S2 | single | n | 1.817 | 0.016 |
| HX2 | C1C | C2C | double | y | 1.434 | 0.010 |
| HX2 | C2C | CMC | single | n | 1.501 | 0.011 |
| HX2 | CBC | H1 | single | n | 0.960 | 0.010 |
| HX2 | CBC | H21 | single | n | 0.960 | 0.010 |
| HX2 | CBC | H31 | single | n | 0.960 | 0.010 |
| HX2 | CAC | H4 | single | n | 0.990 | 0.020 |
| HX2 | S1 | H5 | single | n | 1.212 | 0.020 |
| HX2 | NC | H6 | single | n | 0.879 | 0.020 |
| HX2 | CHC | H7 | single | n | 0.953 | 0.019 |
| HX2 | CHB | H8 | single | n | 0.956 | 0.020 |
| HX2 | NA | H9 | single | n | 0.878 | 0.020 |
| HX2 | NZ | H10 | single | n | 0.890 | 0.020 |
| HX2 | CE | H11 | single | n | 0.942 | 0.018 |
| HX2 | CAA | H13 | single | n | 0.981 | 0.016 |
| HX2 | CAA | H14 | single | n | 0.981 | 0.016 |
| HX2 | CBA | H15 | single | n | 0.981 | 0.017 |
| HX2 | CBA | H16 | single | n | 0.981 | 0.017 |
| HX2 | CMA | H18 | single | n | 0.945 | 0.020 |
| HX2 | CMA | H17 | single | n | 0.945 | 0.020 |
| HX2 | CHD | H19 | single | n | 0.953 | 0.019 |
| HX2 | CMD | H20 | single | n | 0.969 | 0.015 |
| HX2 | CMD | H211 | single | n | 0.969 | 0.015 |
| HX2 | CMD | H22 | single | n | 0.969 | 0.015 |

```
HX2      CAD    H23       single   n    0.978    0.017
HX2      CAD    H24       single   n    0.978    0.017
HX2      CBD    H25       single   n    0.981    0.011
HX2      CBD    H26       single   n    0.981    0.011
HX2      CMB    H27       single   n    0.969    0.015
HX2      CMB    H28       single   n    0.969    0.015
HX2      CMB    H29       single   n    0.969    0.015
HX2      CAB    H30       single   n    0.990    0.020
HX2      CBB    H311      single   n    0.973    0.010
HX2      CBB    H33       single   n    0.973    0.010
HX2      CBB    H32       single   n    0.973    0.010
HX2      S2     H34       single   n    1.212    0.020
HX2      CMC    H35       single   n    0.971    0.014
HX2      CMC    H36       single   n    0.971    0.014
HX2      CMC    H37       single   n    0.971    0.014
HX2      N      H         single   n    0.902    0.010
HX2      N      H2        single   n    0.902    0.010
HX2      N      H3        single   n    0.902    0.010
HX2      CA     HA        single   n    0.991    0.020
HX2      CB     HB2       single   n    0.980    0.017
HX2      CB     HB3       single   n    0.980    0.017
HX2      CG     HG2       single   n    0.987    0.010
HX2      CG     HG3       single   n    0.987    0.010

loop_
_chem_comp_angle.comp_id
_chem_comp_angle.atom_id_1
_chem_comp_angle.atom_id_2
_chem_comp_angle.atom_id_3
_chem_comp_angle.value_angle
_chem_comp_angle.value_angle_esd
HX2      CAC    CBC    H1      109.518    1.500
HX2      CAC    CBC    H21     109.518    1.500
HX2      CAC    CBC    H31     109.518    1.500
HX2      H1     CBC    H21     109.460    1.500
HX2      H1     CBC    H31     109.460    1.500
HX2      H21    CBC    H31     109.460    1.500
HX2      CBC    CAC    C3C     112.400    1.500
HX2      CBC    CAC    S1      112.610    3.000
HX2      CBC    CAC    H4      108.549    2.040
HX2      S1     CAC    C3C     111.652    3.000
HX2      C3C    CAC    H4      108.177    1.500
HX2      S1     CAC    H4      108.757    3.000
HX2      CAC    S1     H5       99.186    3.000
HX2      CAC    C3C    C4C     126.768    3.000
HX2      CAC    C3C    C2C     125.511    3.000
HX2      C4C    C3C    C2C     107.721    1.500
HX2      C3C    C4C    NC      107.299    1.500
HX2      C3C    C4C    CHD     127.218    1.940
HX2      NC     C4C    CHD     125.483    1.570
HX2      C4C    NC     C1C     110.518    1.500
HX2      C4C    NC     H6      124.741    3.000
HX2      C1C    NC     H6      124.741    3.000
HX2      NC     C1C    CHC     125.600    1.570
HX2      NC     C1C    C2C     106.742    1.500
HX2      CHC    C1C    C2C     127.658    3.000
HX2      C1C    CHC    C4B     128.527    3.000
HX2      C1C    CHC    H7      115.410    2.170
HX2      C4B    CHC    H7      116.063    1.500
HX2      CHC    C4B    C3B     125.026    2.880
HX2      CHC    C4B    NB      125.218    1.500
```

| | | | | | |
|---|---|---|---|---|---|
| HX2 | NB | C4B | C3B | 109.756 | 2.280 |
| HX2 | C4B | NB | C1B | 106.462 | 2.670 |
| HX2 | NB | C1B | C2B | 110.179 | 3.000 |
| HX2 | NB | C1B | CHB | 125.177 | 1.680 |
| HX2 | CHB | C1B | C2B | 124.644 | 2.930 |
| HX2 | C1B | CHB | C4A | 125.148 | 3.000 |
| HX2 | C1B | CHB | H8 | 116.958 | 3.000 |
| HX2 | C4A | CHB | H8 | 117.894 | 3.000 |
| HX2 | CHB | C4A | NA | 126.984 | 3.000 |
| HX2 | CHB | C4A | C3A | 126.493 | 3.000 |
| HX2 | NA | C4A | C3A | 106.523 | 3.000 |
| HX2 | C4A | NA | C1A | 109.510 | 3.000 |
| HX2 | C4A | NA | H9 | 126.008 | 3.000 |
| HX2 | C1A | NA | H9 | 124.482 | 3.000 |
| HX2 | NA | C1A | CHA | 130.085 | 3.000 |
| HX2 | NA | C1A | C2A | 107.838 | 3.000 |
| HX2 | CHA | C1A | C2A | 122.076 | 3.000 |
| HX2 | C1A | CHA | NZ | 116.163 | 3.000 |
| HX2 | C1A | CHA | C4D | 122.143 | 3.000 |
| HX2 | NZ | CHA | C4D | 121.694 | 3.000 |
| HX2 | CHA | NZ | CE | 122.624 | 2.880 |
| HX2 | CHA | NZ | H10 | 118.481 | 3.000 |
| HX2 | CE | NZ | H10 | 118.895 | 3.000 |
| HX2 | NZ | CE | CD | 120.688 | 3.000 |
| HX2 | NZ | CE | H11 | 119.999 | 2.540 |
| HX2 | CD | CE | H11 | 119.312 | 1.500 |
| HX2 | CE | CD | CG | 120.858 | 3.000 |
| HX2 | C2A | CD | CG | 117.577 | 3.000 |
| HX2 | CE | CD | C2A | 121.565 | 3.000 |
| HX2 | C1A | C2A | CD | 109.895 | 3.000 |
| HX2 | C1A | C2A | C3A | 101.524 | 2.900 |
| HX2 | C1A | C2A | CAA | 111.194 | 3.000 |
| HX2 | CD | C2A | C3A | 109.856 | 3.000 |
| HX2 | CD | C2A | CAA | 109.486 | 3.000 |
| HX2 | CAA | C2A | C3A | 108.076 | 3.000 |
| HX2 | C2A | CAA | CBA | 114.329 | 3.000 |
| HX2 | C2A | CAA | H13 | 108.328 | 1.500 |
| HX2 | C2A | CAA | H14 | 108.328 | 1.500 |
| HX2 | CBA | CAA | H13 | 108.359 | 1.500 |
| HX2 | CBA | CAA | H14 | 108.359 | 1.500 |
| HX2 | H13 | CAA | H14 | 106.929 | 1.500 |
| HX2 | CAA | CBA | CGA | 113.560 | 3.000 |
| HX2 | CAA | CBA | H15 | 108.638 | 1.500 |
| HX2 | CAA | CBA | H16 | 108.638 | 1.500 |
| HX2 | CGA | CBA | H15 | 108.531 | 1.500 |
| HX2 | CGA | CBA | H16 | 108.531 | 1.500 |
| HX2 | H15 | CBA | H16 | 107.705 | 2.230 |
| HX2 | CBA | CGA | O2A | 118.194 | 3.000 |
| HX2 | CBA | CGA | O1A | 118.194 | 3.000 |
| HX2 | O2A | CGA | O1A | 123.612 | 1.820 |
| HX2 | C4A | C3A | C2A | 108.027 | 3.000 |
| HX2 | C2A | C3A | CMA | 124.704 | 3.000 |
| HX2 | C4A | C3A | CMA | 127.269 | 3.000 |
| HX2 | C3A | CMA | H18 | 119.932 | 1.500 |
| HX2 | C3A | CMA | H17 | 119.932 | 1.500 |
| HX2 | H17 | CMA | H18 | 120.136 | 1.500 |
| HX2 | CHA | C4D | ND | 123.012 | 3.000 |
| HX2 | CHA | C4D | C3D | 127.850 | 3.000 |
| HX2 | ND | C4D | C3D | 109.138 | 3.000 |
| HX2 | C4D | ND | C1D | 107.166 | 2.340 |
| HX2 | ND | C1D | CHD | 125.680 | 3.000 |

| | | | | | |
|---|---|---|---|---|---|
| HX2 | ND | C1D | C2D | 109.717 | 2.280 |
| HX2 | CHD | C1D | C2D | 124.603 | 1.500 |
| HX2 | C4C | CHD | C1D | 128.527 | 3.000 |
| HX2 | C4C | CHD | H19 | 115.410 | 2.170 |
| HX2 | C1D | CHD | H19 | 116.063 | 1.500 |
| HX2 | C1D | C2D | CMD | 126.711 | 1.500 |
| HX2 | C1D | C2D | C3D | 106.953 | 1.500 |
| HX2 | CMD | C2D | C3D | 126.336 | 3.000 |
| HX2 | C2D | CMD | H20 | 109.573 | 1.500 |
| HX2 | C2D | CMD | H211 | 109.573 | 1.500 |
| HX2 | C2D | CMD | H22 | 109.573 | 1.500 |
| HX2 | H20 | CMD | H211 | 109.306 | 2.100 |
| HX2 | H20 | CMD | H22 | 109.306 | 2.100 |
| HX2 | H211 | CMD | H22 | 109.306 | 2.100 |
| HX2 | C4D | C3D | C2D | 107.026 | 1.500 |
| HX2 | C4D | C3D | CAD | 126.632 | 3.000 |
| HX2 | C2D | C3D | CAD | 126.342 | 3.000 |
| HX2 | C3D | CAD | CBD | 113.552 | 1.680 |
| HX2 | C3D | CAD | H23 | 109.334 | 3.000 |
| HX2 | C3D | CAD | H24 | 109.334 | 3.000 |
| HX2 | CBD | CAD | H23 | 109.251 | 3.000 |
| HX2 | CBD | CAD | H24 | 109.251 | 3.000 |
| HX2 | H23 | CAD | H24 | 107.902 | 2.140 |
| HX2 | CAD | CBD | CGD | 113.745 | 3.000 |
| HX2 | CAD | CBD | H25 | 111.034 | 3.000 |
| HX2 | CAD | CBD | H26 | 111.034 | 3.000 |
| HX2 | CGD | CBD | H25 | 108.600 | 1.500 |
| HX2 | CGD | CBD | H26 | 108.600 | 1.500 |
| HX2 | H25 | CBD | H26 | 107.539 | 1.500 |
| HX2 | CBD | CGD | O2D | 118.035 | 1.950 |
| HX2 | CBD | CGD | O1D | 118.035 | 1.950 |
| HX2 | O1D | CGD | O2D | 123.930 | 1.820 |
| HX2 | C1B | C2B | CMB | 126.661 | 1.500 |
| HX2 | C1B | C2B | C3B | 106.802 | 1.500 |
| HX2 | CMB | C2B | C3B | 126.537 | 3.000 |
| HX2 | C2B | CMB | H27 | 109.573 | 1.500 |
| HX2 | C2B | CMB | H28 | 109.573 | 1.500 |
| HX2 | C2B | CMB | H29 | 109.573 | 1.500 |
| HX2 | H27 | CMB | H28 | 109.306 | 2.100 |
| HX2 | H27 | CMB | H29 | 109.306 | 2.100 |
| HX2 | H28 | CMB | H29 | 109.306 | 2.100 |
| HX2 | C4B | C3B | C2B | 106.802 | 1.500 |
| HX2 | C4B | C3B | CAB | 126.599 | 3.000 |
| HX2 | C2B | C3B | CAB | 126.599 | 3.000 |
| HX2 | C3B | CAB | CBB | 111.467 | 2.280 |
| HX2 | C3B | CAB | S2 | 112.915 | 1.840 |
| HX2 | C3B | CAB | H30 | 108.198 | 2.620 |
| HX2 | CBB | CAB | S2 | 112.610 | 3.000 |
| HX2 | CBB | CAB | H30 | 108.549 | 2.040 |
| HX2 | S2 | CAB | H30 | 108.757 | 3.000 |
| HX2 | CAB | CBB | H311 | 109.518 | 1.500 |
| HX2 | CAB | CBB | H33 | 109.518 | 1.500 |
| HX2 | CAB | CBB | H32 | 109.518 | 1.500 |
| HX2 | H311 | CBB | H33 | 109.466 | 1.500 |
| HX2 | H311 | CBB | H32 | 109.466 | 1.500 |
| HX2 | H32 | CBB | H33 | 109.466 | 1.500 |
| HX2 | CAB | S2 | H34 | 99.186 | 3.000 |
| HX2 | C3C | C2C | C1C | 107.721 | 1.500 |
| HX2 | C3C | C2C | CMC | 126.856 | 3.000 |
| HX2 | C1C | C2C | CMC | 125.423 | 1.500 |
| HX2 | C2C | CMC | H35 | 109.572 | 1.500 |

```
HX2       C2C    CMC    H36     109.572    1.500
HX2       C2C    CMC    H37     109.572    1.500
HX2       H35    CMC    H36     109.322    1.870
HX2       H35    CMC    H37     109.322    1.870
HX2       H36    CMC    H37     109.322    1.870
HX2       CA     N      H       109.990    3.000
HX2       CA     N      H2      109.990    3.000
HX2       CA     N      H3      109.990    3.000
HX2       H      N      H2      109.032    3.000
HX2       H      N      H3      109.032    3.000
HX2       H2     N      H3      109.032    3.000
HX2       N      CA     C       109.258    1.500
HX2       N      CA     CB      110.314    2.210
HX2       N      CA     HA      108.387    1.580
HX2       C      CA     CB      110.876    3.000
HX2       C      CA     HA      108.774    1.790
HX2       CB     CA     HA      109.208    1.870
HX2       CA     C      O       117.148    1.600
HX2       CA     C      OXT     117.148    1.600
HX2       O      C      OXT     125.704    1.500
HX2       CA     CB     CG      113.420    2.400
HX2       CA     CB     HB2     108.559    1.500
HX2       CA     CB     HB3     108.559    1.500
HX2       CG     CB     HB2     108.800    1.500
HX2       CG     CB     HB3     108.800    1.500
HX2       HB3    CB     HB2     107.693    2.030
HX2       CD     CG     CB      113.967    3.000
HX2       CB     CG     HG2     108.780    1.500
HX2       CB     CG     HG3     108.780    1.500
HX2       CD     CG     HG2     108.753    1.500
HX2       CD     CG     HG3     108.753    1.500
HX2       HG3    CG     HG2     107.681    2.990

loop_
_chem_comp_tor.comp_id
_chem_comp_tor.id
_chem_comp_tor.atom_id_1
_chem_comp_tor.atom_id_2
_chem_comp_tor.atom_id_3
_chem_comp_tor.atom_id_4
_chem_comp_tor.value_angle
_chem_comp_tor.value_angle_esd
_chem_comp_tor.period
HX2       sp3_sp3_29 S1     CAC    CBC    H1       -60.000   10.000   3
HX2       sp2_sp2_81 C4A    CHB    C1B    C2B      180.000   20.000   2
HX2       sp2_sp2_84 NB     C1B    CHB    H8       180.000   20.000   2
HX2       const_45 NB     C1B    C2B    C3B        0.000    0.000   1
HX2       const_48 CHB    C1B    C2B    CMB        0.000    0.000   1
HX2       sp2_sp2_85 C1B    CHB    C4A    NA       180.000   20.000   2
HX2       sp2_sp2_88 C3A    C4A    CHB    H8       180.000   20.000   2
HX2       const_sp2_sp2_1 C1A    NA     C4A    C3A        0.000    0.000   1
HX2       const_sp2_sp2_4 CHB    C4A    NA     H9         0.000    0.000   1
HX2       const_89 NA     C4A    C3A    C2A        0.000    0.000   1
HX2       const_92 CHB    C4A    C3A    CMA        0.000    0.000   1
HX2       const_sp2_sp2_5 C4A    NA     C1A    C2A        0.000    0.000   1
HX2       const_sp2_sp2_8 CHA    C1A    NA     H9         0.000    0.000   1
HX2       const_sp2_sp2_9 NZ     CHA    C1A    C2A        0.000    0.000   1
HX2       const_12 NA     C1A    CHA    C4D        0.000    0.000   1
HX2       sp2_sp3_2 NA     C1A    C2A    CAA      120.000   10.000   6
HX2       const_13 C1A    CHA    NZ     CE         0.000    0.000   1
HX2       const_16 C4D    CHA    NZ     H10        0.000    0.000   1
```

| | | | | | | | | |
|---|---|---|---|---|---|---|---|---|
| HX2 | sp2_sp2_97 | C1A | CHA | C4D | C3D | 180.000 | 20.000 | 2 |
| HX2 | sp2_sp2_100 | NZ | CHA | C4D | ND | 180.000 | 20.000 | 2 |
| HX2 | const_17 | CHA | NZ | CE | CD | 0.000 | 0.000 | 1 |
| HX2 | const_20 | H10 | NZ | CE | H11 | 0.000 | 0.000 | 1 |
| HX2 | const_21 | NZ | CE | CD | C2A | 0.000 | 0.000 | 1 |
| HX2 | const_24 | H11 | CE | CD | CG | 0.000 | 0.000 | 1 |
| HX2 | sp2_sp3_17 | CAA | C2A | CD | CG | -60.000 | 10.000 | 6 |
| HX2 | sp2_sp3_26 | CE | CD | CG | CB | -90.000 | 10.000 | 6 |
| HX2 | sp3_sp3_40 | C1A | C2A | CAA | CBA | 180.000 | 10.000 | 3 |
| HX2 | sp2_sp3_12 | CAA | C2A | C3A | CMA | 60.000 | 10.000 | 6 |
| HX2 | sp3_sp3_37 | CBC | CAC | S1 | H5 | 180.000 | 10.000 | 3 |
| HX2 | sp2_sp3_32 | CBC | CAC | C3C | C4C | -90.000 | 10.000 | 6 |
| HX2 | sp3_sp3_49 | C2A | CAA | CBA | CGA | 180.000 | 10.000 | 3 |
| HX2 | sp2_sp3_38 | CAA | CBA | CGA | O2A | 120.000 | 10.000 | 6 |
| HX2 | sp2_sp2_93 | C2A | C3A | CMA | H18 | 180.000 | 20.000 | 2 |
| HX2 | sp2_sp2_96 | C4A | C3A | CMA | H17 | 180.000 | 20.000 | 2 |
| HX2 | const_25 | C1D | ND | C4D | C3D | 0.000 | 0.000 | 1 |
| HX2 | const_109 | ND | C4D | C3D | C2D | 0.000 | 0.000 | 1 |
| HX2 | const_112 | CHA | C4D | C3D | CAD | 0.000 | 0.000 | 1 |
| HX2 | const_27 | C4D | ND | C1D | C2D | 0.000 | 0.000 | 1 |
| HX2 | sp2_sp2_105 | C4C | CHD | C1D | C2D | 180.000 | 20.000 | 2 |
| HX2 | sp2_sp2_108 | ND | C1D | CHD | H19 | 180.000 | 20.000 | 2 |
| HX2 | const_29 | ND | C1D | C2D | C3D | 0.000 | 0.000 | 1 |
| HX2 | const_32 | CHD | C1D | C2D | CMD | 0.000 | 0.000 | 1 |
| HX2 | sp2_sp3_43 | C1D | C2D | CMD | H20 | 150.000 | 10.000 | 6 |
| HX2 | const_33 | C4D | C3D | C2D | C1D | 0.000 | 0.000 | 1 |
| HX2 | const_36 | CMD | C2D | C3D | CAD | 0.000 | 0.000 | 1 |
| HX2 | sp2_sp3_50 | C4D | C3D | CAD | CBD | -90.000 | 10.000 | 6 |
| HX2 | sp3_sp3_58 | C3D | CAD | CBD | CGD | 180.000 | 10.000 | 3 |
| HX2 | sp2_sp3_56 | CAD | CBD | CGD | O2D | 120.000 | 10.000 | 6 |
| HX2 | sp2_sp3_61 | C1B | C2B | CMB | H27 | 150.000 | 10.000 | 6 |
| HX2 | const_41 | C4B | C3B | C2B | C1B | 0.000 | 0.000 | 1 |
| HX2 | const_44 | CMB | C2B | C3B | CAB | 0.000 | 0.000 | 1 |
| HX2 | const_51 | NC | C4C | C3C | C2C | 0.000 | 0.000 | 1 |
| HX2 | const_54 | CAC | C3C | C4C | CHD | 0.000 | 0.000 | 1 |
| HX2 | const_67 | C4C | C3C | C2C | C1C | 0.000 | 0.000 | 1 |
| HX2 | const_70 | CAC | C3C | C2C | CMC | 0.000 | 0.000 | 1 |
| HX2 | sp2_sp3_68 | C4B | C3B | CAB | CBB | -90.000 | 10.000 | 6 |
| HX2 | sp3_sp3_70 | S2 | CAB | CBB | H311 | 60.000 | 10.000 | 3 |
| HX2 | sp3_sp3_76 | CBB | CAB | S2 | H34 | 180.000 | 10.000 | 3 |
| HX2 | sp2_sp3_73 | C3C | C2C | CMC | H35 | 150.000 | 10.000 | 6 |
| HX2 | sp3_sp3_4 | C | CA | N | H | 60.000 | 10.000 | 3 |
| HX2 | sp2_sp3_19 | N | CA | C | O | 0.000 | 10.000 | 6 |
| HX2 | sp3_sp3_13 | N | CA | CB | CG | 60.000 | 10.000 | 3 |
| HX2 | sp2_sp2_101 | C3C | C4C | CHD | C1D | 180.000 | 20.000 | 2 |
| HX2 | sp2_sp2_104 | NC | C4C | CHD | H19 | 180.000 | 20.000 | 2 |
| HX2 | const_55 | C3C | C4C | NC | C1C | 0.000 | 0.000 | 1 |
| HX2 | const_58 | CHD | C4C | NC | H6 | 0.000 | 0.000 | 1 |
| HX2 | sp3_sp3_19 | CD | CG | CB | CA | 180.000 | 10.000 | 3 |
| HX2 | const_59 | C4C | NC | C1C | C2C | 0.000 | 0.000 | 1 |
| HX2 | const_62 | CHC | C1C | NC | H6 | 0.000 | 0.000 | 1 |
| HX2 | const_63 | C3C | C2C | C1C | NC | 0.000 | 0.000 | 1 |
| HX2 | const_66 | CHC | C1C | C2C | CMC | 0.000 | 0.000 | 1 |
| HX2 | sp2_sp2_71 | NC | C1C | CHC | C4B | 180.000 | 20.000 | 2 |
| HX2 | sp2_sp2_74 | C2C | C1C | CHC | H7 | 180.000 | 20.000 | 2 |
| HX2 | sp2_sp2_75 | C1C | CHC | C4B | C3B | 180.000 | 20.000 | 2 |
| HX2 | sp2_sp2_78 | NB | C4B | CHC | H7 | 180.000 | 20.000 | 2 |
| HX2 | const_37 | NB | C4B | C3B | C2B | 0.000 | 0.000 | 1 |
| HX2 | const_40 | CHC | C4B | C3B | CAB | 0.000 | 0.000 | 1 |
| HX2 | const_79 | C1B | NB | C4B | C3B | 0.000 | 0.000 | 1 |
| HX2 | const_49 | C4B | NB | C1B | C2B | 0.000 | 0.000 | 1 |

```
loop_
_chem_comp_chir.comp_id
_chem_comp_chir.id
_chem_comp_chir.atom_id_centre
_chem_comp_chir.atom_id_1
_chem_comp_chir.atom_id_2
_chem_comp_chir.atom_id_3
_chem_comp_chir.volume_sign
HX2      chir_1   CA     N      C      CB        positiv
HX2      chir_1   CAC    CBC    S1     C3C       positiv
HX2      chir_2   C2A    C1A    C3A    CD        positiv
HX2      chir_3   CAB    C3B    CBB    S2        positiv

loop_
_chem_comp_plane_atom.comp_id
_chem_comp_plane_atom.plane_id
_chem_comp_plane_atom.atom_id
_chem_comp_plane_atom.dist_esd
HX2      plan-1    CAC       0.020
HX2      plan-1    C3C       0.020
HX2      plan-1    C4C       0.020
HX2      plan-1    NC        0.020
HX2      plan-1    C1C       0.020
HX2      plan-1    CHC       0.020
HX2      plan-1    CHD       0.020
HX2      plan-1    C2C       0.020
HX2      plan-1    CMC       0.020
HX2      plan-1    H6        0.020
HX2      plan-2    C1C       0.020
HX2      plan-2    CHC       0.020
HX2      plan-2    C4B       0.020
HX2      plan-2    H7        0.020
HX2      plan-3    CHC       0.020
HX2      plan-3    C4B       0.020
HX2      plan-3    NB        0.020
HX2      plan-3    C3B       0.020
HX2      plan-4    NB        0.020
HX2      plan-4    C1B       0.020
HX2      plan-4    CHB       0.020
HX2      plan-4    C2B       0.020
HX2      plan-5    C1B       0.020
HX2      plan-5    CHB       0.020
HX2      plan-5    C4A       0.020
HX2      plan-5    H8        0.020
HX2      plan-6    CHB       0.020
HX2      plan-6    C4A       0.020
HX2      plan-6    NA        0.020
HX2      plan-6    C3A       0.020
HX2      plan-7    C4A       0.020
HX2      plan-7    NA        0.020
HX2      plan-7    C1A       0.020
HX2      plan-7    H9        0.020
HX2      plan-8    NA        0.020
HX2      plan-8    C1A       0.020
HX2      plan-8    CHA       0.020
HX2      plan-8    C2A       0.020
HX2      plan-9    C1A       0.020
HX2      plan-9    CHA       0.020
HX2      plan-9    NZ        0.020
HX2      plan-9    C4D       0.020
```

| | | | |
|---|---|---|---|
| HX2 | plan-10 | CHA | 0.020 |
| HX2 | plan-10 | NZ | 0.020 |
| HX2 | plan-10 | CE | 0.020 |
| HX2 | plan-10 | H10 | 0.020 |
| HX2 | plan-11 | NZ | 0.020 |
| HX2 | plan-11 | CE | 0.020 |
| HX2 | plan-11 | CD | 0.020 |
| HX2 | plan-11 | H11 | 0.020 |
| HX2 | plan-12 | CE | 0.020 |
| HX2 | plan-12 | CD | 0.020 |
| HX2 | plan-12 | C2A | 0.020 |
| HX2 | plan-12 | CG | 0.020 |
| HX2 | plan-13 | CBA | 0.020 |
| HX2 | plan-13 | CGA | 0.020 |
| HX2 | plan-13 | O2A | 0.020 |
| HX2 | plan-13 | O1A | 0.020 |
| HX2 | plan-14 | C4A | 0.020 |
| HX2 | plan-14 | C2A | 0.020 |
| HX2 | plan-14 | C3A | 0.020 |
| HX2 | plan-14 | CMA | 0.020 |
| HX2 | plan-15 | C3A | 0.020 |
| HX2 | plan-15 | CMA | 0.020 |
| HX2 | plan-15 | H17 | 0.020 |
| HX2 | plan-15 | H18 | 0.020 |
| HX2 | plan-16 | CHA | 0.020 |
| HX2 | plan-16 | C4D | 0.020 |
| HX2 | plan-16 | ND | 0.020 |
| HX2 | plan-16 | C3D | 0.020 |
| HX2 | plan-17 | ND | 0.020 |
| HX2 | plan-17 | C1D | 0.020 |
| HX2 | plan-17 | CHD | 0.020 |
| HX2 | plan-17 | C2D | 0.020 |
| HX2 | plan-18 | C4C | 0.020 |
| HX2 | plan-18 | C1D | 0.020 |
| HX2 | plan-18 | CHD | 0.020 |
| HX2 | plan-18 | H19 | 0.020 |
| HX2 | plan-19 | C1D | 0.020 |
| HX2 | plan-19 | C2D | 0.020 |
| HX2 | plan-19 | CMD | 0.020 |
| HX2 | plan-19 | C3D | 0.020 |
| HX2 | plan-20 | C4D | 0.020 |
| HX2 | plan-20 | C2D | 0.020 |
| HX2 | plan-20 | C3D | 0.020 |
| HX2 | plan-20 | CAD | 0.020 |
| HX2 | plan-21 | CBD | 0.020 |
| HX2 | plan-21 | CGD | 0.020 |
| HX2 | plan-21 | O1D | 0.020 |
| HX2 | plan-21 | O2D | 0.020 |
| HX2 | plan-22 | C1B | 0.020 |
| HX2 | plan-22 | C2B | 0.020 |
| HX2 | plan-22 | CMB | 0.020 |
| HX2 | plan-22 | C3B | 0.020 |
| HX2 | plan-23 | C4B | 0.020 |
| HX2 | plan-23 | C2B | 0.020 |
| HX2 | plan-23 | C3B | 0.020 |
| HX2 | plan-23 | CAB | 0.020 |
| HX2 | plan-24 | CA | 0.020 |
| HX2 | plan-24 | C | 0.020 |
| HX2 | plan-24 | O | 0.020 |
| HX2 | plan-24 | OXT | 0.020 |

```
data_mod_HX2mod1

loop_
_chem_mod_atom.mod_id
_chem_mod_atom.function
_chem_mod_atom.atom_id
_chem_mod_atom.new_atom_id
_chem_mod_atom.new_type_symbol
_chem_mod_atom.new_type_energy
_chem_mod_atom.new_charge
HX2mod1  delete H5     .      .    .        .
HX2mod1  delete S1     .      .    .        .

loop_
_chem_mod_bond.mod_id
_chem_mod_bond.function
_chem_mod_bond.atom_id_1
_chem_mod_bond.atom_id_2
_chem_mod_bond.new_type
_chem_mod_bond.new_aromatic
_chem_mod_bond.new_value_dist
_chem_mod_bond.new_value_dist_esd
HX2mod1  change CBC    CAC       .        .    1.525    0.015
HX2mod1  delete CAC    S1        single   n        .        .
HX2mod1  delete S1     H5        single   n        .        .

loop_
_chem_mod_angle.mod_id
_chem_mod_angle.function
_chem_mod_angle.atom_id_1
_chem_mod_angle.atom_id_2
_chem_mod_angle.atom_id_3
_chem_mod_angle.new_value_angle
_chem_mod_angle.new_value_angle_esd
HX2mod1  delete CBC    CAC    S1           .        .
HX2mod1  delete S1     CAC    C3C          .        .
HX2mod1  delete S1     CAC    H4           .        .
HX2mod1  delete CAC    S1     H5           .        .
HX2mod1  change CAC    C3C    C2C     127.355    1.500

loop_
_chem_mod_chir.mod_id
_chem_mod_chir.function
_chem_mod_chir.id
_chem_mod_chir.atom_id_centre
_chem_mod_chir.atom_id_1
_chem_mod_chir.atom_id_2
_chem_mod_chir.atom_id_3
_chem_mod_chir.new_volume_sign
HX2mod1  delete chir_1   CAC    CBC    S1     C3C       positiv

data_mod_CYSmod1

loop_
_chem_mod_atom.mod_id
_chem_mod_atom.function
_chem_mod_atom.atom_id
_chem_mod_atom.new_atom_id
_chem_mod_atom.new_type_symbol
_chem_mod_atom.new_type_energy
```

```
_chem_mod_atom.new_charge
CYSmod1  change SG     .      .    S        .
CYSmod1  add    .      HSG1   H    HSH1     0

loop_
_chem_mod_bond.mod_id
_chem_mod_bond.function
_chem_mod_bond.atom_id_1
_chem_mod_bond.atom_id_2
_chem_mod_bond.new_type
_chem_mod_bond.new_aromatic
_chem_mod_bond.new_value_dist
_chem_mod_bond.new_value_dist_esd
CYSmod1  change CB     SG        .         .    1.778    0.010
CYSmod1  add    SG     HSG1      single    n    1.225    0.020

loop_
_chem_mod_angle.mod_id
_chem_mod_angle.function
_chem_mod_angle.atom_id_1
_chem_mod_angle.atom_id_2
_chem_mod_angle.atom_id_3
_chem_mod_angle.new_value_angle
_chem_mod_angle.new_value_angle_esd
CYSmod1  change N      CA     CB      112.343    1.500
CYSmod1  change CB     CA     HA      106.687    1.500
CYSmod1  change CA     CB     SG      110.778    3.000
CYSmod1  change CA     CB     HB3     108.012    1.500
CYSmod1  change CA     CB     HB2     108.012    1.500
CYSmod1  change CB     SG     HG      109.471    3.000
CYSmod1  add    CB     SG     HSG1    109.471    3.000
CYSmod1  add    HG     SG     HSG1    109.471    3.000

data_mod_CYSmod2

loop_
_chem_mod_atom.mod_id
_chem_mod_atom.function
_chem_mod_atom.atom_id
_chem_mod_atom.new_atom_id
_chem_mod_atom.new_type_symbol
_chem_mod_atom.new_type_energy
_chem_mod_atom.new_charge
CYSmod2  change SG     .      .    S        .
CYSmod2  add    .      HSG1   H    HSH1     0

loop_
_chem_mod_bond.mod_id
_chem_mod_bond.function
_chem_mod_bond.atom_id_1
_chem_mod_bond.atom_id_2
_chem_mod_bond.new_type
_chem_mod_bond.new_aromatic
_chem_mod_bond.new_value_dist
_chem_mod_bond.new_value_dist_esd
CYSmod2  change CB     SG        .         .    1.778    0.010
CYSmod2  add    SG     HSG1      single    n    1.225    0.020

loop_
_chem_mod_angle.mod_id
```

```
_chem_mod_angle.function
_chem_mod_angle.atom_id_1
_chem_mod_angle.atom_id_2
_chem_mod_angle.atom_id_3
_chem_mod_angle.new_value_angle
_chem_mod_angle.new_value_angle_esd
CYSmod2  change N      CA     CB      112.343    1.500
CYSmod2  change CB     CA     HA      106.687    1.500
CYSmod2  change CA     CB     SG      110.778    3.000
CYSmod2  change CA     CB     HB3     108.012    1.500
CYSmod2  change CA     CB     HB2     108.012    1.500
CYSmod2  change CB     SG     HG      109.471    3.000
CYSmod2  add    CB     SG     HSG1    109.471    3.000
CYSmod2  add    HG     SG     HSG1    109.471    3.000

data_mod_HX2mod2

loop_
_chem_mod_atom.mod_id
_chem_mod_atom.function
_chem_mod_atom.atom_id
_chem_mod_atom.new_atom_id
_chem_mod_atom.new_type_symbol
_chem_mod_atom.new_type_energy
_chem_mod_atom.new_charge
HX2mod2  delete H34    .      .    .        .
HX2mod2  delete S2     .      .    .        .

loop_
_chem_mod_bond.mod_id
_chem_mod_bond.function
_chem_mod_bond.atom_id_1
_chem_mod_bond.atom_id_2
_chem_mod_bond.new_type
_chem_mod_bond.new_aromatic
_chem_mod_bond.new_value_dist
_chem_mod_bond.new_value_dist_esd
HX2mod2  change CAB    CBB       .        .    1.525    0.015
HX2mod2  delete CAB    S2        single   n        .        .
HX2mod2  delete S2     H34       single   n        .        .

loop_
_chem_mod_angle.mod_id
_chem_mod_angle.function
_chem_mod_angle.atom_id_1
_chem_mod_angle.atom_id_2
_chem_mod_angle.atom_id_3
_chem_mod_angle.new_value_angle
_chem_mod_angle.new_value_angle_esd
HX2mod2  change C4B    C3B    CAB     124.022    3.000
HX2mod2  change C2B    C3B    CAB     129.176    1.500
HX2mod2  delete C3B    CAB    S2            .        .
HX2mod2  delete CBB    CAB    S2            .        .
HX2mod2  delete S2     CAB    H30           .        .
HX2mod2  delete CAB    S2     H34           .        .

loop_
_chem_mod_chir.mod_id
_chem_mod_chir.function
_chem_mod_chir.id
```

```
_chem_mod_chir.atom_id_centre
_chem_mod_chir.atom_id_1
_chem_mod_chir.atom_id_2
_chem_mod_chir.atom_id_3
_chem_mod_chir.new_volume_sign
HX2mod2  delete chir_3   CAB    C3B    CBB    S2        positiv

data_link_HX2-CYS

loop_
_chem_link_bond.link_id
_chem_link_bond.atom_1_comp_id
_chem_link_bond.atom_id_1
_chem_link_bond.atom_2_comp_id
_chem_link_bond.atom_id_2
_chem_link_bond.type
_chem_link_bond.aromatic
_chem_link_bond.value_dist
_chem_link_bond.value_dist_esd
HX2-CYS  1 CAC    2 SG        single   n    1.811    0.020

loop_
_chem_link_angle.link_id
_chem_link_angle.atom_1_comp_id
_chem_link_angle.atom_id_1
_chem_link_angle.atom_2_comp_id
_chem_link_angle.atom_id_2
_chem_link_angle.atom_3_comp_id
_chem_link_angle.atom_id_3
_chem_link_angle.value_angle
_chem_link_angle.value_angle_esd
HX2-CYS  1 CBC    1 CAC    2 SG      107.734    3.000
HX2-CYS  1 C3C    1 CAC    2 SG      112.507    3.000
HX2-CYS  1 H4     1 CAC    2 SG      107.260    1.500
HX2-CYS  1 CAC    2 SG     2 CB      104.305    3.000
HX2-CYS  1 CAC    2 SG     2 HG      109.471    3.000
HX2-CYS  1 CAC    2 SG     2 HSG1    109.471    3.000

loop_
_chem_link_chir.link_id
_chem_link_chir.id
_chem_link_chir.atom_centre_comp_id
_chem_link_chir.atom_id_centre
_chem_link_chir.atom_1_comp_id
_chem_link_chir.atom_id_1
_chem_link_chir.atom_2_comp_id
_chem_link_chir.atom_id_2
_chem_link_chir.atom_3_comp_id
_chem_link_chir.atom_id_3
_chem_link_chir.volume_sign
HX2-CYS  chir_5   1 CAC    1 CBC    2 SG     1 C3C       positiv

data_link_CYS-HX2

loop_
_chem_link_bond.link_id
_chem_link_bond.atom_1_comp_id
_chem_link_bond.atom_id_1
_chem_link_bond.atom_2_comp_id
_chem_link_bond.atom_id_2
```

```
_chem_link_bond.type
_chem_link_bond.aromatic
_chem_link_bond.value_dist
_chem_link_bond.value_dist_esd
CYS-HX2  1 SG     2 CAB       single   n    1.811    0.020

loop_
_chem_link_angle.link_id
_chem_link_angle.atom_1_comp_id
_chem_link_angle.atom_id_1
_chem_link_angle.atom_2_comp_id
_chem_link_angle.atom_id_2
_chem_link_angle.atom_3_comp_id
_chem_link_angle.atom_id_3
_chem_link_angle.value_angle
_chem_link_angle.value_angle_esd
CYS-HX2  1 SG     2 CAB    2 C3B     111.378    3.000
CYS-HX2  1 SG     2 CAB    2 CBB     107.734    3.000
CYS-HX2  1 SG     2 CAB    2 H30     107.260    1.500
CYS-HX2  1 CB     1 SG     2 CAB     104.305    3.000
CYS-HX2  1 HG     1 SG     2 CAB     109.471    3.000
CYS-HX2  1 HSG1   1 SG     2 CAB     109.471    3.000

loop_
_chem_link_chir.link_id
_chem_link_chir.id
_chem_link_chir.atom_centre_comp_id
_chem_link_chir.atom_id_centre
_chem_link_chir.atom_1_comp_id
_chem_link_chir.atom_id_1
_chem_link_chir.atom_2_comp_id
_chem_link_chir.atom_id_2
_chem_link_chir.atom_3_comp_id
_chem_link_chir.atom_id_3
_chem_link_chir.volume_sign
CYS-HX2  chir_5   2 CAB    1 SG     2 C3B    2 CBB       positiv
```

### RADDOSE input

Used for calculating the dose of the ferrous structure from the first dataset. Dose multiplied by 50/20 for the second dataset.

```
##############################################################################
#                              Crystal Block                                 #
##############################################################################

Crystal

Type Cylinder
       # Crystal shape can be Cuboid or Spherical

Dimensions 25 100
       # Diameter and Height

PixelsPerMicron 1
       # This defines the coarseness of the simulation
       # (i.e. how many voxels the crystal is divided into.)
       # Preferably set as high as possible, however for a higher
       # value the simulation will take longer to complete.
       # Recommended to try increasing between 0.5 and 5 and ensure
```

```
        # the reported dose value converges as PixelsPerMicron increases.
        # As a rule of thumb, this needs to be at least 10x the beam
        # FWHM for a Gaussian beam.
        # e.g. 20µm FWHM beam -> 2µm voxels -> 0.5 voxels/µm

# NOTE: Use AngleP/AngleL if your crystal is not face-on to the beam.
# See RD3D user guide for more details
ANGLEP 23
# Also need to specify the crystal composition below (Example case for insulin given):
AbsCoefCalc  RD3D
        # Absorption Coefficients calculated
        # using RADDOSE-3D (Zeldin et al. 2013).

UnitCell   46.380   80.441   84.780
        # unit cell size: a, b, c with alpha, beta and gamma angles default to 90°

NumMonomers  8
        # number of monomers in unit cell

NumResidues  144
        # number of residues per monomer

ProteinHeavyAtoms Fe 1 S 6
        # heavy atoms added to protein part of the
        # monomer, i.e. S, coordinated metals, Se in Se-Met

SolventHeavyConc S 1910 Na 2127
        # concentration of elements in the solvent
        # in mmol/l. Oxygen and lighter elements
        # should not be specified

SolventFraction 0.5247
        # fraction of the unit cell occupied by solvent

##############################################################################
#                                  Beam Block                                #
##############################################################################

Beam

Type Gaussian
        # beam profile can be Gaussian or TopHat
Flux 0.3e12
        # in photons per second (2e12 = 2 * 10^12)
FWHM 50 50
        # in µm, horizontal by vertical for a Gaussian beam
Energy 20
        # photon energy in keV

Collimation Rectangular 150 150
        # Horizontal/Vertical collimation of the beam
        # For 'uncollimated' Gaussians, 3xFWHM recommended


##############################################################################
#                                  Wedge Block                               #
##############################################################################

Wedge -90 90
        # Start and End rotational angle of the crystal with Start < End

ExposureTime 18
        # Total time for entire angular range (seconds)
```
